# Terazosin drives sex-dependent adrenergic–bioenergetic reprogramming to restore network function in Alzheimer’s disease

**DOI:** 10.64898/2026.04.02.716175

**Authors:** Swastik G. Pattanashetty, Peter A. Serrano, Patricia Rockwell, Lei Xie, Maria E. Figueiredo-Pereira

**Author notes:** Correspondence to: Maria Figueiredo-Pereira, Ph.D., 695 Park Avenue, New York, NY 10065.

## Abstract

Alzheimer’s disease (AD) has long been defined by amyloid-β plaques and hyperphosphorylated tau, yet disease-modifying therapies remain critically limited. Growing evidence reframes AD as a system-level failure driven by early dysregulation of synaptic, metabolic, and neuroimmune pathways, preceding overt protein aggregation and originating in selectively vulnerable circuits, including the locus coeruleus (LC)-hippocampal noradrenergic axis. This complexity underscores the need for therapeutic strategies that engage the disease at a network level, early in its trajectory.

To this end, using a machine learning–based systems pharmacology framework for drug repurposing applied to human AD transcriptomic datasets, we identified terazosin (TZ) as a candidate predicted to reverse AD-associated molecular signatures. TZ is an FDA-approved α₁-adrenergic receptor antagonist and phosphoglycerate kinase-1 activator. It was administered chronically via the diet (0.5 mg/kg bw/day) to male and female TgF344-AD rats and wild-type littermates from 5 to 11 months of age, preceding overt pathology.

Bulk hippocampal RNA sequencing revealed sex-specific transcriptional remodeling in transgenic rats, strongly conserved with human AD datasets. Male TgF344-AD rats exhibited suppression of synaptic and transcriptional maintenance pathways with concurrent activation of metabolic, proteostatic, extracellular matrix, and vascular stress responses; females showed suppression of survival and vascular structural signaling alongside heightened DAM-like immune activation, amyloid-associated stress, and cell death programs. TZ reversed these signatures in a sex-dependent manner: males showed enhanced immune surveillance and reduced proteostasis burden, while females showed reinforcement of synaptic, survival, and metabolic pathways. TgF344-AD rats displayed selective LC-derived hippocampal noradrenergic axonopathy without global neuronal loss. TZ preserved fiber integrity preferentially in females and partially reversed LC vulnerability–associated transcriptional signatures in both sexes. TZ also reduced amyloid-β plaque burden in both sexes, attenuated hyperphosphorylated tau exclusively in females, and induced microglial morphological shifts in males. Finally, TZ restored wild-type spatial learning in transgenic animals, with females appearing to derive the greater cognitive benefit.

Together, these findings demonstrate that TZ induced systems-level reprogramming of AD-relevant molecular pathways and preserved vulnerable noradrenergic circuitry in a sex-dependent manner. Moreover, TZ rescued spatial cognition in transgenic rats, with cognitive gains seemingly more pronounced in females. These results support adrenergic–bioenergetic modulation as a translational strategy for early-stage AD and underscore the necessity of sex as a biological variable in disease-modifying treatment development.

## INTRODUCTION

Alzheimer’s disease (AD) is a progressive neurodegenerative disorder characterized by cognitive decline, neuronal loss and the accumulation of amyloid-β (Aβ) plaques and hyperphosphorylated tau (pTau) tangles^1–5^. Growing evidence indicates that early disruption of noradrenergic, synaptic and immune systems plays a critical role in disease onset and progression^6–13^. Among the earliest and most vulnerable neuronal populations are locus coeruleus (LC)–derived noradrenergic neurons^9,14^. The LC provides widespread projections throughout the brain including the hippocampus and prefrontal cortex and regulates arousal, attention, learning and memory^6,7,15^. LC neurodegeneration occurs decades before extensive cortical pathology in AD^6,8,16^ disrupting neuromodulatory balance, impairing synaptic plasticity and reducing immune surveillance, thereby exacerbating amyloid, tau and inflammatory cascades^10,17–19^. Compounding these pathological processes, AD progression is further driven by a profound dysregulation of the transcriptomic landscape, wherein early shifts in transcriptional programs render vulnerable neuronal populations including LC-derived noradrenergic neurons susceptible to degeneration prior to overt clinical manifestation^9^. Preservation of LC–hippocampal integrity and restoration of homeostatic transcriptional programs have consequently emerged as promising strategies to delay and potentially reverse AD progression^20–26^.

AD also manifests with striking sex differences in both incidence and progression^27,28^. Women exhibit higher prevalence and faster cognitive decline than men^29–32^, with emerging evidence of heightened vulnerability of noradrenergic and synaptic pathways in females, including earlier LC fiber degeneration^33,34^. Yet sex has historically been underrepresented as a biological variable in preclinical and clinical research. Dissecting sex-dependent molecular mechanisms and identifying interventions capable of addressing them remain critical unmet needs in AD therapeutics^13,32,33^.

Despite decades of research targeting Aβ and tau, nearly 99% of AD clinical trials have failed^35^, highlighting the need for strategies beyond classical pathological targets. Drug repurposing offers a cost-effective alternative, leveraging compounds with established safety profiles to target disease-relevant mechanisms^36^.

Here, we applied a machine learning–based systems pharmacology algorithm integrating structure-based off-target prediction with drug-induced gene expression modeling pipeline described in our previous publication^37^ and in^38^. Transcriptomic datasets from the AMP-AD consortium defined the molecular signature of AD through differential gene expression between AD patients and healthy controls. Compounds were prioritized if predicted to reverse the AD transcriptome, upregulating suppressed and downregulating aberrantly elevated transcripts. Using this framework, terazosin (TZ) emerged as one of the top candidates^38^.

Independent evidence supports this prediction. TZ is an FDA-approved α₁-adrenergic receptor antagonist used for hypertension and benign prostatic hyperplasia^39^ that also directly binds and activates phosphoglycerate kinase-1 (PGK1), enhancing glycolysis and ATP production^40^. By augmenting cellular bioenergetics, TZ promotes mitochondrial resilience, reduces oxidative stress and apoptosis ^40,41^ and attenuates neuronal death in models of Parkinson’s disease, ALS and ischemic injury^41,42^. In an AD-relevant context, α₁-adrenergic receptor inhibition via TZ ameliorated pathology and behavioral deficits in APPswe/PS1 mice^43^ and population-based analysis linked TZ and related α₁-adrenergic receptor antagonists to reduced dementia with Lewy bodies risk in men^44^.

Whether TZ protects the LC–hippocampal noradrenergic network, which undergoes early retrograde degeneration in AD, remains unknown, as does its impact on hippocampal molecular pathways in a sex-dependent context, particularly given evidence that AD transcriptomic signatures differ between sexes^27,29,34^.

To address this, we investigated AD progression in the TgF344-AD rat model, which recapitulates key pathological, molecular and behavioral features of human AD in an age-dependent manner^45^. Using hippocampal RNA sequencing, we characterized AD-associated transcriptomic pathways and determined how chronic TZ treatment modulates them in male and female TgF344-AD rats. Integrating these findings with human AD hippocampal and LC transcriptomic datasets enabled identification of conserved molecular pathways and TZ-mediated transcriptome reversal, with particular focus on noradrenergic, synaptic and immune pathways. We further examined whether TZ preserves hippocampal noradrenergic fibers, reduces Aβ burden, attenuates tau hyperphosphorylation, modulates microglial activation and improves hippocampal-dependent spatial memory in a sex dependent manner.

Collectively, our findings identify TZ as a promising repurposing candidate in AD, demonstrating that PGK1-mediated bioenergetic enhancement and adrenergic modulation restore AD-associated molecular networks, preserve noradrenergic projections and reduce hallmark neuropathology in a sex-dependent manner, underscoring biological sex as a critical determinant of therapeutic efficacy and the need for precision approaches in AD treatment^32,33^.

## RESULTS

### Sex-specific hippocampal transcriptional dysregulation in TgF344-AD rats

To characterize the AD-associated transcriptome in the TgF344-AD rat model, we performed bulk RNA sequencing of the right dorsal hippocampus from non-treated 11-month-old male and female transgenic rats (TGNT) and age-matched wild-type littermates (WTNT). Applying thresholds of P < 0.05 and |log₂ fold change| > 1.0, we identified 574 differentially expressed genes (DEGs) in males and 383 DEGs in females (Supplemental Tables 1 and 2). Unsupervised hierarchical clustering revealed clear genotype-dependent segregation in both sexes (Fig. 1A, B), with coordinated gene modules displaying opposing expression patterns between WTNT and TGNT groups, consistent with hippocampal transcriptional remodeling driven by overexpression of the AD-linked transgenes APPswe (K670M/N671L) and PSEN1ΔE9.

**Figure 1.**
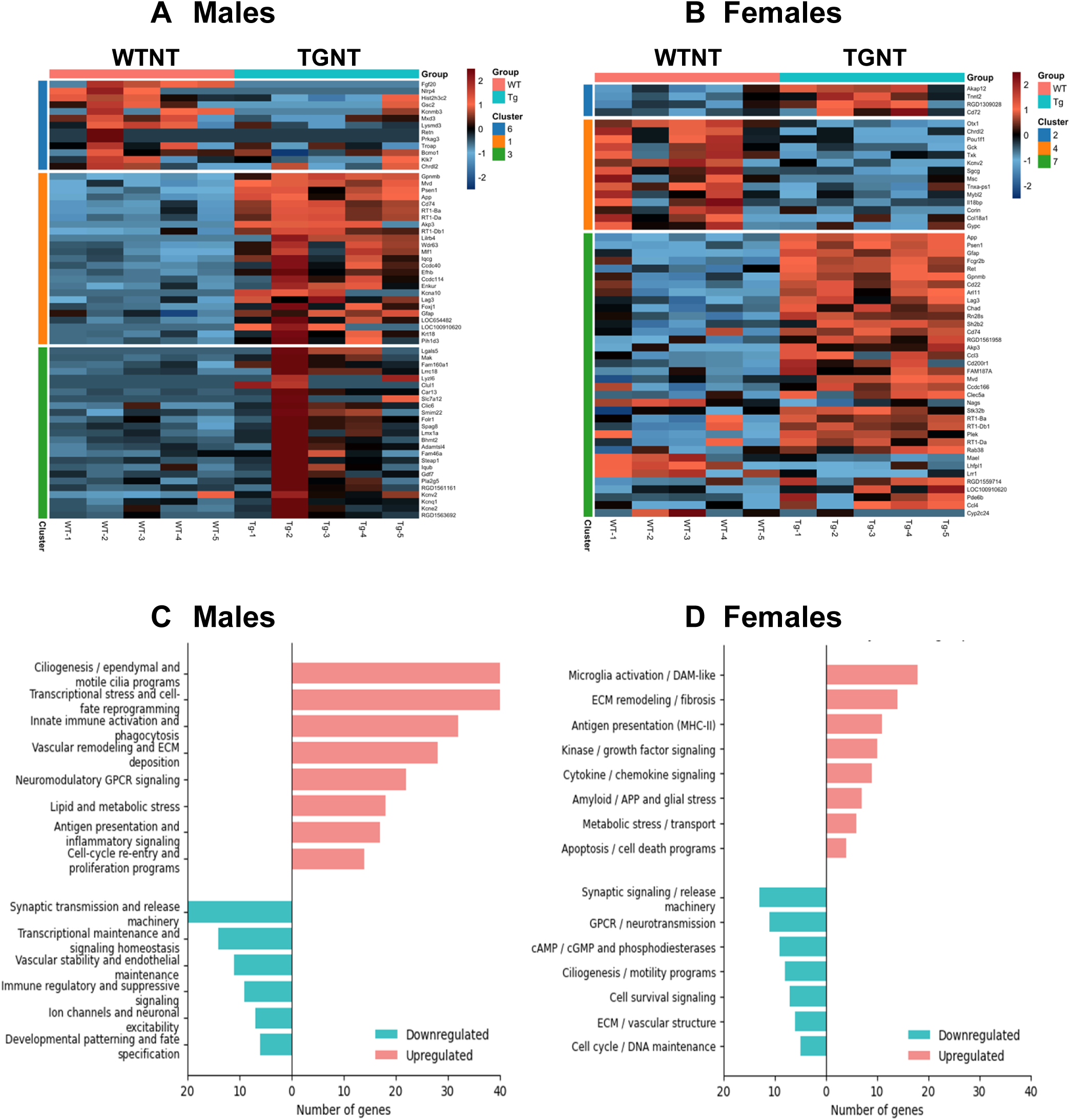
Sex-specific hippocampal transcriptional dysregulation in TgF344-AD rats. **(A, B)** Unsupervised hierarchical clustering heatmaps of differentially expressed genes (DEGs; *P* < 0.05, |log₂FC| > 1.5) from bulk hippocampal RNA sequencing of 11-month-old non-treated transgenic (TGNT) and wild-type (WTNT) rats in males **(A**, n = 574 DEGs) and females (**B**, n = 383 DEGs), with 5 rats per each of the four groups. Gene modules display reciprocal expression patterns between WTNT and TGNT groups, reflecting genotype-dependent transcriptional remodeling. **(C, D)** Pathway enrichment analysis of upregulated (red) and downregulated (blue) DEGs in male **(C)** and female **(D)** TGNT rats relative to WTNT controls. Pathways are ranked by enrichment score. Convergent suppression of synaptic transmission and activation of neuroinflammatory signaling are observed across sexes. Males exhibit prominent upregulation of ciliogenesis, vascular remodeling, innate immune and inflammatory programs with suppression of synaptic and neuronal signaling pathways. Females display enrichment of microglial/immune, ECM remodeling, and kinase signaling pathways, alongside downregulation of synaptic transmission and GPCR-related signaling modules. Gene lists are provided in Supplementary Tables 1–4. WTNT, wild-type not treated; TGNT, transgenic not treated.

Pathway enrichment analysis uncovered extensive yet sexually divergent transcriptional reorganization in TgF344 AD rats. In males (Fig. 1C), upregulated pathways encompassed ciliogenesis and motile cilia programs, innate immune activation and phagocytosis, transcriptional and metabolic stress responses, vascular remodeling with ECM deposition, antigen presentation, cytokine-driven inflammatory signaling, and cell-cycle re-entry. Conversely, downregulated pathways were dominated by synaptic transmission and vesicle release machinery, neuronal signaling homeostasis, endothelial and vascular maintenance, immune regulatory programs, and ion channel activity governing neuronal excitability. In females (Fig. 1D), upregulated pathways were characterized by microglial activation and disease-associated microglia (DAM)-like signatures, ECM remodeling and fibrosis, MHC-II–mediated antigen presentation, cytokine and kinase signaling, amyloid/APP-associated stress responses, metabolic dysregulation, and pro-apoptotic programs. Downregulated pathways paralleled males in suppressing synaptic signaling and vesicle release machinery, but additionally included GPCR-mediated neurotransmission, cAMP/cGMP signaling, ciliogenesis-related programs, and ECM/vascular structural integrity.

Despite these sex-specific signatures, two molecular features converged across sexes: suppression of synaptic transmission and activation of inflammatory signaling (Fig. 1C, D). Nevertheless, the molecular context diverged markedly: males exhibited stronger enrichment of vascular remodeling, ciliogenesis, and lipid/metabolic stress programs, whereas females displayed greater activation of microglial, antigen presentation, and apoptotic pathways. Together, these findings demonstrate that synaptic dysfunction and neuroinflammation represent convergent pathological outcomes in the TgF344-AD hippocampus, yet arise from partially distinct, sex-specific upstream molecular programs, implicating divergent cellular mechanisms in driving sex-differential vulnerability to AD pathology. Genes contributing to these pathways are listed in Supplementary Tables 3 and 4.

### Human core AD transcriptional signatures are conserved in the TgF344-AD rat hippocampus

To establish the translational validity of these transcriptional signatures, we cross-referenced TGNT hippocampal DEGs against the AMP-AD AGORA database ^46^ an open repository of over 900 candidate AD therapeutic targets derived from large-scale human multi-omic studies. The degree of overlap was striking: 497 of 574 male TGNT DEGs (73.1%; Fig. 2A) and 360 of 383 female TGNT DEGs (66.2%; Fig. 2B) mapped directly to human AD-associated genes, with only a minor fraction remaining species-specific. Conserved gene sets are provided in Supplementary Tables 5 and 6.

**Figure 2.**
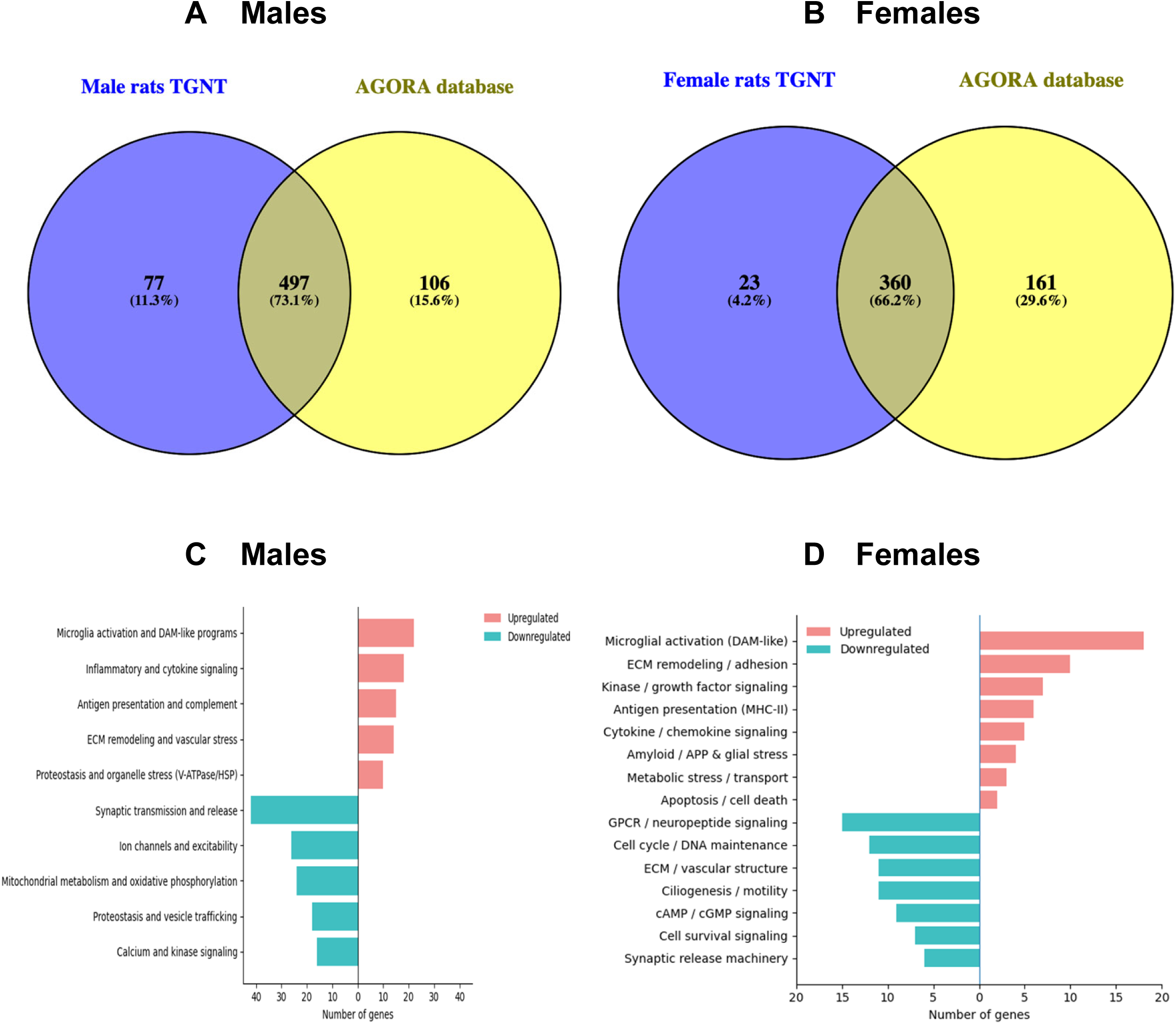
Human core AD transcriptional pathways are conserved in the TgF344-AD rat hippocampus. **(A, B)** Venn diagrams depicting overlap between TGNT hippocampal DEGs and human AD-associated genes from the AMP-AD AGORA database (>900 candidate targets) in males **(A,** 497/574 DEGs, 73.1%) and females **(B,** 360/383 DEGs, 66.2%**), (C, D)** Pathway enrichment analysis of the cross-species conserved gene sets in males **(C)** and females **(D)**, showing upregulated (red) and downregulated (blue) pathways. Conserved pathways faithfully recapitulate the principal biological programs identified in the full TGNT transcriptome (Fig. 1), spanning neuroinflammation, ECM and vascular remodeling, metabolic and proteostatic stress, and synaptic failure in both sexes. Conserved gene sets are provided in Supplementary Tables 5 and 6. TGNT, transgenic not treated, 5 rats per each of the two groups.

Pathway enrichment of these cross-species conserved gene sets recapitulated the principal biological programs identified in the full TGNT hippocampal transcriptomes. In males (Fig. 2C), conserved upregulated pathways encompassed microglial activation and DAM-like programs, inflammatory and cytokine signaling, complement activation, antigen presentation, ECM and vascular remodeling, and proteostatic stress responses, alongside coordinated suppression of synaptic transmission and vesicle release, neuronal excitability, mitochondrial metabolism and oxidative phosphorylation, vesicle trafficking, and calcium-dependent signaling. In females (Fig. 2D), conserved signatures similarly captured microglial activation, MHC-II–mediated antigen presentation, cytokine and kinase signaling, ECM remodeling, amyloid/APP-associated stress, and pro-apoptotic programs, with concordant downregulation of GPCR/neuropeptide signaling, cAMP/cGMP pathways, cell survival signaling, and synaptic release machinery.

The high degree of cross-species transcriptional concordance, spanning neuroinflammation, ECM and vascular remodeling, metabolic and proteostatic stress, and synaptic failure in both males and females, establishes that the TgF344-AD rat hippocampal transcriptome faithfully mirrors the core molecular architecture of human AD. These findings provide strong translational support for leveraging this model to dissect sex-dependent disease mechanisms and to evaluate candidate therapeutics targeting conserved pathological networks.

### Terazosin reverses AD transcriptional signatures in the TgF344-AD rat hippocampus

To determine whether chronic TZ treatment remodels the AD-associated hippocampal transcriptome, we compared TGNT and TGTR groups in both sexes. Unsupervised hierarchical clustering revealed clear treatment separation in males (Fig. 3A) and females (Fig. 3B), with top-ranked gene clusters displaying reciprocal expression patterns relative to TGNT, indicative of broad reversal of disease-associated transcriptional programs.

**Figure 3.**
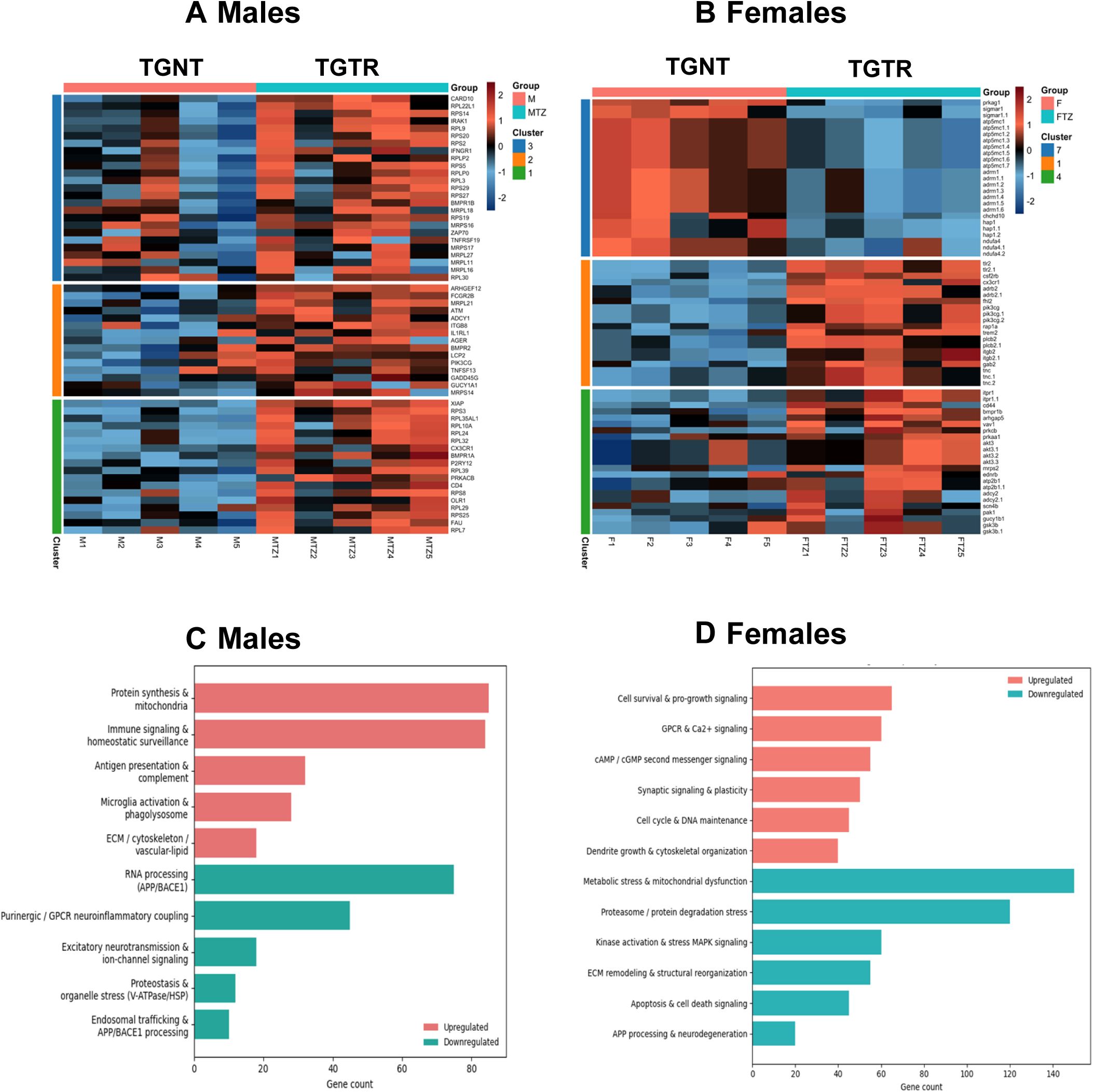
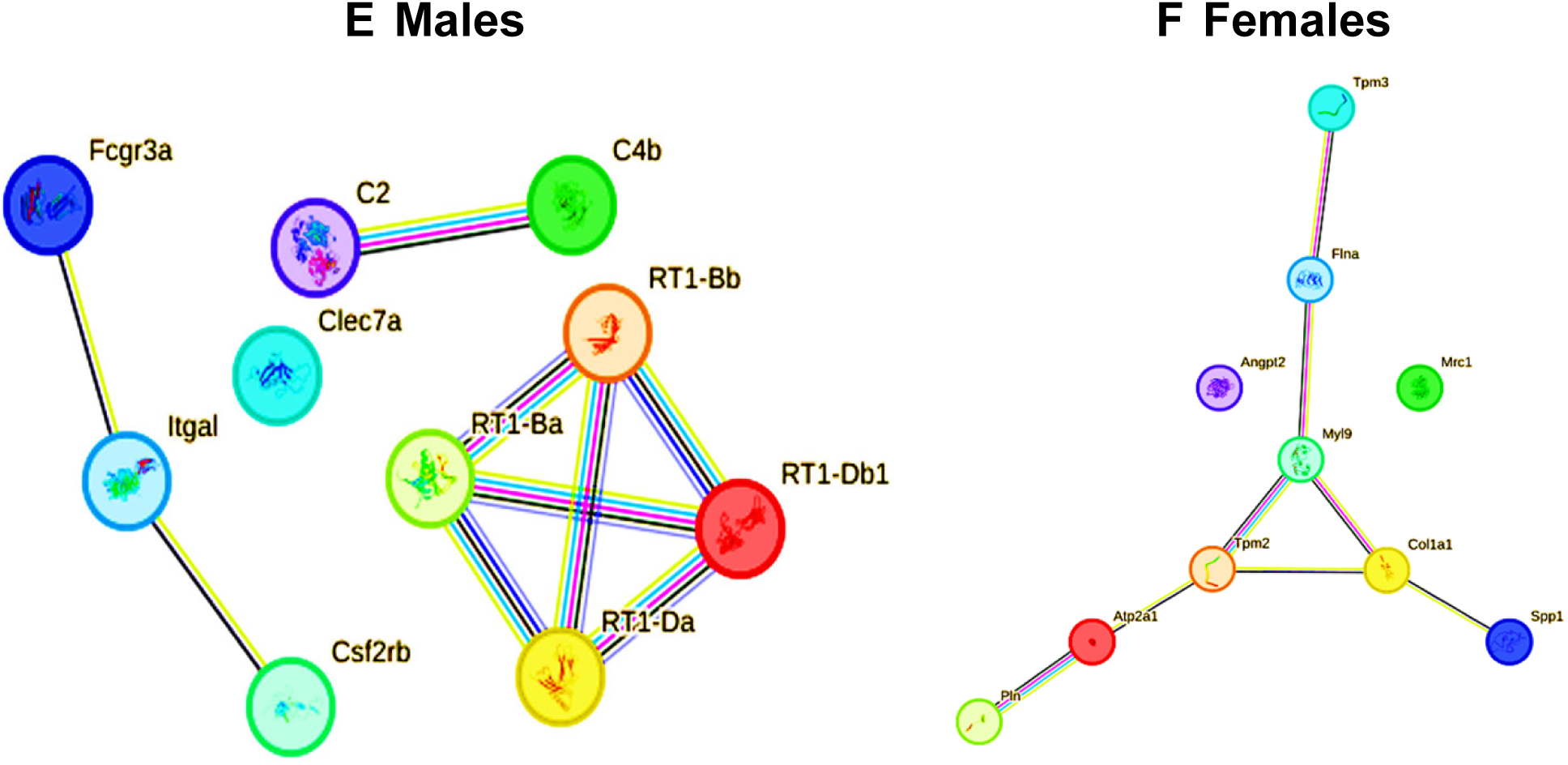
Terazosin reverses AD transcriptional signatures in the TgF344-AD rat hippocampus in a sex-dependent manner. **(A, B)** Unsupervised hierarchical clustering heatmaps of DEGs comparing non-treated (TGNT) and terazosin-treated (TGTR) transgenic rats in males **(A)** and females **(B)**, demonstrating genotype–treatment separation with reciprocal gene expression patterns indicative of broad transcriptional reversal. **(C, D)** Pathway enrichment analysis of TZ-regulated DEGs in males **(C)** and females **(D)**, (upregulated in red and downregulated in blue). In males, TZ upregulated mitochondrial protein synthesis, immune homeostatic surveillance, antigen presentation, complement, and ECM/vascular organization pathways, while suppressing RNA processing, proteostatic stress, purinergic/GPCR inflammatory signaling, and endosomal trafficking including APP/BACE1 processing. In females, TZ upregulated cell survival and pro-growth signaling, GPCR and second messenger pathways, synaptic plasticity, and dendritic cytoskeletal organization, while suppressing metabolic and mitochondrial stress, stress-MAPK/kinase cascades, proteasome-dependent degradation, ECM remodeling, and APP processing pathways. Upregulated pathways in red, and downregulated pathways in blue. Gene lists are provided in Supplementary Tables 7–10. **(E, F)** STRING protein–protein interaction networks of TZ-responsive DEGs in males **(E)** and females **(F)**. The male network is anchored by MHC class I/II genes (RT1-Ba, RT1-Bb, RT1-Db1, RT1-Da), complement components (C2, C4b), Fc receptor signaling (Fcgr3a), integrin signaling (Itgal), and innate immune regulators including Clec7a. The female network is organized around cytoskeletal and synaptic regulators (Tpm2, Tpm3, Myl9, Flna, Col1a1), calcium signaling components (Atp2a1, Pln), and ECM/neurovascular mediators (Spp1, Angpt2, Mrc1), supporting coordinated reinforcement of structural and synaptic stability. Nodes represent proteins encoded by TZ-modulated genes; node colors distinguish proteins and do not indicate expression direction. Edges represent predicted or experimentally supported functional associations. Light blue (curated databases), pink (experimental evidence), green (gene neighborhood), red (gene fusion), dark blue (gene co-occurrence), yellow (text mining), black (co-expression), and purple (protein homology). TZ, terazosin; TGNT, transgenic not treated; TGTR, transgenic TZ-treated, with 5 rats per each of the four groups.

Pathway enrichment analysis demonstrated that TZ systematically counteracted the major pathways dysregulated in TGNT rats (Fig. 1C, D) and in the conserved human-overlapping gene sets (Fig. 2C, D). In males (Fig. 3C), TZ upregulated mitochondrial protein synthesis, immune homeostatic surveillance, antigen presentation, complement pathways, and ECM/cytoskeletal/vascular-lipid organization, while suppressing RNA processing, purinergic and GPCR-coupled inflammatory signaling, excitatory neurotransmission, ion channel activity, proteostatic and organelle stress (V-ATPase/HSP), and endosomal trafficking including APP/BACE1 processing, thus collectively opposing the mitochondrial dysfunction, proteostatic stress, inflammatory dysregulation, and synaptic suppression that characterize TGNT males (Supplementary Tables 7 and 8). In females (Fig. 3D), TZ upregulated cell survival and pro-growth signaling, GPCR and second messenger pathways, synaptic signaling and plasticity, cell cycle/DNA maintenance, and dendritic cytoskeletal organization, while downregulating metabolic and mitochondrial stress, proteasome-dependent protein degradation, stress-MAPK and kinase signaling, ECM remodeling, and APP processing and neurodegeneration pathways, directly countering the metabolic stress, apoptotic activation, and synaptic impairment observed in TGNT females (Supplementary Tables 9 and 10). The mechanistic basis of these effects is twofold: modulation of mitochondrial, metabolic and purinergic pathways is consistent with TZ’s role as a PGK1 activator^40^, whereas effects on GPCR and cAMP/cGMP signaling might reflect its α₁-adrenergic receptor antagonism^39^.

To resolve the network architecture of TZ-responsive gene sets, we performed STRING protein–protein interaction analysis. In males (Fig. 3E), DEGs formed a tightly interconnected immune network anchored by MHC class I and II genes (RT1-Ba, RT1-Bb, RT1-Db1, RT1-Da), complement components (C2, C4b), Fc receptor signaling (Fcgr3a), integrin signaling (Itgal), and innate immune regulators (Clec7a, Csf2rb), collectively linking antigen presentation, complement activation, and phagocytic clearance into a coordinated immune surveillance module. In females (Fig. 3F), the interaction network was organized around cytoskeletal and synaptic regulators (Tpm2, Tpm3, Myl9, Flna, Col1a1), calcium signaling components (Atp2a1, Pln), and ECM/neurovascular mediators (Spp1, Angpt2, Mrc1), supporting coordinated reinforcement of structural and synaptic stability pathways.

Together, these findings demonstrate that TZ induces structured, sex-dependent transcriptional reprogramming of conserved AD-associated molecular networks in TgF344-AD rats, enhancing expression of mitochondrial genes and immune surveillance in males, and restoring synaptic and cell survival signaling in females, thus providing mechanistic evidence for broad reversal of disease-relevant hippocampal transcriptional pathology.

### Terazosin preserves LC-derived noradrenergic fibers in the hippocampus of female TgF344-AD rats

Having established broad transcriptional remodeling of synaptic, metabolic, and immune pathways in TGNT and TGTR rats (Figs. 1–3), we next examined whether these changes were accompanied by structural neuronal loss and whether TZ conferred neuroprotection at the cellular level. NeuN immunostaining revealed no significant effect of genotype [*F*(1,43) = 0.05, *P* = 0.82] or treatment [*F*(1,43) = 1.52, *P* = 0.22] on global hippocampal neuronal density at 11 months (Supplementary Table 11-15), indicating that overt neuronal loss was not yet manifested at this disease stage in either sex.

Given that LC noradrenergic neurons are among the earliest and most vulnerable population to degenerate in AD^8,9,16^, and that LC-derived projections densely innervate the hippocampus, we quantified norepinephrine transporter (NET) immunoreactivity as a marker of LC axonal integrity in the dorsal hippocampus. TGNT animals of both sexes exhibited significantly reduced NET-positive fiber density relative to WTNT controls, confirming progressive noradrenergic denervation (Fig. 4A–H). Strikingly, the magnitude of NET loss was greater in females than in males, consistent with reported heightened vulnerability of the female LC to AD-associated degeneration^33,34^.

**Figure 4.**
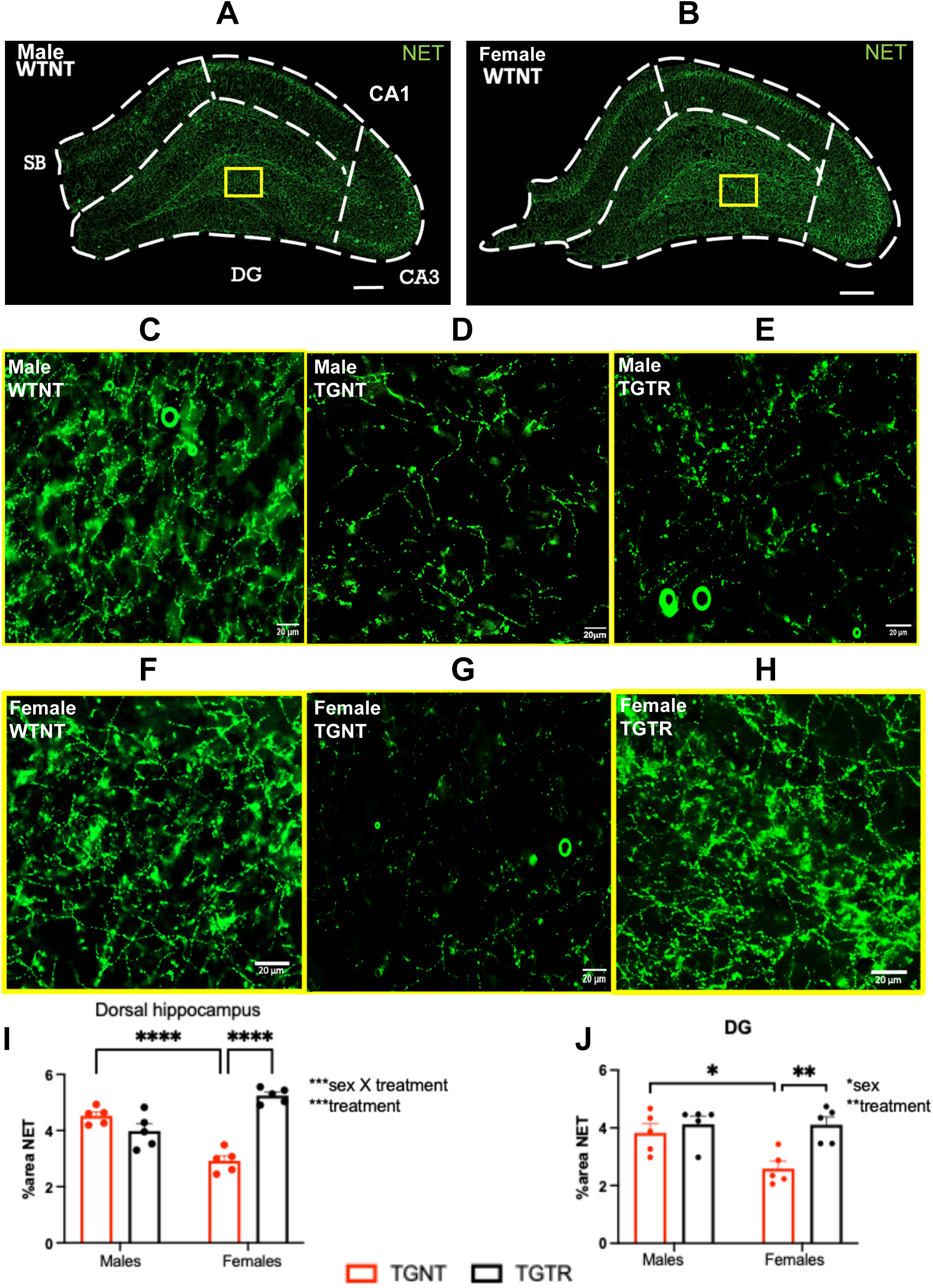
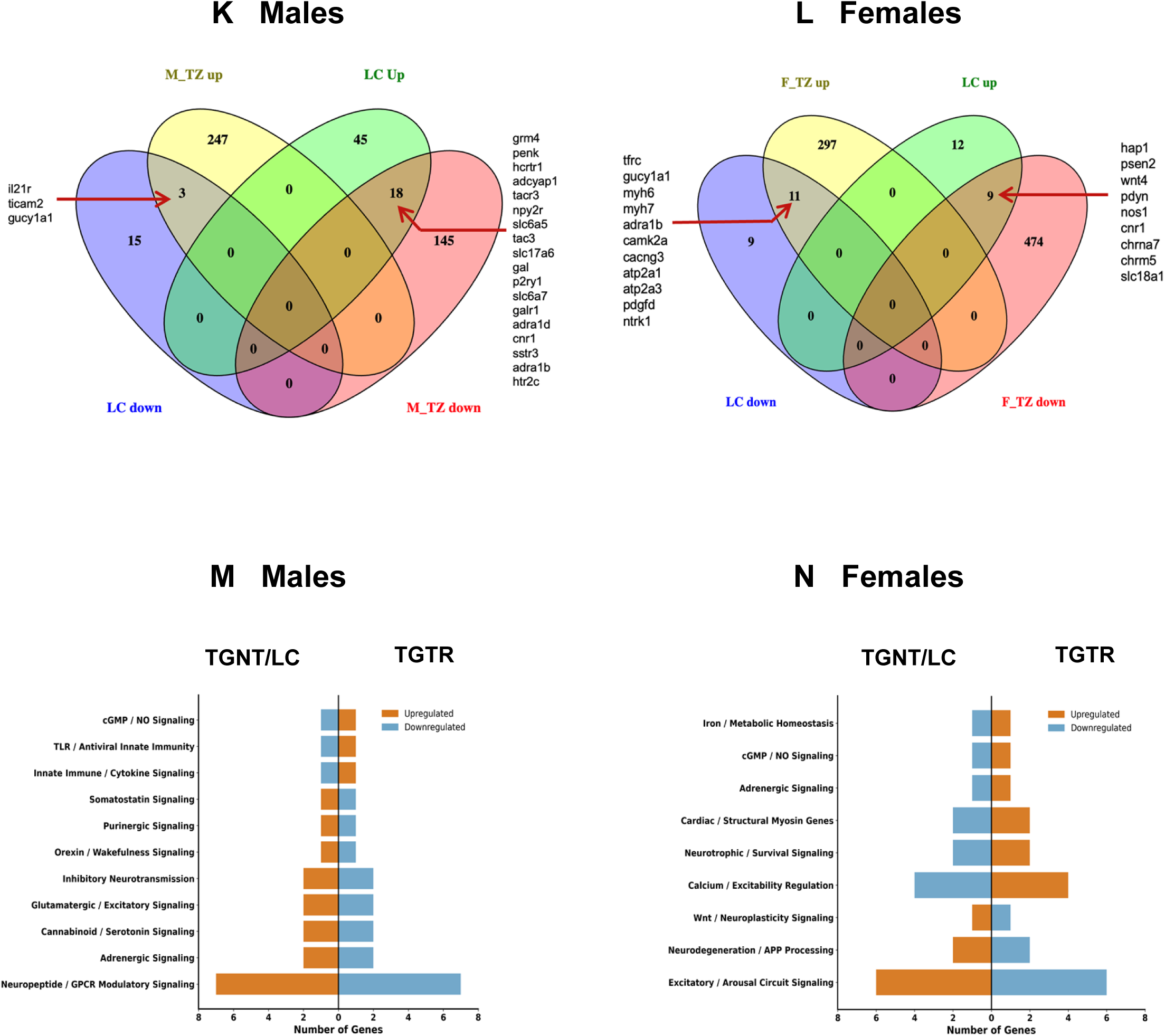
Terazosin preserves LC-hippocampal noradrenergic projections and reverses LC vulnerability-associated transcriptional signatures in a sex-dependent manner. **(A–H)** Representative widefield images of norepinephrine transporter (NET) immunostaining in the dorsal hippocampus of male and female WTNT, TGNT, and TGTR rats. Scale bars = 200 µm (A,B) and 20 μm (C-H). **(I, J)** Quantification of NET-positive fiber density across the whole dorsal hippocampus **(I)**, and dentate gyrus **(J)** by sex and treatment group in transgenic animals. Two-way ANOVA (N = 5 rats per group) revealed significant sex × treatment interactions [whole hippocampus: *F*(1,16) = 58.71, *P* < 0.0001; dentate gyrus: *F*(1,16) = 4.65, *P* = 0.047]. TZ robustly preserved NET fiber density in female but not male TGTR rats. Data are mean ± SEM; \**P* < 0.05, \*\**P* < 0.01, \*\*\**P* < 0.001. **(K, L)** Depiction of TZ-mediated reversal of LC vulnerability-associated transcriptional signatures [cross-referenced from^9^] in male **(K)** and female **(L)** TGTR rats. **(M, N)** Pathway enrichment analysis of TZ-reversed LC-associated genes in males **(M)** and females **(N),** upregulated pathways in red, and downregulated pathways in blue. Male-reversed pathways include neuropeptide/GPCR and adrenergic signaling, glutamatergic and inhibitory neurotransmission, somatostatin, cannabinoid/serotonin and orexin/wakefulness circuits, and innate immune/TLR and cGMP/NO signaling. Female-reversed pathways encompass synaptic and excitatory/arousal signaling, Wnt/neuroplasticity, neurotrophic/survival pathways, calcium/excitability regulation, adrenergic and cGMP/NO signaling, iron/metabolic homeostasis, and APP processing. Full gene lists are provided in Supplementary Tables 21-22. LC, locus coeruleus; TZ, terazosin; CA1, Cornu Ammonis 1; CA3, Cornu Ammonis 3; DG, Dentate Gyrus; SB, Subiculum; WTNT, wild-type not treated; TGNT, transgenic not treated; TGTR, transgenic terazosin-treated.

Three-way ANOVA (genotype × sex × treatment) confirmed a significant main effect of genotype [*F*(1,16) = 122.7, *P* < 0.0001], significant treatment × genotype [*F*(1,16) = 15.01, *P* = 0.0013], and treatment × sex × genotype interaction effect [*F*(1,16) = 31.57, *P* < 0.0001]. Two-way ANOVA restricted to transgenic animals revealed a significant sex × treatment interaction across the whole dorsal hippocampus [*F*(1,16) = 58.71, *P* < 0.0001] and dentate gyrus [*F*(1,16) = 4.65, *P* = 0.047] (Fig. 4I, J). TZ treatment robustly preserved NET-positive fiber density in female TGTR rats but conferred no significant protection in males. No significant effects were detected in CA1, CA3, or subiculum subregions (Supplementary table 16-18), indicating anatomical specificity of both noradrenergic vulnerability and TZ-mediated preservation.

Together, these findings demonstrate that at 11 months of age, TgF344-AD rats exhibit selective degeneration of LC-derived hippocampal projections without overt global neuronal loss, with females displaying heightened noradrenergic vulnerability. Chronic TZ treatment specifically preserved hippocampal noradrenergic fiber integrity in females, identifying a sex-selective neuroprotective mechanism with direct relevance to the LC–hippocampal circuit disruption that characterizes early AD.

### Terazosin reverses transcriptional signatures associated with locus coeruleus vulnerability in TgF344-AD rats

To determine whether TZ modulates transcriptional programs linked to LC neuronal vulnerability, we cross-referenced hippocampal DEGs from TgF344-AD rats (TGNT) with LC-specific DEGs defined by Ehrenberg et al.^9^ across human Braak stages, a dataset capturing the transcriptional basis of selective LC degeneration in human AD (refer to methods section for details). This analysis identified 81 overlapping genes in males and 41 in females (Supplementary Tables 19 and 20). In TGNT males, 63 of these genes were upregulated and 18 downregulated; in TGNT females, 21 were upregulated and 20 downregulated, indicating sex-divergent engagement of LC vulnerability programs in the TgF344-AD hippocampus.

TZ treatment partially normalized these LC-associated transcriptional signatures in both sexes. In TGTR males (Fig. 4K), 18 of 63 upregulated genes were reversed to downregulation and 3 of 18 downregulated genes were restored; in TGTR females (Fig. 4L), 9 of 21 upregulated genes were downregulated and 11 of 20 downregulated genes were upregulated. Pathway enrichment of TZ-reversed LC genes revealed sex-divergent functional targets. In males (Fig. 4M), reversed pathways encompassed neuropeptide/GPCR and adrenergic signaling, glutamatergic and inhibitory neurotransmission, somatostatin, cannabinoid/serotonin and orexin/wakefulness circuits, and innate immune/TLR and cGMP/NO signaling. In females (Fig. 4N), reversed pathways mapped to synaptic and excitatory/arousal signaling, Wnt/neuroplasticity and neurotrophic/survival pathways, calcium/excitability regulation, adrenergic and cGMP/NO signaling, iron/metabolic homeostasis, and APP processing (Supplementary Tables 21 and 22). Notably, antioxidant signaling pathways, a key vulnerability signature identified by Ehrenberg et al.^9^, were shifted in the opposite direction by TZ in both sexes, indicating direct engagement of LC-relevant stress response programs.

Together, these findings demonstrate that TZ partially reverses LC vulnerability-associated transcriptional programs in TgF344-AD rats, with greater normalization in females. This molecular realignment mechanistically corroborates the sex-selective preservation of hippocampal noradrenergic projections observed following TZ treatment (Fig. 4) and identifies restoration of LC-associated transcriptional homeostasis as a candidate mechanism underlying TZ-mediated neuroprotection.

### Terazosin reduces amyloid plaque burden in the TgF344-AD rat hippocampus independently of sex

We next quantified Aβ plaque burden in the dorsal hippocampus of TGNT and TGTR rats. Baseline amyloid pathology was comparable between sexes in TGNT animals (Fig. 5A–D). Two-way ANOVA revealed that TZ treatment robustly reduced Aβ plaque area in the CA1 region [*F*(1,19) = 34.01, *P* < 0.0001; Fig. 5E] and subiculum [*F*(1,19) = 22.26, *P* = 0.0001; Fig. 5F], with no significant main effect of sex and no sex × treatment interaction in either subregion, indicating that TZ-mediated plaque reduction is sex-independent. No significant genotype or treatment effects were detected in other hippocampal subregions (Supplementary Tables 23–25), demonstrating regional specificity of the anti-amyloid effect.

**Figure 5.**
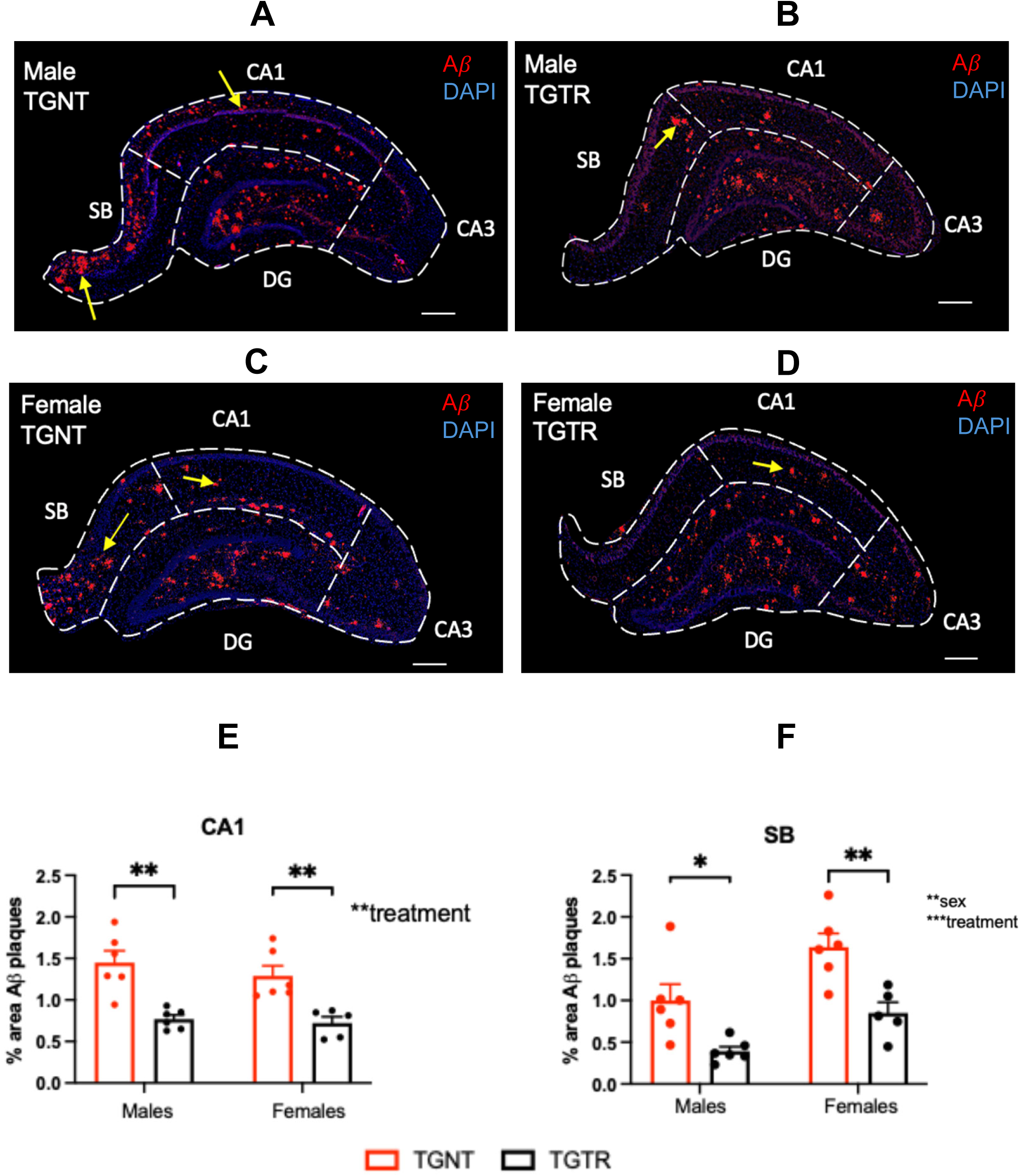
Terazosin reduces amyloid-β plaque burden in hippocampal CA1 and subiculum independently of sex. **(A–D)** Representative widefield images of Aβ immunostaining in the dorsal hippocampus (CA1 and subiculum) of male and female TGNT and TGTR rats. Scale bar = 200 µm. **(E, F)** Quantification of Aβ plaque area in CA1 **(E)** and subiculum **(F)** across all groups. Two-way ANOVA (N = 5-6 rats per group) revealed a significant main effect of treatment in CA1 [*F*(1,19) = 34.01, *P* < 0.0001] and subiculum [*F*(1,19) = 22.26, *P* = 0.0001], with no significant main effect of sex, and no sex × treatment interaction in either subregion. Data are mean ± SEM; \**P* < 0.05, \*\**P* < 0.01, \*\*\**P* < 0.001. No significant treatment effects were detected in other hippocampal subregions (Supplementary Tables 23–25). CA1, Cornu Ammonis 1; CA3, Cornu Ammonis 3; DG, dentate gyrus; SB, subiculum; TGNT, transgenic not treated; TGTR, transgenic terazosin treated.

These findings establish that chronic TZ treatment reduces hippocampal Aβ plaque burden in both male and female TgF344-AD rats, with effects confined to CA1 and subiculum, regions critically implicated in memory encoding and AD-associated neurodegeneration, providing in vivo evidence for TZ-mediated attenuation of amyloid pathology independent of sex.

### Terazosin attenuates tau hyperphosphorylation in female TgF344-AD rats

We next examined whether TZ treatment attenuates tau hyperphosphorylation using AT8 immunostaining to quantify hyperphosphorylated tau (pTau) deposition in the dorsal hippocampus. TGNT animals of both sexes exhibited elevated AT8-positive signal relative to WTNT controls (Fig. 6A–H). Two-way ANOVA across the whole dorsal hippocampus revealed a significant main effect of treatment [*F*(1,16) = 5.187, *P* = 0.0368; Fig. 6I]. Although no significant sex × treatment interaction [F(1,16) = 3.96, P = 0.064] was detected, post hoc analysis showed that TZ significantly reduced pTAU levels only in female TgF344-AD rats (p = 0.0382). Analysis of the dentate gyrus confirmed a significant sex × treatment interaction [*F* (1, 16) = 6.454, *P=*0.0218] and main effect of treatment [*F* (1, 16) = 5.952, *P*=0.0267; Fig. 6J], with TZ selectively reducing AT8-positive counts in females. No significant effects were detected in other hippocampal subregions (Supplementary Tables 26–29), indicating anatomical specificity of tau pathology suppression.

**Figure 6.**
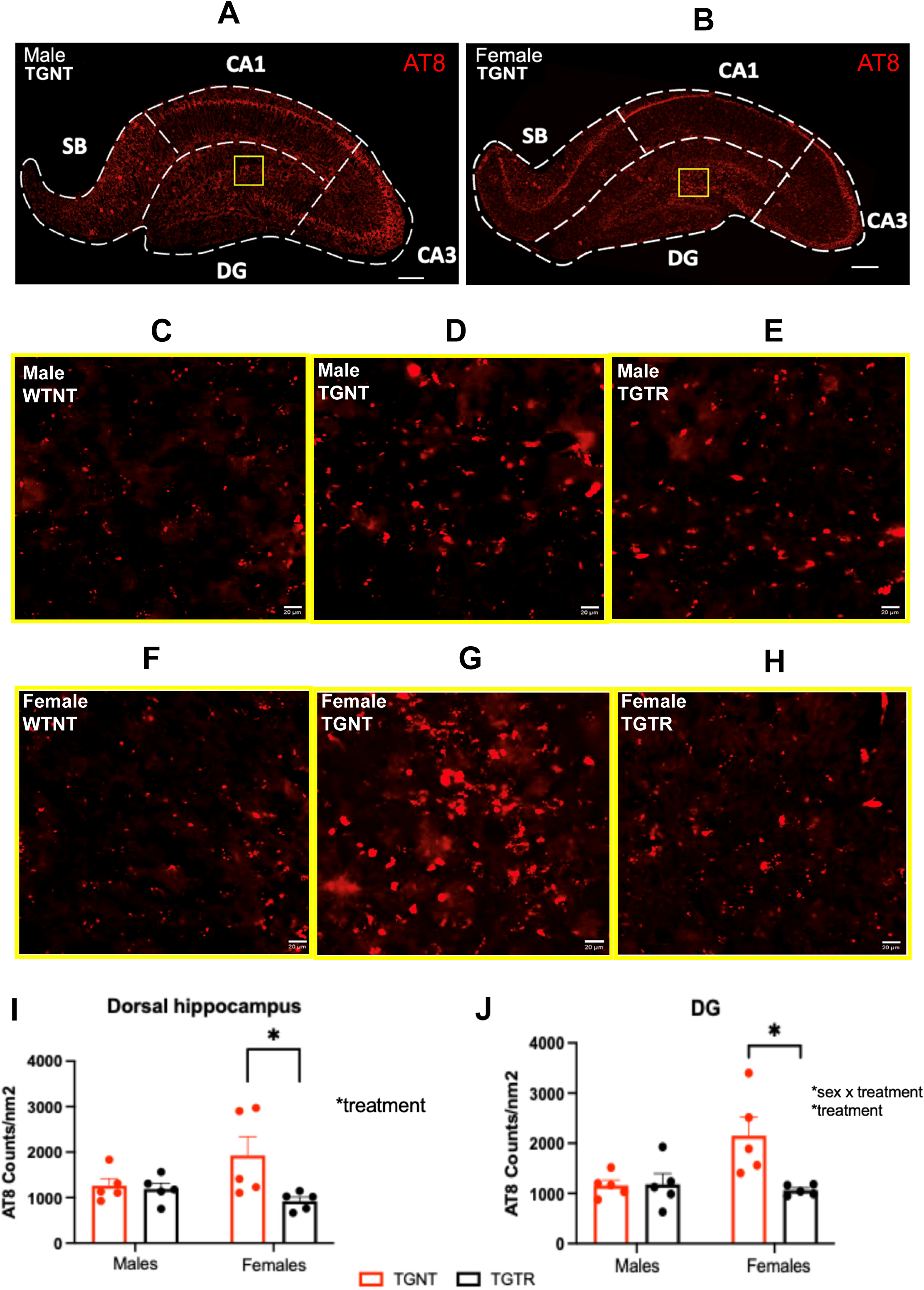
Terazosin selectively attenuates tau hyperphosphorylation in female TgF344-AD rats. **(A–H)** Representative widefield images of AT8 (phospho-tau Ser202/Thr205) immunostaining in the dorsal hippocampus of male and female WTNT, TGNT, and TGTR rats. A, B - Scale bar = 200 µm, C-H - Scale bar = 20 µm. **(I, J)** Quantification of AT8-positive signal across the whole dorsal hippocampus **(I)** and dentate gyrus **(J)**. Two-way ANOVA (N = 5 rats per group) of the whole dorsal hippocampus revealed a significant main effect of treatment [*F*(1,16) = 5.32, *P* = 0.035] driven exclusively by females. Dentate gyrus analysis confirmed a significant sex × treatment interaction [*F*(1,20) = 7.00, *P* = 0.016] and main effect of treatment [*F*(1,20) = 6.46, *P* = 0.019], with TZ selectively reducing AT8-positive counts in females. Data are mean ± SEM; \**P* < 0.05. No significant effects were detected in other subregions (Supplementary Tables 26–29). CA1, Cornu Ammonis 1; CA3, Cornu Ammonis 3; DG, Dentate Gyrus; SB, Subiculum; WTNT, wild-type not treated; TGNT, transgenic not treated; TGTR, transgenic terazosin-treated.

The sex-selective reduction in pTau is mechanistically coherent with the transcriptomic profile of TZ-treated females, which showed marked downregulation of stress-MAPK and kinase signaling pathways (Fig. 3D), upstream regulators of tau hyperphosphorylation. Together, these data identify female-specific attenuation of hippocampal tau pathology as a key neuroprotective outcome of TZ treatment, adding pathological specificity to the sex-dependent molecular and noradrenergic effects observed in this study.

### Terazosin drives sex-specific microglial morphological activation in TgF344-AD rats

To determine whether TZ modulates microglial states in the dorsal hippocampus, we classified microglia as ramified (homeostatic surveillance), reactive, or amoeboid (phagocytic activation) based on form factor circularity (Methods section). Three-way ANOVA confirmed elevated amoeboid proportions in TGNT animals relative to WTNT controls [genotype effect: *F*(1,39) = 14.97, *P* = 0.0004], reflecting heightened baseline inflammatory tone in transgenic rats at 11 months. A significant treatment × genotype interaction effect was also observed[*F*(1,39) = 6.52, *P* = 0.015].

Two-way ANOVA within transgenic animals revealed significant sex × treatment interactions in both CA1 [*F* (1, 20) = 20.51, *P*=0.0002; Fig. 7K] and dentate gyrus [*F*(1,20) = 17.68, *P* = 0.0004; Fig. 7L], with significant main effects of sex and treatment in both CA1 [*F* (1, 20) = 6.683, *P* =0.0177; *F* (1, 20) = 6.606, *P* = 0.0183 respectively] and DG [*F* (1, 20) = 16.91, *P* =0.0005; *F* (1, 20) = 5.097, *P*=0.0353 respectively] subregions. In males, TZ shifted the microglial balance toward amoeboid morphology in both CA1 and dentate gyrus (Fig. 7G–J), indicating enhanced phagocytic activation. Females showed no significant morphological redistribution in either subregion, demonstrating a male-specific effect of TZ on microglial state. No significant effects were detected in other hippocampal subregions (Supplementary tables 30-32).

**Figure 7.**
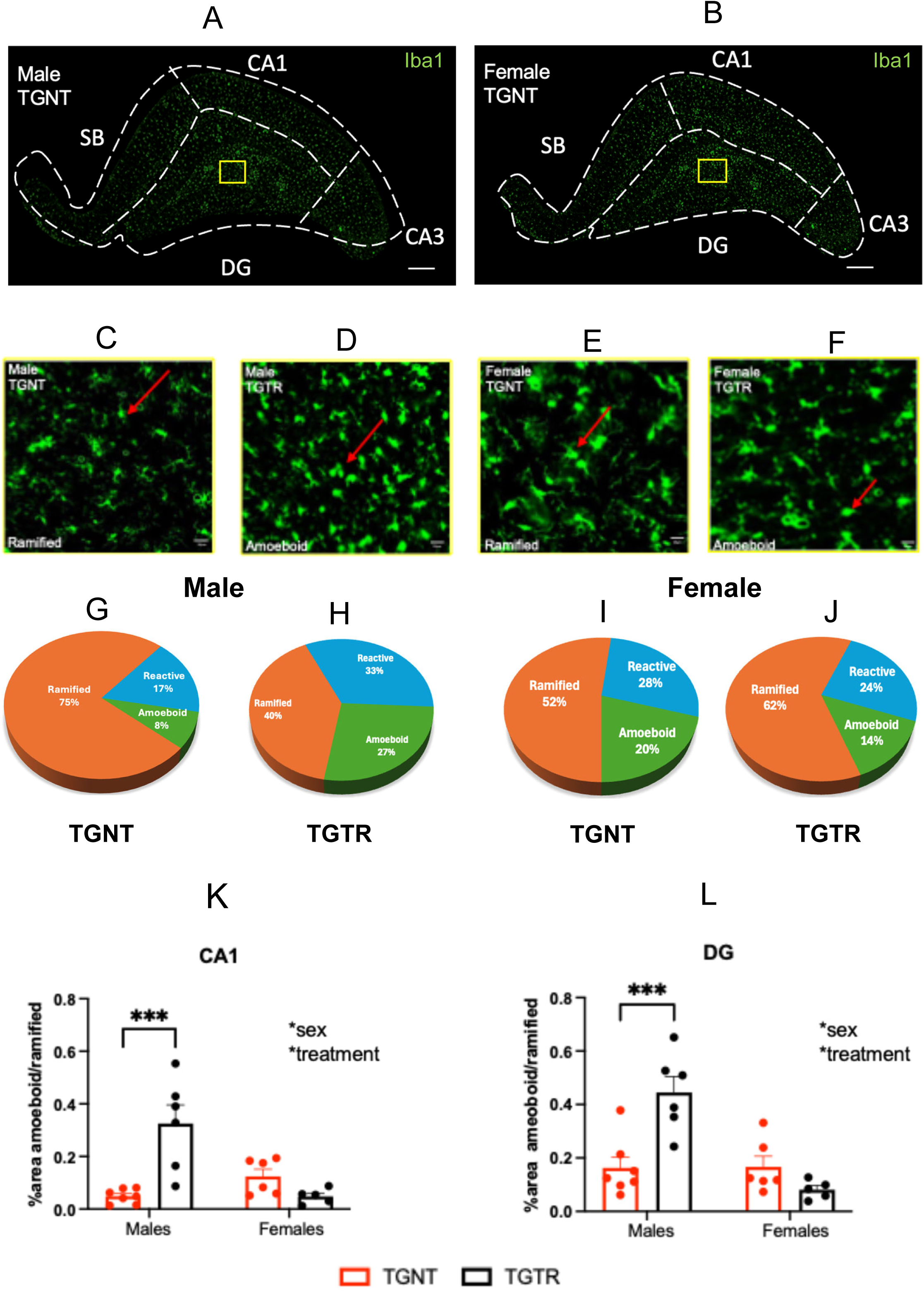
Terazosin drives sex-specific microglial activation in TgF344-AD male rats. **(A–F)** Representative widefield images of Iba1-immunostained microglia in the dorsal hippocampus (**A, B**) and dentate gyrus (**C-F**) of male and female WTNT (**A, B**), TGNT, and TGTR rats (**C-F**), illustrating the morphological spectrum from ramified (homeostatic) to amoeboid (phagocytic) states. A,B Scale bars = 200µm, C-F- Scale bars = 20 µm**. (G–J)** Pie charts depicting the proportional distribution of ramified (orange), reactive (blue), and amoeboid (green) microglial morphologies in the dentate gyrus, shown separately by sex and treatment group. **(K, L)** Quantification of amoeboid (normalized to ramified) microglial levels in CA1 **(K)** and dentate gyrus **(L)** of transgenic animals by sex and treatment. Two-way ANOVA (N = 5-7 rats per group) revealed significant sex × treatment interactions in both subregions [CA1: *F* (1, 20) = 20.51, *P*=0.0002; Fig. 7K] and [DG: *F*(1,20) = 17.68, *P* = 0.0004; Fig. 7L]. TZ significantly increased amoeboid levels in male but not female TGTR rats. Data are mean ± SEM (Supplementary Tables 30-32). \**P* < 0.05, \*\**P* < 0.01, \*\*\**P* < 0.001. Tz, terazosin; CA1, Cornu Ammonis 1; CA3, Cornu Ammonis 3; DG, dentate gyrus; SB, subiculum; TGNT, transgenic not treated; TGTR, transgenic terazosin-treated.

This sex-selective promotion of microglial activation in males is consistent with the transcriptomic enrichment of immune surveillance, complement, and antigen presentation pathways observed in TGTR males (Fig. 3C, E), suggesting that TZ engages a coordinated neuroimmune response program preferentially in males, a mechanistic divergence that may underlie the distinct neuroprotective outcomes observed across sexes.

### Terazosin rescues spatial learning deficits in female TgF344-AD rats

To determine whether the molecular and histopathological protection conferred by TZ translates into functional cognitive benefit, we assessed hippocampal-dependent spatial learning and memory using the active place avoidance task (aPAT). Three-way ANOVA across six training sessions revealed significant main effects of training [*F*(4.15,174.3) = 9.01, *P* < 0.0001], genotype [*F*(1,42) = 8.71, *P* = 0.0052], and treatment [*F*(1,42) = 8.41, *P* = 0.0059]. TGNT rats exhibited significantly shorter latency to first entrance to shock zone relative to WTNT controls (Fig. 8A), confirming AD-associated spatial learning impairment.

**Figure 8.**
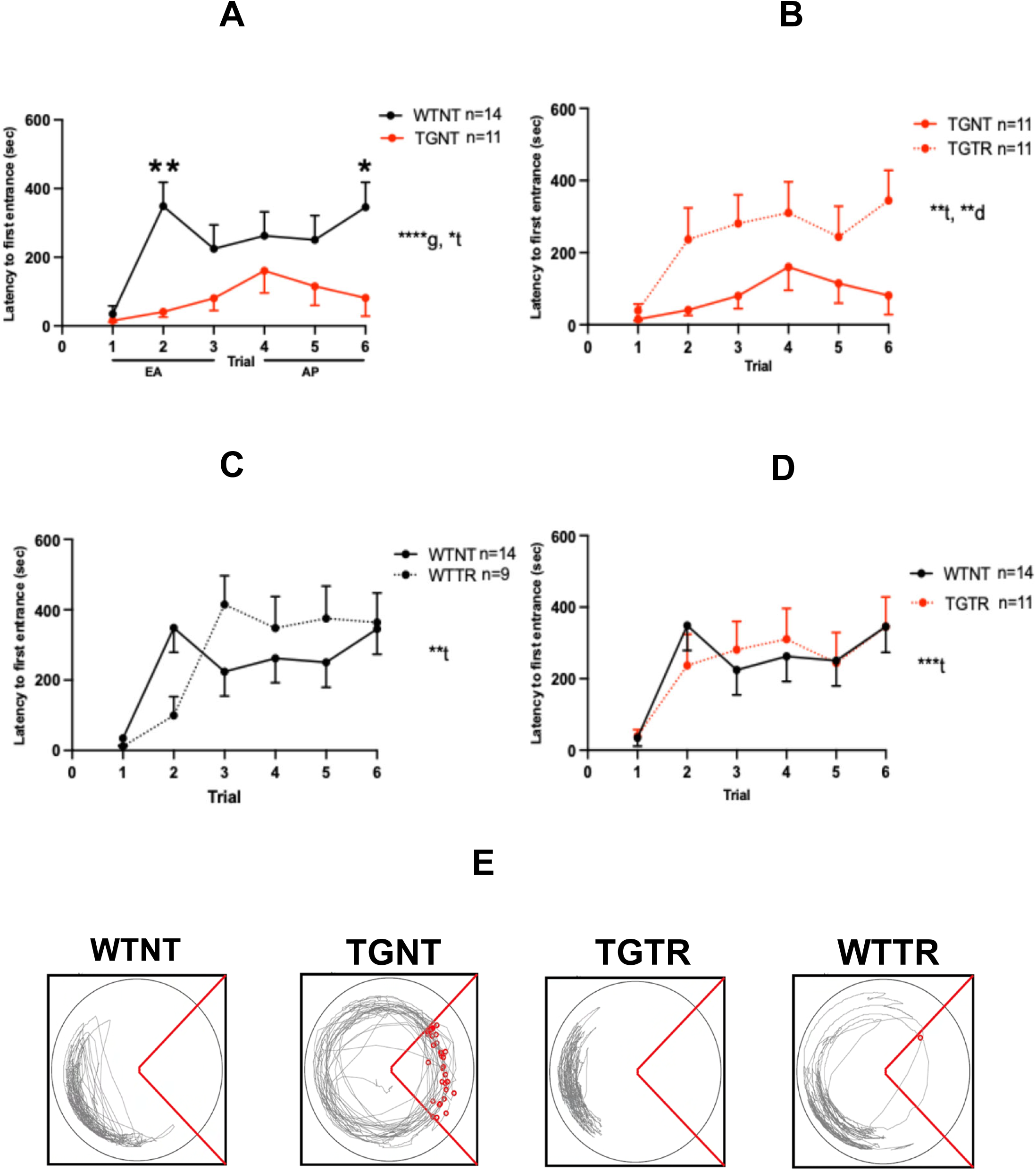
Terazosin rescues hippocampal-dependent spatial learning in TgF344-AD rats. **(A)** Active place avoidance task (aPAT) performance across six training sessions in all groups (both sexes combined). TGNT rats exhibited significantly shorter latencies to shock zone entry than WTNT controls, confirming AD-associated spatial learning impairment. Three-way ANOVA significant main effects of training [*F*(4.15,174.3) = 9.01, *P* < 0.0001], genotype [*F*(1,42) = 8.71, *P* = 0.0052], and treatment [*F*(1,42) = 8.41, *P* = 0.0059]. Post hoc analysis identified significant WTNT vs. TGNT differences at trials 2 (*P* = 0.004) and 6 (*P* = 0.043). **(B)** TZ treatment in TGNT animals significantly improved performance during both early acquisition (trials 1–3; *F*(1,20) = 10.00, *P* = 0.0049) and asymptotic performance (trials 4–6; *F*(1,20) = 5.04, *P* = 0.036). **(C)** TZ had no effect on WT animals in either phase, confirming disease-state specificity. **(D)** TZ-treated TGNT (TGTR) rats performed indistinguishably from WTNT controls [*F*(1,23) = 0.0009, *P* = 0.976], confirming full rescue to wild-type levels. Data are mean ± SEM, (Supplementary Tables 33 and 34). \**P* < 0.05, \*\**P* < 0.01, \*\*\**P* < 0.001, \*\*\*\**P* < 0.0001. Long-term memory and locomotor activity were unaffected by genotype or treatment. **(E)** Representative trajectory plots during task performance for WTNT, TGNT, TGTR, and WTTR rats (female, trial 3). The red sector indicates the shock zone. TZ, terazosin; g = genotype; t = trial; d = drug effect; WTNT, wild-type not treated (N = 14); WTTR, wild-type terazosin-treated (N = 9); TGNT, transgenic not treated (N = 11); TGTR, transgenic terazosin-treated (N = 11).

To resolve learning dynamics, trials were grouped into early acquisition (EA; trials 1–3) and asymptotic performance (AP; trials 4–6). TGNT rats performed significantly worse than WTNT controls in both phases [EA: *F*(1,23) = 18.87, *P* = 0.0002; AP: *F*(1,23) = 11.62, *P* = 0.0024], with post hoc analysis identifying significant group differences at trials 2 and 6 (*P* = 0.004 and *P* = 0.043, respectively; Fig. 8A). TZ treatment significantly improved TGNT performance in both EA [*F*(1,20) = 10.00, *P* = 0.0049] and AP [*F*(1,20) = 5.04, *P* = 0.036; Fig. 8B], restoring performance to WTNT levels [*F*(1,23) = 0.0009, *P* = 0.976; Fig. 8D]. TZ had no effect on WT animals in either phase (Fig. 8C), confirming specificity to the disease state. Long-term memory and locomotor activity were unaffected by genotype or treatment (Supplementary Tables 28 and 29), excluding confounds of altered mobility or motivation.

Four-way ANOVA incorporating sex as a variable revealed that the cognitive rescue was driven predominantly by females: TZ produced a significant treatment main effect in female TgF344-AD rats [*F*(1,38) = 15.52, *P* = 0.003] but not in males [*F*(1,38) = 0.694, *P* = 0.426] or in WT animals of either sex. This sex-selective behavioral rescue converges with the female-predominant preservation of noradrenergic projections, attenuation of tau hyperphosphorylation, and transcriptomic remodeling observed in this study, collectively identifying a coherent, sex-specific neuroprotective program through which TZ restored hippocampal cognitive function in TgF344-AD rats.

## DISCUSSION

This study establishes that chronic TZ treatment initiated prior to overt pathology remodeled the AD hippocampal transcriptome, preserved LC–hippocampal noradrenergic integrity, attenuated amyloid and tau pathology, and restored spatial cognition, with sex-specific mechanistic and functional profiles throughout. Outcomes were assessed at a stage paralleling the prodromal window of human AD^47–49^, when intervention carries the greatest potential. These data situate AD as a systems-level failure of neuronal–glial homeostasis^50–52^ and position TZ as a multi-axis repurposing candidate whose efficacy is sex-dependent.

At 11 months, the TgF344-AD hippocampal transcriptome recapitulated core molecular signatures of human AD across synaptic, immune, metabolic, proteostatic, and vascular domains^53–55^, validating the model’s cross-species reliability. Sex-divergent pathway organization was a defining feature. Males exhibited inflammatory activation, complement signaling, vascular remodeling, and ciliogenesis alongside suppression of synaptic and transcriptional homeostasis. Females showed metabolic and kinase stress, synaptic vulnerability, and dysregulated cell survival programs. These distinctions align with evidence that males and females follow fundamentally different molecular trajectories in AD^27,29,32,33^.

In males, TZ enhanced immune and complement signaling while suppressing RNA-processing pathways linked to amyloidogenic transcript stabilization^56–58^ and proteostasis stress, systems whose impairment drives AD progression^59–61^ and compromises microglial phagocytic capacity^62,63^. TZ also upregulated vascular–lipid, ECM, and cytoskeletal programs, suggesting remodeling of neurovascular components increasingly implicated in mixed dementia and vascular contributions to AD^52,64–66^.

Microglia are critical determinants of AD trajectory. Early phagocytic activation supports plaque containment and limits neuronal damage^11,12,51,67–70^. STRING network analysis in TZ-treated males identified immune interaction clusters anchored by Clec7a, a microglial gene that directly restrains amyloid-β plaque pathology^71^. Clec7a-anchored clustering indicates that TZ drives coordinated phagocytic engagement rather than generalized neuroinflammation. The male-specific shift toward amoeboid morphology corroborates this at the cellular level. Reduced amyloid burden in CA1 and subiculum provides the histopathological endpoint, linking transcriptional reprogramming to measurable plaque attenuation.

In females, TZ produced a mechanistically distinct response. Upregulation of survival, synaptic plasticity, and GPCR/second messenger signaling was accompanied by downregulation of metabolic stress, stress-MAPK/kinase cascades, and APP processing pathways. Mitochondrial dysfunction is an early driver of neurodegeneration^29,64^, and TZ’s PGK1-activating properties directly augment cellular ATP production^41^, providing a bioenergetic basis for this transcriptional shift. Stress-activated MAPK cascades drive aberrant tau hyperphosphorylation^72,73^ and show heightened activity in female TgF344-AD populations^74,75^. Their downregulation in TZ-treated TgF344-AD females directly explains the selective reduction in AT8-positive tau pathology. Preserved hippocampal noradrenergic fiber density confirmed that transcriptional reinforcement of synaptic and survival programs extends to structural circuit protection.

The locus coeruleus (LC) is among the earliest neuronal populations to degenerate in AD^15,19,24,76,77^. Overlap between TgF344-AD hippocampal DEGs and human LC vulnerability gene sets^9^ revealed partial transcriptional reversal following TZ treatment, more pronounced in females. Adrenergic signaling showed sex-specific directionality. We hypothesize that males and females differ fundamentally in their adrenergic disease state: in males, excessive compensatory noradrenergic signaling drives early circuit hyperactivity, such that its attenuation restores network homeostasis^16,78^; in females, declining adrenergic tone renders synaptic integrity dependent on its preservation^78^. This divergence is consistent with TZ’s dual α1-adrenergic antagonist and PGK1 activator profile operating in distinct sex-specific neuromodulatory contexts and represents a priority for direct mechanistic follow-up.

The behavioral outcome (analyzed by a 4-way ANOVA) reflected this sex-specific biology. Female TgF344-AD rats treated with TZ (TGTR) showed restoration of spatial learning to wild-type levels. Males did not. TZ had no effect in wild-type animals of either sex, confirming that cognitive rescue reflects correction of disease-associated impairment rather than nonspecific enhancement^27,47^. The female-predominant benefit converges with selective noradrenergic fiber preservation, hyperphosphorylated tau attenuation, and transcriptomic remodeling, collectively delineating a coherent, sex-specific neuroprotective program.

The absence of cognitive rescue in male TgF344-AD rats warrants consideration. Several non-mutually exclusive explanations exist. First, the active place avoidance task indexes hippocampal spatial learning, but male TgF344-AD rats may express cognitive deficits preferentially in other domains, such as executive function, reversal learning, or anxiety-related behavior, that were not assessed here. Second, 11 months may not be the optimal timepoint for detecting TZ-mediated behavioral rescue in males. The male transcriptome showed stronger inflammatory and vascular remodeling signatures than females, potentially reflecting a more advanced or mechanistically distinct disease state that requires earlier intervention or longer treatment duration to achieve functional benefit. Third, the dose employed was selected based on human clinical exposure equivalence and demonstrated clear efficacy in female rats; males may require dose adjustment to engage the same protective threshold, particularly if sex differences in adrenergic receptor density, drug metabolism, or PGK1 activity alter the pharmacodynamic response. Importantly, TZ reduced amyloid plaque burden in males, confirming biological activity in the male TgF344-AD brain. The dissociation between pathological attenuation and behavioral rescue in males suggests that amyloid reduction alone is insufficient for cognitive improvement at this stage, a conclusion consistent with findings from anti-amyloid clinical trials^79,80^, and that the noradrenergic and tau-related mechanisms more prominent in females may be the critical drivers of functional recovery.

Current anti-amyloid immunotherapies reduce plaque burden but frequently fail to produce meaningful cognitive benefit and carry substantial safety risks^35,79–81^. TZ operates at a different level of disease biology. As an FDA-approved compound^21^, TZ engages interconnected networks spanning bioenergetics, immune regulation, proteostasis, vascular remodeling, and neurotransmission. AD frequently coexists with vascular and mixed dementia pathologies^52,82^, and effective intervention may require pathway-level correction across neuronal, glial, and vascular compartments. The alignment of transcriptomic, structural, pathological, and behavioral outcomes here supports that framework.

Several limitations of the present study merit consideration. First, outcomes were assessed at a single timepoint, 11 months, corresponding to early-to-mid disease progression in the TgF344-AD model^45^. Whether TZ-mediated transcriptional, pathological, and behavioral effects are sustained at later stages, or whether earlier intervention confers more complete protection, remains to be determined. Longitudinal assessment across multiple timepoints would clarify the therapeutic window and durability of these effects. Validation in additional preclinical models, including those with distinct genetic backgrounds or tau-dominant pathology, would further strengthen cross-model validity. The sex-specific profiles identified here may also evolve differently as pathology progresses, an important consideration for future study design.

Second, microglial states were classified by morphological criteria, form factor circularity, which provides a validated index of activation but cannot resolve the full spectrum of microglial phenotypes now recognized in AD^11,61,67,68^. Transcriptomic subtypes, including homeostatic, disease-associated (DAM), and interferon-responsive states, have distinct functional consequences that morphology alone cannot distinguish. Future studies employing single-cell RNA sequencing, spatial transcriptomics, or multiplexed protein imaging will be needed to determine whether TZ promotes a bona fide phagocytic state in males and whether a distinct microglial program underlies the female neuroprotective profile. The STRING network data, with Clec7a as a major node in males^71^, provide a transcriptional foundation for these more targeted investigations.

Third, the transcriptomic analyses reported here are correlative and cannot establish causal directionality between pathway-level changes and the observed pathological and behavioral outcomes. While the mechanistic coherence across transcriptomic, cellular, and histopathological levels is compelling, direct validation experiments, including PGK1 activity assays^40,41^, MAPK expression quantification in sex-stratified samples, and targeted manipulation of network nodes such as Clec7a, will be required to establish causality. The consistency between TZ-modulated pathways in rats and conserved human AD gene sets^46^ provides a translational rationale for prioritizing these experiments, and the sex-specific STRING interaction networks define a tractable set of molecular targets for mechanistic follow-up.

Overall, our results establish terazosin as a systems-level disease-modifying agent capable of reversing AD-associated transcriptional programs, preserving noradrenergic circuitry, and restoring cognition. These effects are mechanistically coherent yet divergent across sexes. This sex-dependence is not a confound but a finding: it reveals that the noradrenergic system fails through distinct molecular trajectories in males and females, and that effective intervention must be calibrated accordingly. Given terazosin’s existing FDA approval and favorable tolerability, these data provide strong preclinical justification for prospective sex-stratified trials in prodromal AD, where adrenergic–bioenergetic modulation may yet alter the course of a disease that has resisted intervention for decades.

## MATERIALS AND METHODS

### Animals and terazosin treatment

Fischer 344 TgF344-AD rats co-express human APPswe (K670M/N671L) and PS1ΔE9 under the prion promoter and develop age-dependent, progressive AD-like pathology including amyloid deposition, tau pathology, neuroinflammation, and cognitive decline^45^, making them a well-validated model for AD research. TgF344-AD rats and wild-type (WT) littermates were obtained from the Rat Resource and Research Center (RRRC#699; Columbia, MO) at approximately four weeks of age and housed in pairs under a 12-hour light/dark cycle with food and water available ad libitum. All procedures were approved by the Institutional Animal Care and Use Committee at Hunter College (protocol: MFP-NeuroInflame 1/26) and conducted in accordance with ARRIVE guidelines.

Terazosin hydrochloride (InvivoChem, cat# V1147) was administered ad libitum via Purina 5001 rodent chow (Research Diets Inc., NJ) at a target dose of 0.5 mg/kg bw/day, beginning at 5 months of age and continuing until 11 months. Treatment was initiated during the early phase of pathology, a stage at which therapeutic intervention is predicted to confer maximal benefit. This dose was derived by allometric scaling of the human clinical dose^83^ (1–5 mg/day; 0.083 mg/kg/day for a 60 kg individual) using a rat body surface area conversion factor of 6, and is consistent with prior efficacy studies in APPswe/PS1 transgenic mice^43^. Untreated animals received identical chow without drug. Body weight and food intake were monitored weekly to confirm dosing accuracy (Supplementary Fig 1-4). Four experimental groups were studied: WTNT (7 males, 7 females), WTTR (5 males, 4 females), TGNT (5 males, 6 females), and TGTR (6 males, 5 females).

### Active place avoidance task

Hippocampal-dependent spatial learning and memory were assessed at 11 months using the active place avoidance task (aPAT)^84^. The apparatus comprised a rotating circular platform (1 rev/min) surrounded by four fixed extra maze visual cues. One quadrant was designated as the shock zone; entry for ≥1.5 s triggered a mild foot shock (0.2 mA, repeated every 1.5 s until exit). Animals first underwent a 10-min habituation session (no shock), followed by six 10-min training trials with 10-min inter-trial intervals. Memory retention was assessed 24 h later in a 5-min probe trial with the shock zone deactivated. Position was continuously tracked via overhead camera.

### Tissue collection

At 11 months, rats were anesthetized (ketamine 100 mg/kg / xylazine 10 mg/kg, i.p.) and transcardially perfused with chilled RNase-free PBS. The right hemisphere was micro-dissected, snap-frozen on dry ice, and hippocampal tissue reserved for RNA sequencing. The left hemisphere was immersion-fixed in 4% paraformaldehyde/PBS at 4°C for 48 h, cryoprotected in 30% sucrose/PBS (until the tissue sinks to the bottom of flacon tube containing sucrose), flash-frozen in 2-methylbutane, and stored at −80°C until sectioning.

### Immunohistochemistry and image quantification

Immunohistochemistry (IHC) was performed on coronal dorsal hippocampal sections (Bregma −3.36 to −4.36 mm) as previously described^37^. Sections were imaged on a Zeiss Axio Imager M2 microscope (10× magnification) with Z-stack. Quantitative analysis used custom batch-processing scripts with pixel-intensity thresholding following the rolling-ball method^85^. The positive threshold was defined as mean pixel intensity + k × SD, where k = 4 for Aβ (4G8), and k = 2 for Iba1, NeuN, AT8 (pTau), and NET. Two sections per animal were analyzed and values averaged. Particle size ranges and circularity parameters for each marker are provided in Supplementary Table 37. Primary and secondary antibodies are listed in Supplementary Table 35 and 36.

Microglial morphology was classified by form factor (*FF* = 4π × area/perimeter²): ramified (*FF* < 0.50), reactive (*FF* = 0.50–0.70), and amoeboid (*FF* > 0.70)^47,50,51^.

### RNA sequencing and bioinformatic analysis

Whole left hippocampal RNA was submitted for bulk RNA sequencing to the UCLA Technology Center for Genomics & Bioinformatics (5 untreated and 5 TZ-treated TgF344-AD rats per sex) as previously described^37^. Differential expression was determined using DESeq2 pipeline on R; significantly differentially expressed genes (DEGs) were defined by |log₂ fold change| > 1.0 and *P* < 0.05. Pathway enrichment analysis and data visualization were performed in R Studio. Protein–protein interaction networks were constructed using STRING (v11.5).

### Statistical analysis

All statistical analyses were performed in GraphPad Prism 11 (Boston, MA). Data are presented as mean ± SEM. Three-way ANOVA was used to assess effects of genotype, sex, and treatment; two-way ANOVAs with Sidák’s post hoc tests were applied for pairwise comparisons. Four-way ANOVA for behavior data was performed on JASP software to assess the effects of genotype, sex, trial and treatment. The significance threshold was set at α = 0.05. Image analysis macros are publicly available at https://github.com/SwastikPG209/Pattanashetty.git.

## Data Availability

All data supporting the findings of this study are available from the corresponding author upon request. RNA sequencing datasets will be deposited in the Gene Expression Omnibus (GEO) repository prior to publication, and accession numbers will be included in the final version of the manuscript. Behavioral datasets, imaging data, and custom analysis scripts are likewise available upon request.

## Author contributions

S.G.P., P.S., P.R., and M.F.P., conceived the project and designed the experiments. L.X. performed the computational analysis supporting TZ treatment for AD. S.G.P. performed all experiments and analyzed the data. S.G.P. and M.F.P. wrote the manuscript. P.S. edited the manuscript. All authors approved the manuscript for submission.

## Acknowledgements

Results are based in part on data from AGORA, a platform developed by the NIA-funded AMP-AD consortium supporting Alzheimer’s disease target discovery (doi:10.57718/agora-adknowledgeportal). We thank Dr. Alexander J. Ehrenberg and Dr. Lea T. Grinberg for generating the locus coeruleus differential gene expression dataset in Alzheimer’s disease patient cohorts and for permission to use these data.

## Supplemental material

Supplemental material is available.

