## Supplementary data for "Terazosin drives sex-dependent adrenergic–bioenergetic reprogramming to restore network function in Alzheimer’s disease"

**Supplementary table 1: Male differentially expressed genes (DEGs) in TgF344-AD rats (WTNT vs TGNT)**

| | MALE TGNT DEGs | |
|---|---|---|
| Gene symbol | P value | Log2 Fold change |
| krt34 | 0.000916397 | -4.973800857 |
| syndig1l | 0.001008468 | -4.968007353 |
| lhx8 | 0.0148 | -4.957635323 |
| trpv5 | 0.026023479 | -4.956533729 |
| htr1d | 0.017287605 | -4.811582687 |
| isl1 | 0.0483 | -4.767495392 |
| loc689927 | 0.0136 | -4.63927781 |
| nlrp4 | 0.020187616 | -4.614233636 |
| fam71f1 | 0.010095178 | -4.389255317 |
| fgf20 | 0.009784828 | -4.367237596 |
| gbx2 | 0.007593022 | -4.309980724 |
| adora2a | 0.000152568 | -4.126134252 |
| tbc1d10c | 0.012268062 | -3.425234445 |
| drd2 | 0.022208633 | -3.395031531 |
| gsc2 | 0.0416 | -3.136141861 |
| rgs9 | 0.001083563 | -3.09966529 |
| serpinc1 | 0.0196 | -3.077681541 |
| kcnmb3 | 0.0476 | -3.076088084 |
| nepn | 0.0073 | -3.02922065 |
| krt75 | 0.009856344 | -2.843618063 |
| tns4 | 0.0481 | -2.720391877 |
| tac1 | 0.033463804 | -2.71967758 |
| sct | 0.0091 | -2.705163265 |
| olr1653 | 0.0308 | -2.701400547 |
| s100a5 | 0.0142 | -2.699338058 |
| hepacam2 | 0.0046 | -2.684683438 |

| | | |
|---|---|---|
| cartpt | 0.049292638 | -2.679429914 |
| retn | 0.045 | -2.622065951 |
| gbx1 | 0.0319 | -2.617963163 |
| prkag3 | 0.0418 | -2.609560586 |
| pcp4l1 | 0.00956063 | -2.601415547 |
| gpr6 | 0.044791117 | -2.506889471 |
| nexn | 0.049901329 | -2.46306657 |
| gpr88 | 0.0128 | -2.393881258 |
| krt16 | 0.013384094 | -2.389008039 |
| krt86 | 0.002049727 | -2.329688033 |
| rgd1561870 | 0.0082 | -2.32705738 |
| sh3rf2 | 0.0283 | -2.309384189 |
| nmu | 0.0458 | -2.299239819 |
| drd1 | 0.009924751 | -2.25950207 |
| mxd3 | 0.0208 | -2.253662931 |
| dmkn | 0.0163 | -2.244888394 |
| penk | 0.00806351 | -2.183835665 |
| loc100361092 | 0.0398 | -2.147670655 |
| tgm3 | 0.0376 | -2.120127088 |
| scn4b | 0.004872473 | -2.04769346 |
| rasd2 | 9.11133E-05 | -1.984490805 |
| slc18a3 | 0.0042 | -1.976132608 |
| sv2c | 0.00499171 | -1.966645378 |
| gng7 | 0.000296281 | -1.966142158 |
| itk | 0.0368 | -1.956164347 |
| hist1h2aa | 0.0391 | -1.909584155 |
| pof1b | 0.044969038 | -1.887918024 |
| neu2 | 0.0259 | -1.879959339 |
| unc13c | 0.0145 | -1.81587446 |
| adra2b | 0.0079 | -1.808609059 |
| foxp2 | 0.0334 | -1.793291449 |
| trat1 | 0.0368 | -1.785824591 |
| slc35d3 | 0.048168182 | -1.784838266 |
| fgf19 | 0.0147 | -1.766715264 |
| ppp1r1b | 0.008514624 | -1.758865551 |
| nr5a1 | 0.0219 | -1.751573637 |
| krt71 | 0.005 | -1.744586941 |
| ces2 | 0.0456 | -1.735457396 |
| spink5 | 0.0275 | -1.730225219 |

| | | |
|---|---|---|
| rgd1559714 | 0.0345 | -1.661617214 |
| asic4 | 0.0084 | -1.6412625 |
| rasgrp2 | 0.0126 | -1.627148613 |
| adcy5 | 0.012163827 | -1.625562269 |
| rarb | 0.024246222 | -1.603289398 |
| vwce | 0.0157 | -1.600416378 |
| rt1-ce3 | 0.035548534 | -1.594258813 |
| pdzk1ip1 | 0.0271 | -1.547562303 |
| tnni3 | 0.017 | -1.543885052 |
| troap | 0.0287 | -1.54265574 |
| pde10a | 0.000323221 | -1.540748632 |
| jph2 | 0.0138 | -1.535695067 |
| col7a1 | 0.006 | -1.534140059 |
| clec4e | 0.0329 | -1.518085757 |
| cd200r1 | 0.0041 | -1.512778896 |
| pitx2 | 0.0317 | -1.507056246 |
| il2rg | 0.0306 | -1.496099366 |
| slamf6 | 0.0424 | -1.494245958 |
| pou4f3 | 0.0117 | -1.487632903 |
| klk7 | 0.0321 | -1.479561525 |
| ccnb1ip1 | 0.0192 | -1.449842319 |
| chrdl2 | 0.0049 | -1.443149424 |
| gpr153 | 0.0168 | -1.442425292 |
| pdzd2 | 0.038683392 | -1.432781942 |
| pde7b | 0.039127861 | -1.415615466 |
| rgd1308065 | 0.0469 | -1.411205704 |
| tssk3 | 0.0418 | -1.398802893 |
| rgd1561778 | 0.017 | -1.382200972 |
| kb15 | 0.0272 | -1.374017979 |
| runx1 | 0.0273 | -1.33799343 |
| grm4 | 0.0284 | -1.337088463 |
| hist2h3c2 | 0.0462 | -1.326817845 |
| meis2 | 0.0129 | -1.306436438 |
| slc10a4 | 0.0098 | -1.304159453 |
| kcna5 | 0.0133 | -1.302494436 |
| actn3 | 0.0408 | -1.298795855 |
| itpr3 | 0.0087 | -1.298622209 |
| trh | 0.0225 | -1.296535202 |
| pde1b | 0.056265413 | -1.295849745 |

| | | |
|---|---|---|
| cdhr1 | 0.0062 | -1.285230396 |
| ncf2 | 0.0314 | -1.283622508 |
| aldoart2 | 0.0401 | -1.282786677 |
| pbx3 | 0.0206 | -1.276686832 |
| lrrk2 | 0.034591949 | -1.276381766 |
| prkch | 0.0181 | -1.27074277 |
| tpm2 | 0.0269 | -1.268862142 |
| ptpn5 | 0.036111164 | -1.255217086 |
| crabp1 | 0.0358 | -1.2526808 |
| camp | 0.0386 | -1.250914595 |
| arpp21 | 0.0048 | -1.230201939 |
| cdk1 | 0.0337 | -1.208218991 |
| dlx5 | 0.0344 | -1.201843969 |
| kcnh4 | 0.0383 | -1.201753218 |
| bcmo1 | 0.0159 | -1.19979729 |
| ecel1 | 0.0105 | -1.199505229 |
| pcp4 | 0.0413 | -1.195075626 |
| irx6 | 0.0167 | -1.190053647 |
| htr6 | 0.0127 | -1.188835585 |
| ltbp2 | 0.0194 | -1.180699258 |
| hist2h4 | 0.0448 | -1.17720063 |
| ngef | 0.0086 | -1.17564368 |
| gnal | 0.0211 | -1.172722623 |
| rgd1561102 | 0.0421 | -1.171124415 |
| egfr | 0.0339 | -1.167310238 |
| krt1 | 0.0064 | -1.158631491 |
| unc13d | 0.0415 | -1.156044439 |
| art3 | 0.0056 | -1.14778209 |
| cmtm8 | 0.0221 | -1.145933045 |
| insrr | 0.0294 | -1.145310763 |
| mme | 0.0399 | -1.143364471 |
| cenpf | 0.0129 | -1.140344279 |
| kcnk2 | 0.0261 | -1.136276643 |
| lysmd3 | 0.0291 | -1.135933955 |
| cpne5 | 0.0183 | -1.132506813 |
| kremen1 | 0.0204 | -1.128784443 |
| sh2b2 | 0.0099 | -1.124523613 |
| vgll2 | 0.0419 | -1.119732676 |
| pdp1 | 0.0369 | -1.117088639 |

| | | |
|---|---|---|
| xcl1 | 0.0219 | -1.115847289 |
| rin3 | 0.0363 | -1.114631496 |
| iqgap3 | 0.0348 | -1.112889619 |
| cmah | 0.0115 | -1.108072252 |
| leap2 | 0.0467 | -1.10803446 |
| atp6ap1l | 0.0119 | -1.107871988 |
| pcp2 | 0.0128 | -1.10744689 |
| prdm1 | 0.0136 | -1.105187906 |
| s100g | 0.0322 | -1.105138521 |
| itga5 | 0.0125 | -1.104367484 |
| col5a2 | 0.0162 | -1.096384398 |
| veph1 | 0.0415 | -1.095440732 |
| loc102548399 | 0.0176 | -1.093551994 |
| slc5a12 | 0.0253 | -1.092686221 |
| dclk3 | 0.0081 | -1.092286127 |
| prkar2b | 0.0302 | -1.091632802 |
| rgd1305627 | 0.0089 | -1.087654544 |
| rgd1565655 | 0.0069 | -1.081046285 |
| cobl | 0.0454 | -1.075733588 |
| rgd1306625 | 0.0436 | -1.07373809 |
| gpr52 | 0.0332 | -1.068883393 |
| ube2l6 | 0.0195 | -1.067415769 |
| adh7 | 0.0338 | -1.064227614 |
| ino80d | 0.0352 | -1.062096447 |
| bmp2 | 0.0071 | -1.057524859 |
| bcl2l11 | 0.0149 | -1.056976378 |
| gucy1a3 | 0.0098 | -1.053965912 |
| loc689713 | 0.0143 | -1.053027448 |
| hcls1 | 0.0184 | -1.049279306 |
| ret | 0.0171 | -1.042807217 |
| plek | 0.0046 | -1.040677375 |
| fam222a | 0.0369 | -1.038916169 |
| c3ar1 | 0.0194 | -1.035632931 |
| prima1 | 0.0309 | -1.026522383 |
| pnpla1 | 0.0042 | -1.024547354 |
| rrh | 0.0427 | -1.020654763 |
| capg | 0.0459 | -1.02023542 |
| rmrp | 0.0165 | -1.01378135 |
| myo1f | 0.0108 | -1.004818154 |

| | | |
|---|---|---|
| fyb | 0.0078 | -1.001091052 |
| olr1869 | 0.0491 | -1 |
| mirlet7d | 0.0452 | -1 |
| cryge | 0.0412 | -1 |
| ugt1a9 | 0.0387 | -1 |
| ugt1a5 | 0.0365 | -1 |
| vom1r16 | 0.0345 | -1 |
| olr176 | 0.0314 | -1 |
| mir421 | 0.0289 | -1 |
| testin | 0.0276 | -1 |
| magea4 | 0.0237 | -1 |
| mirlet7c-2 | 0.0224 | -1 |
| reg3a | 0.0211 | -1 |
| cb707485 | 0.0198 | -1 |
| gykl1 | 0.0173 | -1 |
| sell | 0.0156 | -1 |
| spetex-2a | 0.0123 | -1 |
| trim40 | 0.0112 | -1 |
| dcm5 | 0.0095 | -1 |
| ifna4 | 0.0087 | -1 |
| mir3572 | 0.0061 | -1 |
| il22ra1 | 0.0049 | -1 |
| olr1667 | 0.0034 | -1 |
| il20ra | 0.0182 | 1.000663382 |
| myl6 | 0.0443 | 1.002057439 |
| ajuba | 0.0139 | 1.003373268 |
| cdca7 | 0.0025 | 1.005290231 |
| krt85 | 0.0267 | 1.00604893 |
| slc22a8 | 0.0476 | 1.007471174 |
| exo1 | 0.0398 | 1.008592605 |
| fbxo39 | 0.0204 | 1.010903093 |
| vav1 | 0.0482 | 1.014349944 |
| rab38 | 0.0057 | 1.015467857 |
| olr200 | 0.0351 | 1.015539418 |
| stk32b | 0.0091 | 1.017841031 |
| h19 | 0.0191 | 1.021115549 |
| col18a1 | 0.0405 | 1.024069057 |
| timd2 | 0.0187 | 1.025478372 |
| mapk4 | 0.0421 | 1.026386208 |

| | | |
|---|---|---|
| hist1h2bo | 0.0281 | 1.026458795 |
| arhgap30 | 0.0132 | 1.03067264 |
| rgd1303271 | 0.0306 | 1.035671746 |
| fam83a | 0.0249 | 1.038839011 |
| pappa | 0.0083 | 1.040489859 |
| slc4a11 | 0.0393 | 1.041858165 |
| spock3 | 0.0143 | 1.049825069 |
| otx1 | 0.0349 | 1.050231093 |
| fbp1 | 0.0372 | 1.05342245 |
| loc287004 | 0.0289 | 1.053962739 |
| ccl9 | 0.0201 | 1.061243401 |
| lipogenin | 0.0412 | 1.066290292 |
| six3 | 0.0341 | 1.071367825 |
| kctd8 | 0.0158 | 1.072434889 |
| entpd3 | 0.0246 | 1.073507103 |
| stra6 | 0.0227 | 1.075807452 |
| akap12 | 0.0371 | 1.078729891 |
| rn28s | 0.0172 | 1.079148613 |
| kif23 | 0.0298 | 1.08107081 |
| naglt1 | 0.0282 | 1.084042237 |
| tnxa-ps1 | 0.0058 | 1.087964046 |
| hist1h2ak | 0.0264 | 1.089418365 |
| kcnn4 | 0.0213 | 1.090977211 |
| rdh7 | 0.0063 | 1.091036149 |
| msc | 0.0381 | 1.091631861 |
| arl11 | 0.0308 | 1.091729198 |
| tnnt2 | 0.0261 | 1.098396488 |
| rgd1561958 | 0.0121 | 1.099780719 |
| ttc26 | 0.0153 | 1.106087424 |
| serpine1 | 0.0209 | 1.110511492 |
| slc5a7 | 0.0316 | 1.115250524 |
| loc100363193 | 0.0394 | 1.115427968 |
| clec2d | 0.0054 | 1.116111981 |
| dmrtc1c1 | 0.0151 | 1.121505659 |
| oas1h | 0.0207 | 1.126429487 |
| rrm2 | 0.0486 | 1.127170079 |
| ccdc166 | 0.0472 | 1.133566835 |
| crb1 | 0.0092 | 1.135689591 |
| tinagl1 | 0.0415 | 1.140794335 |

| | | |
|---|---|---|
| mrgprf | 0.0258 | 1.142404517 |
| tmem170a | 0.0379 | 1.147397317 |
| gulp1 | 0.0138 | 1.152345735 |
| pou1f1 | 0.0473 | 1.152373848 |
| slc31a1 | 0.0481 | 1.153964121 |
| igsf9 | 0.0216 | 1.159607334 |
| slc16a3 | 0.0392 | 1.163500392 |
| slc5a5 | 0.0043 | 1.164089264 |
| rbp4 | 0.0087 | 1.166148154 |
| has1 | 0.0376 | 1.182505713 |
| gapdh-ps1 | 0.0051 | 1.190962959 |
| cmtm1 | 0.0075 | 1.200593784 |
| txk | 0.0274 | 1.201014694 |
| arl4d | 0.0316 | 1.208894849 |
| zic2 | 0.018 | 1.209915179 |
| lif | 0.0335 | 1.210785536 |
| rnf152 | 0.0452 | 1.233077708 |
| adcy7 | 0.0291 | 1.233317794 |
| mael | 0.0072 | 1.235171355 |
| cgnl1 | 0.0315 | 1.236394155 |
| cldn22 | 0.0187 | 1.239588745 |
| hp | 0.0076 | 1.239633032 |
| ldhal6b | 0.0378 | 1.243822747 |
| ptprq | 0.0461 | 1.247130074 |
| mgp | 0.045 | 1.262228305 |
| top2a | 0.0473 | 1.262924689 |
| mcoln3 | 0.0465 | 1.272718969 |
| def6 | 0.0071 | 1.286880751 |
| odf3l2 | 0.0096 | 1.287729438 |
| ugt1a6 | 0.0089 | 1.287871904 |
| lsp1 | 0.0267 | 1.288137027 |
| il18bp | 0.0347 | 1.293780649 |
| mdfic | 0.0065 | 1.301760581 |
| atp8b1 | 0.049 | 1.303271973 |
| trpm3 | 0.0067 | 1.303981504 |
| krt42 | 0.0068 | 1.304191812 |
| bst1 | 0.0202 | 1.305370199 |
| loxl2 | 0.0218 | 1.312737295 |
| ddc8 | 0.0312 | 1.313557646 |

| | | |
|---|---|---|
| avpr2 | 0.0114 | 1.316399165 |
| gfap | 0.037272321 | 1.321852452 |
| spp1 | 0.0297 | 1.331318674 |
| reck | 0.0189 | 1.336882327 |
| fhod3 | 0.0328 | 1.339908167 |
| cdr2 | 0.033 | 1.344462671 |
| tmem196 | 0.0203 | 1.351834359 |
| apobec1 | 0.042 | 1.354225105 |
| fscn2 | 0.02 | 1.354790043 |
| mdk | 0.0056 | 1.355200752 |
| rgd1309028 | 0.0135 | 1.358807627 |
| stat4 | 0.0437 | 1.362932961 |
| sp8 | 0.0276 | 1.363461807 |
| lhfpl1 | 0.0286 | 1.36411163 |
| mcm10 | 0.0241 | 1.365384759 |
| ccdc60 | 0.001112422 | 1.372642254 |
| mospd1 | 0.0349 | 1.37326538 |
| fcgr2b | 0.043608436 | 1.379440562 |
| iqub | 0.0256 | 1.38058583 |
| bcl3 | 0.0112 | 1.387756646 |
| tead2 | 0.0466 | 1.38871077 |
| tshr | 0.0261 | 1.390323899 |
| gypc | 0.0289 | 1.393972956 |
| loxl4 | 0.0257 | 1.402386545 |
| sgcg | 0.0151 | 1.410580078 |
| kiss1 | 0.0453 | 1.420393072 |
| rab11fip1 | 0.0075 | 1.421259041 |
| sim1 | 0.028 | 1.421370569 |
| mvd | 7.49071E-05 | 1.454057763 |
| zic3 | 0.0198 | 1.457597599 |
| glb1l | 0.0097 | 1.459328978 |
| pabpc2 | 0.0293 | 1.464708025 |
| slc16a2 | 0.0061 | 1.465532751 |
| mmp2 | 0.0174 | 1.466410256 |
| elovl7 | 0.0344 | 1.468976749 |
| mpz | 0.0131 | 1.477249676 |
| ect2 | 0.0149 | 1.478396123 |
| evpl | 0.0153 | 1.482850442 |
| synpo2 | 0.0295 | 1.483575681 |

| | | |
|---|---|---|
| npr3 | 0.011 | 1.485478648 |
| slc12a7 | 0.0468 | 1.48616303 |
| f13a1 | 0.0279 | 1.487390527 |
| col5a1 | 0.019 | 1.488981934 |
| cd22 | 0.0452 | 1.494612158 |
| nags | 0.0154 | 1.497725194 |
| a2m | 0.0045 | 1.505123869 |
| irx3 | 0.0331 | 1.512826579 |
| klra5 | 0.0228 | 1.515801991 |
| slc4a2 | 0.0165 | 1.519730826 |
| chad | 0.0394 | 1.524467441 |
| rt1-ce15 | 0.0363 | 1.529608175 |
| st6galnac2 | 0.0214 | 1.535595256 |
| prelp | 0.0423 | 1.535656729 |
| tbx6 | 0.0116 | 1.536000744 |
| efhc1 | 0.0254 | 1.537349764 |
| gck | 0.0378 | 1.546632395 |
| gucy2f | 0.0175 | 1.555700191 |
| phactr2 | 0.0337 | 1.569659058 |
| dab2 | 0.0044 | 1.60460013 |
| wdr16 | 0.0487 | 1.606812044 |
| clec5a | 0.0212 | 1.62012652 |
| coch | 0.0119 | 1.621451877 |
| ccl4 | 0.0104 | 1.631619025 |
| hes3 | 0.0268 | 1.639668797 |
| tex15 | 0.0121 | 1.64222963 |
| fam109b | 0.0132 | 1.649863998 |
| slc2a12 | 0.0094 | 1.666072739 |
| melk | 0.0396 | 1.67604954 |
| gpnmb | 7.44028E-07 | 1.680622709 |
| pld5 | 0.0301 | 1.687528254 |
| brs3 | 0.0307 | 1.688304451 |
| zim1 | 0.0311 | 1.693995305 |
| epha1 | 0.0146 | 1.709072901 |
| dnali1 | 0.041424929 | 1.710465443 |
| tc2n | 0.0169 | 1.735175289 |
| gpx2 | 0.0457 | 1.75390144 |
| car14 | 0.0426 | 1.755677999 |
| naa11 | 0.0157 | 1.765303473 |

| | | |
|---|---|---|
| aldh1a3 | 0.0073 | 1.787483382 |
| foxc2 | 0.0093 | 1.788835922 |
| hist1h1b | 0.0134 | 1.79661214 |
| baiap2l1 | 0.026 | 1.80123678 |
| mcoln2 | 0.0124 | 1.850164548 |
| zic1 | 0.0089 | 1.855215976 |
| dpp4 | 0.0184 | 1.862576852 |
| pde6b | 0.0217 | 1.862868067 |
| enkur | 0.008366262 | 1.866176985 |
| txndc2 | 0.031 | 1.867141199 |
| cyp11b3 | 0.0089 | 1.871141995 |
| itgb1bp2 | 0.0389 | 1.876373179 |
| igfals | 0.0441 | 1.879621121 |
| cyp11a1 | 0.0484 | 1.881553434 |
| cd68 | 0.004035689 | 1.892419024 |
| cd69 | 0.0462 | 1.933413887 |
| prlr | 0.0202 | 1.938206083 |
| cox8b | 0.0411 | 1.938719315 |
| scara5 | 0.0054 | 1.946854599 |
| loc691083 | 0.0436 | 1.962402999 |
| lrrc52 | 0.0147 | 1.977595764 |
| sulf1 | 0.0057 | 2.006084353 |
| ropn1l | 0.057476096 | 2.020496107 |
| rgd1560608 | 0.0374 | 2.022123718 |
| epn3 | 0.039044776 | 2.028123123 |
| llgl2 | 0.0346 | 2.031111187 |
| adamts19 | 0.020727371 | 2.034565223 |
| nkx2-1 | 0.036313473 | 2.059541241 |
| runx3 | 0.0186 | 2.064434637 |
| fmo2 | 0.050166318 | 2.070271669 |
| fam46a | 0.0335 | 2.071856143 |
| olr387 | 0.0052 | 2.079548754 |
| mybl2 | 0.0309 | 2.097045814 |
| slc23a3 | 0.0427 | 2.097859123 |
| dnai2 | 0.013139008 | 2.10662455 |
| ccl3 | 0.047536962 | 2.111974277 |
| atp4a | 0.0192 | 2.115561694 |
| bhmt | 0.0318 | 2.118804035 |
| lepr | 0.0464 | 2.126573554 |

| | | |
|---|---|---|
| adamtsl4 | 0.035195948 | 2.127597377 |
| sfrp1 | 0.0117 | 2.141463152 |
| col3a1 | 0.0453 | 2.145538134 |
| rhod | 0.039 | 2.155649386 |
| cxcl13 | 0.0122 | 2.201709668 |
| app | 5.80102E-10 | 2.217520176 |
| zmynd10 | 0.003970383 | 2.236976757 |
| psen1 | 3.41957E-13 | 2.243999669 |
| aox3 | 0.0265 | 2.248629722 |
| igfbp2 | 0.0068 | 2.285258887 |
| frem1 | 0.026563554 | 2.288536893 |
| tnn | 0.0065 | 2.293025012 |
| slc16a12 | 0.055675932 | 2.335047944 |
| cd72 | 0.03399141 | 2.335726081 |
| col1a2 | 0.0386 | 2.345389371 |
| best3 | 0.0332 | 2.348210946 |
| otc | 0.0159 | 2.352940962 |
| cpn1 | 0.0124 | 2.364311796 |
| klhl1 | 0.0077 | 2.364312086 |
| rgd1562658 | 0.017773207 | 2.366964481 |
| foxb1 | 0.0118 | 2.389371368 |
| rgd1563104 | 0.0264 | 2.463831749 |
| gfi1b | 0.0049 | 2.464260307 |
| chrna9 | 0.0303 | 2.46426391 |
| acot12 | 0.0479 | 2.46441498 |
| fam187a | 0.056194221 | 2.471374811 |
| cdkn1c | 0.029628046 | 2.476568793 |
| ido1 | 0.0095 | 2.493165054 |
| gja6 | 0.0365 | 2.493535003 |
| ogn | 0.0204 | 2.49785142 |
| dazl | 0.0342 | 2.518520549 |
| vom2r60 | 0.0422 | 2.528975064 |
| spag11c | 0.0063 | 2.529168714 |
| tcp10b | 0.027067082 | 2.529199804 |
| crygf | 0.02706524 | 2.529430448 |
| vwa5b1 | 0.003487636 | 2.59313008 |
| crb3 | 0.050611958 | 2.614477383 |
| kcnv2 | 0.0286 | 2.692314397 |
| msx1 | 0.013457742 | 2.740573939 |

| | | |
|---|---|---|
| klk1 | 0.034206394 | 2.778877791 |
| lhx5 | 0.036937723 | 2.867980284 |
| scgb1c1 | 0.056019202 | 2.869483736 |
| cyp2c24 | 0.0096 | 2.878811483 |
| spag8 | 0.017739315 | 2.943796665 |
| smim22 | 0.032572505 | 2.963146455 |
| fgf1 | 0.033365314 | 2.984185077 |
| sult1c3 | 0.0281 | 3.004389426 |
| foxj1 | 0.001432266 | 3.115277907 |
| lrr1 | 0.0362 | 3.137818099 |
| folr1 | 0.002164568 | 3.165649544 |
| lilrb4 | 0.003350786 | 3.178212011 |
| slc13a4 | 0.04962834 | 3.193350298 |
| igf2 | 0.0397 | 3.203952103 |
| rgd1559696 | 0.0382 | 3.206210824 |
| efcab1 | 0.003612565 | 3.232413745 |
| samd3 | 0.0219 | 3.240020555 |
| glycam1 | 0.002216144 | 3.300954039 |
| lag3 | 0.00060893 | 3.307997554 |
| kcnj13 | 0.007013337 | 3.330162273 |
| efhb | 0.000621129 | 3.350924256 |
| rgd1561161 | 0.022887286 | 3.355280398 |
| lhx1 | 0.0053 | 3.426710321 |
| h1foo | 0.0425 | 3.437164553 |
| ifltd1 | 0.014132002 | 3.459715463 |
| igfbpl1 | 0.02675047 | 3.559941613 |
| hnf1b | 0.0113 | 3.613931092 |
| lmx1a | 0.001639121 | 3.629151826 |
| uncx | 0.041073353 | 3.665696962 |
| tmem184a | 0.053534936 | 3.687097874 |
| mfrp | 0.002516724 | 3.695799485 |
| slc16a8 | 0.00548204 | 3.711568418 |
| car13 | 0.000806284 | 3.723997507 |
| sytl3 | 0.030311178 | 3.729626803 |
| scube3 | 0.025256551 | 3.732218767 |
| cd74 | 2.34734E-17 | 3.745997152 |
| rgd1561795 | 0.055950812 | 3.791570516 |
| capsl | 0.009844163 | 3.795120655 |
| clic6 | 0.004214637 | 3.805656924 |

| | | |
|---|---|---|
| lyzl6 | 0.034627499 | 3.87882065 |
| ccdc114 | 0.000852527 | 3.881815653 |
| cdh3 | 0.047486723 | 3.884351775 |
| calcr | 0.053423592 | 3.930035517 |
| loc654482 | 0.001732398 | 3.946278001 |
| loc100910620 | 0.031640709 | 3.967670838 |
| clul1 | 0.0292 | 3.969100358 |
| fgg | 0.0187 | 3.989893255 |
| il13ra2 | 0.015711516 | 4.002530666 |
| slc4a5 | 0.001444953 | 4.002617431 |
| pon3 | 0.007405018 | 4.003595894 |
| wdr63 | 0.000446265 | 4.007807852 |
| akp3 | 0.03062923 | 4.04053697 |
| rt1-da | 7.7854E-13 | 4.067130789 |
| c1qtnf3 | 0.015625163 | 4.070634555 |
| cldn2 | 0.00581944 | 4.083588411 |
| six1 | 0.0139 | 4.116297131 |
| crygs | 0.006302575 | 4.20672846 |
| trpv4 | 0.033213686 | 4.208908014 |
| pon1 | 0.054452445 | 4.227626766 |
| mc3r | 0.05316734 | 4.288560138 |
| slc22a3 | 0.011481345 | 4.292287065 |
| upk1b | 0.0367 | 4.318392191 |
| itpripl1 | 0.023543905 | 4.359837184 |
| lgals5 | 0.026945043 | 4.366038525 |
| rbm47 | 0.018625085 | 4.368644904 |
| fam160a1 | 0.002409006 | 4.379774969 |
| slc7a12 | 0.0193 | 4.399630707 |
| slpil2 | 0.029 | 4.406147384 |
| pax8 | 0.0072 | 4.409217217 |
| rt1-db1 | 7.8347E-09 | 4.415096438 |
| vom1r88 | 0.0461 | 4.430283383 |
| sostdc1 | 0.007943348 | 4.439728235 |
| mlf1 | 0.000346501 | 4.442961563 |
| slco1a5 | 0.00257336 | 4.448383876 |
| ttr | 0.007349162 | 4.459996464 |
| aqp1 | 0.010459209 | 4.463057265 |
| steap1 | 0.002170313 | 4.467267371 |
| tcf21 | 0.008608217 | 4.498096789 |

| | | |
|---|---|---|
| bhmt2 | 0.001731871 | 4.58156154 |
| krt18 | 0.000770578 | 4.618409619 |
| lect1 | 0.00123633 | 4.636553192 |
| rt1-ba | 2.48778E-12 | 4.640354357 |
| kcnq1 | 0.008970116 | 4.658734252 |
| ccdc40 | 0.001171888 | 4.660969734 |
| loc171161 | 0.006309592 | 4.672480925 |
| tmem72 | 0.006442168 | 4.709753843 |
| rgd1563692 | 0.011897944 | 4.714890955 |
| kl | 0.002225866 | 4.720538088 |
| zscan10 | 0.006162267 | 4.725199321 |
| agtr2 | 0.025651931 | 4.729211163 |
| lenep | 0.036635908 | 4.764256588 |
| hnf4a | 0.05866896 | 4.778330915 |
| gdf7 | 0.009486827 | 4.794744831 |
| mpzl2 | 0.01500919 | 4.810859358 |
| sbk2 | 0.0285 | 4.818240317 |
| inmt | 0.014831821 | 4.823613929 |
| ace | 0.012327405 | 4.834893537 |
| pih1d3 | 0.005402847 | 4.84807788 |
| f5 | 0.000755156 | 4.8651459 |
| enpp2 | 0.007183236 | 4.889461154 |
| ces1d | 0.000727855 | 4.897267355 |
| lrrc18 | 0.007314711 | 4.911216836 |
| otx2 | 0.001370258 | 4.923277884 |
| pla2g5 | 0.009640822 | 4.932686288 |
| tmem27 | 0.034375119 | 4.945275648 |
| mak | 0.001973619 | 4.951423764 |
| itgb6 | 0.017800726 | 5.003553004 |
| kcna10 | 0.0215 | 5.00919122 |
| ms4a14 | 0.0152 | 5.00922868 |
| slc14a2 | 0.045688242 | 5.013552037 |
| fetub | 0.002704325 | 5.014892896 |
| adipoq | 0.043759023 | 5.030519337 |
| hs3st3a1 | 0.0182 | 5.04418223 |
| corin | 0.004998593 | 5.046299301 |
| cdx4 | 0.0419 | 5.053446666 |
| iqcg | 0.001105116 | 5.064436259 |
| abca4 | 0.005849554 | 5.07792078 |

| | | |
|---|---|---|
| kcne2 | 0.008019575 | 5.083984347 |
| klk4 | 0.058885525 | 5.08900984 |

**Supplementary table 2: Female differentially expressed genes (DEGs) in TgF344-AD rats (WTNT vs TGNT)**

| | **FEMALE TGNT DEGs** | |
|---|---|---|
| Gene_symbol | PValue | log2 Foldchange |
| hepacam2 | 0.049060237 | -5.834685796 |
| cyp11a1 | 0.049214931 | -5.824247952 |
| slc18a3 | 0.002255172 | -5.824203248 |
| crygf | 0.049403008 | -5.810560982 |
| krt85 | 0.017212358 | -5.685844082 |
| neu2 | 0.002522829 | -5.605601086 |
| mcoln3 | 0.010296603 | -5.577033408 |
| mpzl2 | 0.005802334 | -5.539978101 |
| slc13a4 | 0.002144718 | -5.490627215 |
| clec4e | 0.032444739 | -5.489097408 |
| igfbpl1 | 0.000965676 | -5.457367069 |
| drd1 | 0.000186619 | -5.414083386 |
| smim22 | 0.033166046 | -5.364985784 |
| kcnv2 | 0.009257784 | -5.336256502 |
| penk | 0.000321602 | -5.287294927 |
| pih1d3 | 0.010252744 | -5.202821901 |
| rbm47 | 0.00518788 | -5.150822803 |
| tac1 | 0.001053216 | -5.089164182 |
| lrrc18 | 0.016439184 | -5.063238977 |
| msc | 0.037442223 | -4.903395486 |
| cldn22 | 0.048261139 | -4.89478665 |
| prlr | 0.00272059 | -4.871383834 |
| cyp2c24 | 0.047812449 | -4.833354032 |
| clec2d | 0.039008845 | -4.816360839 |
| ecel1 | 0.0000849 | -4.812014203 |
| gdf7 | 0.046534504 | -4.7908073 |
| nexn | 0.001884323 | -4.783755207 |
| scn4b | 0.000545622 | -4.72959896 |
| klk7 | 0.021544529 | -4.675438847 |
| cyp11b3 | 0.026282238 | -4.667821248 |
| scube3 | 0.007993893 | -4.61861351 |

| | | |
|---|---|---|
| sulf1 | 0.000657947 | -4.604583154 |
| fgf1 | 0.00337452 | -4.570126423 |
| rrh | 0.047257827 | -4.470617276 |
| itk | 0.005853163 | -4.382501689 |
| cartpt | 0.00373508 | -4.315002407 |
| stat4 | 0.042517003 | -4.231727223 |
| corin | 0.035625146 | -4.193762512 |
| slc16a8 | 0.011612302 | -4.134922945 |
| mak | 0.020801948 | -4.132623087 |
| ppp1r1b | 0.000803423 | -4.027044203 |
| krt71 | 0.001000619 | -4.006988628 |
| igfbp2 | 0.007561379 | -3.935859369 |
| rgd1561161 | 0.006230402 | -3.927838676 |
| efhb | 0.033644362 | -3.895518425 |
| sytl3 | 0.045659337 | -3.85406966 |
| gpr52 | 0.005249261 | -3.835201717 |
| chrdl2 | 0.04458752 | -3.832350969 |
| lrr1 | 0.048986635 | -3.795095298 |
| pou1f1 | 0.046240214 | -3.7907119 |
| atp4a | 0.015659191 | -3.742734472 |
| zic1 | 0.001091912 | -3.701068968 |
| car13 | 0.02703681 | -3.674617614 |
| ccdc60 | 0.043738913 | -3.64110608 |
| hist1h1b | 0.019374741 | -3.637862376 |
| sell | 0.048352213 | -3.615998437 |
| tgm3 | 0.043486401 | -3.612257728 |
| atp6ap1l | 0.003150546 | -3.556817438 |
| cdk1 | 0.010966934 | -3.556072749 |
| cpn1 | 0.028969574 | -3.543273706 |
| crb3 | 0.017576188 | -3.540000932 |
| slc35d3 | 0.002512311 | -3.518450022 |
| fam160a1 | 0.005250986 | -3.518028915 |
| asic4 | 0.002474923 | -3.468336241 |
| loc100361092 | 0.021489549 | -3.434999393 |
| bhmt | 0.033608916 | -3.345221852 |
| otc | 0.02816296 | -3.322010251 |
| ogn | 0.001882511 | -3.318508298 |
| ccl9 | 0.003621062 | -3.305348716 |
| trh | 0.002368494 | -3.299983763 |

| | | |
|---|---|---|
| lepr | 0.002972585 | -3.290420631 |
| txk | 0.032118762 | -3.27941112 |
| rgd1560608 | 0.019922918 | -3.264762765 |
| mael | 0.031720345 | -3.260393193 |
| atp5po | 0.00342 | -3.2321 |
| ifltd1 | 0.037507774 | -3.191825188 |
| krt75 | 0.031299852 | -3.099035898 |
| exo1 | 0.042481836 | -3.062937124 |
| hnf1b | 0.044796992 | -3.004528341 |
| adamtsl4 | 0.013448284 | -2.99665116 |
| coch | 0.012771639 | -2.978140901 |
| loc654482 | 0.010891784 | -2.947801043 |
| krt18 | 0.027788286 | -2.94389384 |
| slc16a12 | 0.008544782 | -2.936287068 |
| mcm10 | 0.040096861 | -2.931450868 |
| pcp4l1 | 0.003539121 | -2.911647857 |
| lhfpl1 | 0.02982663 | -2.882665773 |
| rasgrp2 | 0.015361745 | -2.878942234 |
| frem1 | 0.00565816 | -2.862250306 |
| mybl2 | 0.016725762 | -2.853192278 |
| iqgap3 | 0.008205694 | -2.84628606 |
| krt1 | 0.049677256 | -2.84224286 |
| top2a | 0.022904948 | -2.83110044 |
| slc10a4 | 0.020120982 | -2.825683286 |
| cenpf | 0.026854724 | -2.811674037 |
| rarb | 0.006467469 | -2.793156514 |
| slamf6 | 0.04926346 | -2.752595137 |
| fam46a | 0.015496389 | -2.749046838 |
| rasd2 | 0.002247495 | -2.744676039 |
| mdfic | 0.00765124 | -2.744101367 |
| adra2b | 0.011099462 | -2.743615181 |
| best3 | 0.039708766 | -2.732943255 |
| msx1 | 0.018496711 | -2.690859208 |
| klhl1 | 0.03742636 | -2.687748187 |
| slc4a2 | 0.007627827 | -2.680047032 |
| capsl | 0.011891822 | -2.664456273 |
| loxl4 | 0.022114505 | -2.661113721 |
| baiap2l1 | 0.03754456 | -2.60757485 |
| adamts19 | 0.048876706 | -2.606113897 |

| | | |
|---|---|---|
| stra6 | 0.002937783 | -2.542785891 |
| ect2 | 0.042429856 | -2.52940864 |
| kremen1 | 0.007239386 | -2.521030647 |
| aox3 | 0.028082045 | -2.510594505 |
| kcna5 | 0.026191855 | -2.469113203 |
| gck | 0.032172226 | -2.465091242 |
| kif23 | 0.016191893 | -2.454606944 |
| tex15 | 0.002195268 | -2.443760632 |
| bmp2 | 0.023455788 | -2.443389673 |
| rgd1305627 | 0.034229562 | -2.44196131 |
| gucy1a3 | 0.010757803 | -2.428562484 |
| foxj1 | 0.016872062 | -2.391651269 |
| veph1 | 0.011011052 | -2.386164965 |
| mdk | 0.019725507 | -2.378077267 |
| crabp1 | 0.01039775 | -2.369561312 |
| cdkn1c | 0.014131503 | -2.362662498 |
| rhod | 0.029183082 | -2.359511332 |
| tmx3 | 0.001628 | -2.3434 |
| sgcg | 0.019739474 | -2.338383841 |
| iqcg | 0.03990227 | -2.335289007 |
| tc2n | 0.026126269 | -2.332276045 |
| st6galnac2 | 0.025818249 | -2.305893968 |
| spag8 | 0.039808313 | -2.304203455 |
| mlf1 | 0.02894849 | -2.300590195 |
| efcab1 | 0.020516243 | -2.280259095 |
| mmp2 | 0.030987837 | -2.277886068 |
| a2m | 0.020441097 | -2.271572512 |
| ccdc40 | 0.020678805 | -2.269620443 |
| ugt1a6 | 0.013743118 | -2.243115878 |
| fmo2 | 0.031456657 | -2.2301209 |
| dchs2 | 0.039732 | -2.2281 |
| foxc2 | 0.049745799 | -2.218669368 |
| slc2a12 | 0.012236431 | -2.206726441 |
| rgd1562658 | 0.031410148 | -2.19330108 |
| cdr2 | 0.00881332 | -2.193004046 |
| mme | 0.038163749 | -2.185067402 |
| meis2 | 0.00447832 | -2.179744277 |
| phactr2 | 0.035663178 | -2.177579533 |
| nkx2-1 | 0.021665364 | -2.149418274 |

| | | |
|---|---|---|
| zic3 | 0.041969705 | -2.141653606 |
| cdca7 | 0.046859388 | -2.141115666 |
| dclk3 | 0.013073264 | -2.116004731 |
| kcnh4 | 0.00954168 | -2.114672756 |
| tshr | 0.024831457 | -2.105065352 |
| rrm2 | 0.022181901 | -2.10287208 |
| pld5 | 0.046618906 | -2.088728221 |
| ccdc114 | 0.046726257 | -2.08872039 |
| wdr63 | 0.036370053 | -2.08511817 |
| epn3 | 0.049063663 | -2.084583077 |
| nags | 0.034334508 | -2.080688507 |
| rab11fip1 | 0.03879936 | -2.069411612 |
| pde10a | 0.007790575 | -2.056034697 |
| sfrp1 | 0.038557033 | -2.027334499 |
| mrgprf | 0.050094193 | -2.015498857 |
| trpm3 | 0.011836281 | -2.006748072 |
| pnpla1 | 0.024313512 | -2.003035925 |
| fscn2 | 0.043672691 | -1.995826518 |
| cmtm8 | 0.026220072 | -1.953229092 |
| art3 | 0.03398741 | -1.952871953 |
| sv2c | 0.030180042 | -1.938421536 |
| enkur | 0.007247059 | -1.929841025 |
| slc12a7 | 0.02569484 | -1.92895302 |
| gng7 | 0.029409212 | -1.925458637 |
| cdhr1 | 0.011945485 | -1.920667751 |
| rnf152 | 0.019384175 | -1.919886927 |
| ropn1l | 0.048912642 | -1.908465321 |
| tmem196 | 0.012893482 | -1.904947105 |
| elovl7 | 0.023567157 | -1.898330655 |
| cgnl1 | 0.032790334 | -1.894602458 |
| car14 | 0.038610531 | -1.893348086 |
| pcp4 | 0.004793751 | -1.890753347 |
| gypc | 0.037289743 | -1.889193661 |
| arpp21 | 0.01548476 | -1.881942635 |
| wdr16 | 0.035086877 | -1.879149221 |
| pde7b | 0.005317533 | -1.875256511 |
| gulp1 | 0.036254816 | -1.873737616 |
| htr6 | 0.027202831 | -1.873251341 |
| efhc1 | 0.048707494 | -1.872537072 |

| | | |
|---|---|---|
| crb1 | 0.046681155 | -1.862309394 |
| adcy5 | 0.02556711 | -1.855426181 |
| glb1l | 0.03648001 | -1.843557194 |
| entpd3 | 0.040472538 | -1.837958467 |
| pde1b | 0.012921251 | -1.826289356 |
| dnai2 | 0.045941131 | -1.824190064 |
| ajuba | 0.04509261 | -1.818637338 |
| mgp | 0.015872217 | -1.794458222 |
| synpo2 | 0.028229872 | -1.793177019 |
| slc5a5 | 0.033451315 | -1.785534665 |
| ngef | 0.011030604 | -1.779712948 |
| kcnk2 | 0.03588922 | -1.776253857 |
| il18bp | 0.045030819 | -1.764940477 |
| prkch | 0.019636104 | -1.759839036 |
| fam222a | 0.032405 | -1.745162497 |
| mospd1 | 0.030632031 | -1.739262913 |
| slc4a11 | 0.031017356 | -1.730187462 |
| dnali1 | 0.040621926 | -1.727554729 |
| npr3 | 0.038897286 | -1.726952586 |
| zic2 | 0.021866362 | -1.72140576 |
| dab2 | 0.050056623 | -1.721319796 |
| dlx5 | 0.034989467 | -1.712325875 |
| slc16a2 | 0.024724829 | -1.705648355 |
| prima1 | 0.027832916 | -1.69822529 |
| gpr153 | 0.037179369 | -1.696696094 |
| slc22a8 | 0.004446667 | -1.680997592 |
| slc31a1 | 0.038764033 | -1.675846742 |
| reck | 0.016234856 | -1.667822943 |
| rbp4 | 0.014518707 | -1.656040569 |
| unc13c | 0.023741902 | -1.643882481 |
| pbx3 | 0.048755338 | -1.63922706 |
| gnal | 0.011718377 | -1.634370872 |
| pdzd2 | 0.001674712 | -1.630261055 |
| tinagl1 | 0.041659169 | -1.616784363 |
| otx1 | 0.049037111 | -1.599034995 |
| col18a1 | 0.013303141 | -1.582550978 |
| cpne5 | 0.025647401 | -1.581127032 |
| fhod3 | 0.013723718 | -1.57501259 |
| mapk4 | 0.001914438 | -1.574898324 |

| | | |
|---|---|---|
| ptpn5 | 0.0299791 | -1.572017795 |
| pdp1 | 0.007555467 | -1.564884399 |
| grm4 | 0.035426652 | -1.563244736 |
| aspm | 0.0193032 | -1.556321479 |
| prkar2b | 0.014904587 | -1.554285603 |
| cobl | 0.042397256 | -1.548444797 |
| spock3 | 0.014515066 | -1.548321305 |
| lysmd3 | 0.042694574 | -1.546681904 |
| tnxa-ps1 | 0.039089974 | -1.534048573 |
| lrrk2 | 0.012354757 | -1.53399242 |
| arl4d | 0.017113892 | -1.52229816 |
| parp8 | 0.00026 | -1.434 |
| ablim2 | 0.010872022 | -1.263667768 |
| fbxl16 | 0.020655623 | -1.192157425 |
| nnt | 0.028565616 | -1.121049335 |
| spcs1 | 0.034490047 | -1.085119232 |
| actn2 | 0.057443761 | -1.078454923 |
| aadat | 0.04701434 | -1.03085743 |
| dnm1l | 0.054066083 | -1.030301559 |
| stx1b | 0.053329791 | -1.027537944 |
| ica1l | 0.048293132 | -1.023863323 |
| dlg4 | 0.047203117 | -1.019938326 |
| slit3 | 0.056557627 | -1.012924982 |
| pcdha6 | 0.05853034 | 1.00312314 |
| dld | 0.052759765 | 1.007649602 |
| mmp17 | 0.059865893 | 1.011637457 |
| slc9a7 | 0.058257385 | 1.012018548 |
| atcay | 0.04947356 | 1.013591708 |
| mrpl1 | 0.056871982 | 1.013738293 |
| lrpprc | 0.011192154 | 1.019480409 |
| ndufa11 | 0.056928326 | 1.021855021 |
| syn1 | 0.041661675 | 1.03309639 |
| slc1a2 | 0.052056105 | 1.035048014 |
| ndufs3 | 0.051414925 | 1.040718859 |
| mrpl30 | 0.054743682 | 1.049435305 |
| ddx1 | 0.055707738 | 1.051761856 |
| stxbp1 | 0.059443066 | 1.054004792 |
| efna5 | 0.045758946 | 1.055273602 |
| uqcrc2 | 0.051191201 | 1.060858489 |

| | | |
|---|---|---|
| slc25a4 | 0.052146504 | 1.071577041 |
| map1b | 0.054316713 | 1.080838065 |
| atp6v1e1 | 0.048803097 | 1.082965713 |
| slit1 | 0.053080385 | 1.091855839 |
| mrpl15 | 0.035739605 | 1.093077514 |
| mdh2 | 0.034762831 | 1.096284409 |
| cox5b | 0.019295679 | 1.114622045 |
| atp1a3 | 0.046113876 | 1.145751653 |
| hdac9 | 0.036111509 | 1.148091388 |
| pnpla3 | 0.049011845 | 1.170726685 |
| tomm70 | 0.002134 | 1.27238 |
| dock3 | 0.0170049 | 1.3245 |
| myo1f | 0.034741908 | 1.521536498 |
| unc13d | 0.046888368 | 1.524973399 |
| sh2b2 | 0.045458214 | 1.525581847 |
| plek | 0.019528101 | 1.532959911 |
| timd2 | 0.04686981 | 1.544986034 |
| ret | 0.001255903 | 1.563921541 |
| mvd | 0.000452782 | 1.577136871 |
| hcls1 | 0.011939878 | 1.609440256 |
| akap12 | 0.000118676 | 1.617107062 |
| ube2l6 | 0.040078029 | 1.654595073 |
| stk32b | 0.040235163 | 1.707798252 |
| rn28s | 0.002220397 | 1.725377211 |
| c3ar1 | 0.020342746 | 1.732802402 |
| fcgr2b | 0.000344735 | 1.752929881 |
| capg | 0.002911062 | 1.768239586 |
| gpnmb | 0.000017 | 1.801908992 |
| rt1-db1 | 0.043941583 | 1.80527256 |
| rab38 | 0.036819961 | 1.812687286 |
| ino80d | 0.02662815 | 1.816163169 |
| clec5a | 0.044948014 | 1.817380215 |
| rt1-ba | 0.023039512 | 1.822263016 |
| fyb | 0.026590312 | 1.83479121 |
| vav1 | 0.008546425 | 1.844506813 |
| arhgap30 | 0.019519239 | 1.859926456 |
| rt1-da | 0.016329497 | 1.864771106 |
| tpm2 | 0.040544359 | 1.867751306 |
| ttc26 | 0.018870309 | 1.938615815 |

| | | |
|---|---|---|
| arl11 | 0.002894024 | 1.942112943 |
| cd74 | 0.000739197 | 1.978959196 |
| gfap | 0.00000168 | 1.978993189 |
| itpr3 | 0.033641004 | 2.017529948 |
| chad | 0.026720524 | 2.041989354 |
| actn3 | 0.038334651 | 2.074877588 |
| rgd1309028 | 0.030369435 | 2.089050815 |
| app | 0.000000237 | 2.094685567 |
| ccdc166 | 0.037226099 | 2.109564197 |
| rt1-ce15 | 0.044818562 | 2.132332995 |
| cd22 | 0.000601887 | 2.138816519 |
| egfr | 0.040637994 | 2.155962221 |
| psen1 | 1.83E-14 | 2.195328155 |
| rt1-ce3 | 0.038698473 | 2.203868413 |
| cd68 | 0.002502175 | 2.212277945 |
| f13a1 | 0.033144367 | 2.230306075 |
| col25a1 | 0.003792 | 2.3231 |
| lsp1 | 0.002168612 | 2.371755028 |
| il2rg | 0.046237595 | 2.399498523 |
| def6 | 0.047413102 | 2.419212013 |
| col5a2 | 0.003553608 | 2.453284148 |
| spp1 | 0.047366988 | 2.462661416 |
| bcl3 | 0.030671927 | 2.493177288 |
| lag3 | 0.011843739 | 2.51077322 |
| itga5 | 0.024061827 | 2.5744065 |
| tmem170a | 0.021443017 | 2.636042943 |
| kcnn4 | 0.036065033 | 2.679776256 |
| slc16a3 | 0.039365467 | 2.707517709 |
| bst1 | 0.046019195 | 2.723233146 |
| rgd1559714 | 0.045290237 | 2.881282371 |
| cd72 | 0.038003774 | 2.933477607 |
| ncf2 | 0.007201558 | 2.957852011 |
| rgd1561958 | 0.025324673 | 2.975119097 |
| ltbp2 | 0.005463806 | 3.018671221 |
| hist2h3c2 | 0.039924358 | 3.051171829 |
| atp8b1 | 0.015209414 | 3.07145959 |
| pabpc2 | 0.022198769 | 3.078984328 |
| ccl3 | 0.005962152 | 3.098954149 |
| tnnt2 | 0.006053102 | 3.142348212 |

| | | |
|---|---|---|
| sim1 | 0.028599452 | 3.158747507 |
| fam109b | 0.026221652 | 3.188404463 |
| cox6b1 | 0.02203 | 3.2281 |
| has1 | 0.013642738 | 3.244373872 |
| loxl2 | 0.02613649 | 3.265485988 |
| ap3b2 | 0.00457 | 3.2829 |
| irx6 | 0.014346128 | 3.287290175 |
| serpine1 | 0.031848967 | 3.320712833 |
| hist1h2ak | 0.017576479 | 3.340877575 |
| hist2h4 | 0.03333672 | 3.37854974 |
| bcl2l11 | 0.014689059 | 3.40785099 |
| pde6b | 0.036666433 | 3.461997629 |
| fam187a | 0.012004572 | 3.585043815 |
| ccl4 | 0.033776167 | 3.657733748 |
| naglt1 | 0.001233369 | 3.698630377 |
| loc287004 | 0.00255303 | 3.728998604 |
| loc100363193 | 0.010436061 | 3.859918294 |
| prdm1 | 0.015743499 | 3.892981941 |
| aldoart2 | 0.017710683 | 3.983748826 |
| runx1 | 0.003513824 | 3.984637329 |
| col7a1 | 0.003608733 | 4.220155346 |
| snx32 | 0.0102 | 4.2289 |
| loc100910620 | 0.031452809 | 4.245853119 |
| lif | 0.033243717 | 4.315304012 |
| bcmo1 | 0.014134742 | 4.352704703 |
| cd200r1 | 0.004234 | 4.387565928 |
| avpr2 | 0.011960447 | 4.451751709 |
| naa11 | 0.010141473 | 4.457278371 |
| col5a1 | 0.000298791 | 4.541327014 |
| tnn | 0.011650813 | 4.621110389 |
| krt42 | 0.014645004 | 4.815306542 |
| tssk3 | 0.004743749 | 4.946519467 |
| olr200 | 0.031901743 | 5.050790081 |
| akp3 | 0.025870923 | 5.154030913 |
| hp | 0.004756671 | 5.234255755 |
| hist1h2aa | 0.017536722 | 5.268717018 |
| ddc8 | 0.020163285 | 5.37083411 |
| rmrp | 0.002045513 | 5.493773665 |
| gapdh-ps1 | 0.001689871 | 5.83911966 |

| | | |
|---|---|---|
| rgd1561102 | 0.002551461 | 5.853474947 |

**Supplementary table 3 : List of genes in TGNT pathways (males)**

| Pathway | Constituent Genes (Representative) | Risk in AD Context | Key References |
|---|---|---|---|
| **Ciliogenesis / ependymal & motile cilia programs (↑)** | FOXJ1, TPPP3, TEKT2/3/4, DNAH1, DNAI2, DNALI1, CCDC40, SPEF2 | Impaired ependymal cilia and CSF flow may reduce amyloid clearance and promote ventricular/inflammatory dysfunction in neurodegeneration. | Iqbal et al., *Brain Commun.*, 2022; Nedergaard & Goldman, *Science*, 2020 |
| **Transcriptional stress & cell-fate reprogramming (↑)** | ATF3, BTG2, TP73, NKX2-1, MSX1, EBF3 | Chronic stress-response activation and altered cell-fate programs reflect neuronal injury and maladaptive glial remodeling in AD. | Gjoneska et al., *Nat Neurosci.*, 2015; Mathys et al., *Nature*, 2019 |
| **Innate immune activation & phagocytosis (↑)** | CD68, CLEC7A, CYBB, CCL2, CCL3, LAG3, P2RX6 | Microglial activation is genetically and transcriptionally enriched in AD; excessive phagocytosis contributes to synaptic loss. | Keren-Shaul et al., *Cell*, 2017; Sims et al., *Nat Genet.*, 2017 |
| **Vascular remodeling & ECM deposition (↑)** | THBS1, THBS2, TGFB1, ADAMTS19, ADAMTSL4, COL8A1, SELP | BBB dysfunction and ECM remodeling promote neuroinflammation and cognitive decline in AD. | Sweeney et al., *Nat Neurosci.*, 2018; Montagne et al., *Neuron*, 2015 |
| **Neuromodulatory / GPCR signaling (↑)** | AVPR1A, CRHR2, HCRT, GALR3, SSTR5, AGTR2 | Altered neuromodulatory GPCR tone impacts cognition, stress signaling, and neuroimmune coupling in AD. | Thathiah & De Strooper, *Nat Rev Neurosci.*, 2011 |
| **Lipid / cholesterol & metabolic stress (↑)** | CYP27A1, ACAT2, LCAT, SLC25A4, BAAT | Lipid metabolism is a major genetic risk axis in AD; dysregulated cholesterol homeostasis influences amyloid processing. | Karch & Goate, *Nat Rev Genet.*, 2015; Sims et al., *Nat Genet.*, 2017 |
| **Antigen presentation & inflammatory signaling (↑)** | CIITA, CD74, RT1-DA, RT1-DB1, MR1, CXCL16 | MHC-II pathways are strongly upregulated in AD and correlate with microglial activation states. | Mathys et al., *Nature*, 2019; Keren-Shaul et al., *Cell*, 2017 |

| | | | |
|---|---|---|---|
| **Cell-cycle re-entry / proliferation programs (↑)** | AURKB, UBE2C, PRC1, CDCA2, CDKN1C | Aberrant neuronal cell-cycle re-entry is associated with neurodegeneration and precedes apoptosis in AD. | Moh et al., *Neuron*, 2011; Herrup, *Nat Rev Neurosci.*, 2010 |
| **Synaptic transmission & release machinery (↓)** | SNAP25, SYP, SYT1, STX1A/B, UNC13A, RIMS2, GRIA2/3, GRIN2A/B | Synapse loss is the strongest correlate of cognitive decline; complement-mediated pruning contributes to early AD pathology. | DeKosky & Scheff, *Ann Neurol.*, 1990; Hong et al., *Science*, 2016 |
| **Transcriptional maintenance & neuronal homeostasis (↓)** | MEF2C, ELAVL4, CDYL, ZPR1 | Reduced neuronal maintenance programs reflect vulnerable neuronal states in AD. | Gjoneska et al., *Nat Neurosci.*, 2015; Mathys et al., *Nature*, 2019 |
| **Vascular stability & endothelial maintenance (↓)** | KDR, ROBO4, ESAM, APLN | Loss of vascular integrity exacerbates hypoperfusion and neurodegeneration. | Sweeney et al., *Nat Neurosci.*, 2018 |
| **Immune regulatory / suppressive signaling (↓)** | SIGIRR, IRF4 | Reduced immune regulation promotes chronic neuroinflammation in AD brains. | Sims et al., *Nat Genet.*, 2017 |
| **Ion channels & neuronal excitability (↓)** | SCN2A, SCN8A, KCNQ5, KCNC2, TRPC3 | Network hyperexcitability and ion channel dysregulation contribute to cognitive decline and seizure susceptibility in AD. | Busche & Konnerth, *Nat Rev Neurosci.*, 2016 |
| **Developmental patterning & fate specification (↓)** | NEUROD6, EPHA5, KALRN, CNTNAP2 | Loss of synaptic organization and guidance programs destabilizes circuit connectivity in AD. | Selkoe, *Physiol Rev.*, 2002 |

#### Supplementary table 4 : List of genes in TGNT pathways (females)

| Pathway | Constituent Genes (Representative) | Risk in AD Context | Key References |
|---|---|---|---|
| **Microglia activation / DAM-like (↑)** | CD68, CLEC5A, C3AR1, GPNMB, FCGR2B, NCF2, LSP1 | Disease-associated microglia (DAM) states are strongly enriched in AD brains and genetically linked to risk loci; excessive activation promotes synaptic pruning and neuroinflammation. | Keren-Shaul et al., *Cell*, 2017; Sims et al., *Nat Genet.*, 2017 |

| | | | |
|---|---|---|---|
| **ECM remodeling / fibrosis (↑)** | COL5A1, COL5A2, LOXL2, SERPINE1, LTBP2, HAS1 | ECM remodeling contributes to vascular stiffening, BBB compromise, and altered synaptic microenvironment in AD. | Sweeney et al., *Nat Neurosci.*, 2018; Montagne et al., *Neuron*, 2015 |
| **Antigen presentation (MHC-II) (↑)** | CD74, RT1-CE3, RT1-CE15, RT1-DB1, CIITA | MHC-II–mediated antigen presentation is elevated in AD microglia and correlates with inflammatory progression. | Mathys et al., *Nature*, 2019 |
| **Kinase / growth factor signaling (↑)** | EGFR, RET, ITGA5, VAV1, SH2B2 | Dysregulated kinase signaling impacts synaptic plasticity, inflammation, and APP processing in AD. | Thathiah & De Strooper, *Nat Rev Neurosci.*, 2011 |
| **Cytokine / chemokine signaling (↑)** | CCL3, CCL4, IL2RG, LIF | Chronic cytokine signaling drives neuroinflammation and synaptic dysfunction in AD. | Heneka et al., *Lancet Neurol.*, 2015 |
| **Amyloid / APP & glial stress (↑)** | APP, PSEN1, GFAP | Altered APP processing and astrocytic activation are central drivers of amyloid pathology and reactive gliosis. | Hardy & Selkoe, *Science*, 2002 |
| **Metabolic stress / transport (↑)** | SLC16A3, ATP8B1, MVD, BCL2L11 | Metabolic dysfunction and mitochondrial stress precede overt pathology and contribute to neuronal vulnerability. | Wang et al., *Nat Med.*, 2020 |
| **Apoptosis / cell death programs (↑)** | BCL2L11, BAD, CDKN1C | Apoptotic priming and aberrant cell-cycle activity contribute to neuronal loss in AD. | Moh et al., *Neuron*, 2011 |
| **Synaptic signaling / release machinery (↓)** | GRM4, SLC17A6, TAC3, GAL, P2RY1 | Loss of synaptic signaling genes correlates strongly with cognitive decline and circuit dysfunction. | DeKosky & Scheff, *Ann Neurol.*, 1990 |
| **GPCR / neurotransmission (↓)** | HTR2C, ADR A1B, ADR A1D, CNR1, SSTR3 | Impaired neuromodulatory GPCR tone affects cognition and neuronal resilience in AD. | Thathiah & De Strooper, *Nat Rev Neurosci.*, 2011 |
| **cAMP / cGMP & phosphodiesterases (↓)** | PDE1B, PDE7B, PDE10A, ADCY5 | Dysregulated cyclic nucleotide signaling impairs memory-related plasticity pathways in AD. | Kelly, *Trends Mol Med.*, 2018 |
| **Ciliogenesis / motility programs (↓)** | DNAI2, WDR16, ENKUR, ROPN1L | Ciliary dysfunction may influence CSF flow and neuroinflammatory balance. | Iqbal et al., *Brain Commun.*, 2022 |

| | | | |
|---|---|---|---|
| **Cell survival signaling (↓)** | AKT1, PRKCH, NGEF | Reduced pro-survival signaling increases neuronal susceptibility to degeneration. | Wang et al., *Nat Med.*, 2020 |
| **ECM / vascular structure (↓)** | MMP2, ADAMTS19, COL18A1 | Loss of vascular structural integrity contributes to BBB instability and neurodegeneration. | Sweeney et al., *Nat Neurosci.*, 2018 |
| **Cell cycle / DNA maintenance (↓)** | CDK1, TOP2A, MCM10, EXO1 | Aberrant reactivation of cell-cycle machinery in neurons is associated with apoptosis and neurodegeneration. | Herrup, *Nat Rev Neurosci.*, 2010 |

#### Supplementary table 5 : Conservation of human AD transcriptome with TgF344-AD rat transcriptome (Males)

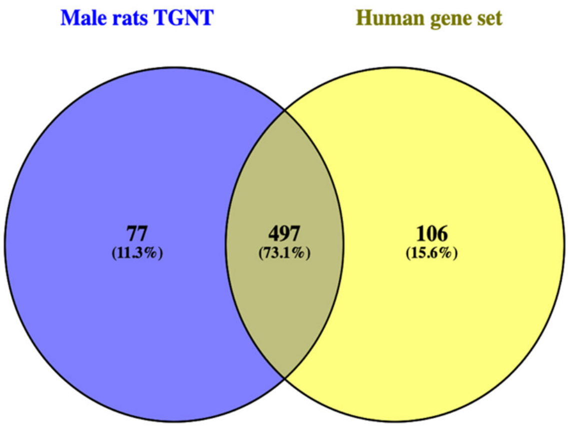

| Males rat TGNT | AGORA dataset for Males | Common genes |
|---|---|---|
| psen1 | ifit3 | psen1 |
| app | syne4 | app |
| gfap | efcab10 | gfap |
| gpnmb | lrrc46 | gpnmb |
| gpr88 | odf3b | gpr88 |
| rgs9 | cp | rgs9 |
| six3 | drc1 | six3 |
| adora2a | myo7a | adora2a |
| lhx8 | tgfbi | lhx8 |
| gpr6 | nxnl1 | gpr6 |
| ecel1 | pqlc3 | ecel1 |
| sh3rf2 | aldh1l2 | sh3rf2 |
| akap12 | enkur | akap12 |

| | | |
|---|---|---|
| syndig1l | olfm4 | syndig1l |
| slc22a3 | rgs1 | slc22a3 |
| drd1 | rsph1 | drd1 |
| foxp2 | cd68 | foxp2 |
| slc4a5 | sik1 | slc4a5 |
| h19 | erich2 | h19 |
| igf2 | mfap2 | igf2 |
| folr1 | efcab1 | folr1 |
| tmem27 | cdc153 | tmem27 |
| col5a1 | tekt2 | col5a1 |
| tmem72 | fbxl13 | tmem72 |
| penk | ciita | penk |
| fcgr2b | ccdc146 | fcgr2b |
| tnni3 | prc1 | tnni3 |
| ace | impg2 | ace |
| mvd | tyrp1 | mvd |
| scn4b | ifltd1 | scn4b |
| kb15 | hist1h2bd | kb15 |
| cd22 | dydc2 | cd22 |
| f5 | igfbpl1 | f5 |
| drd2 | gpr139 | drd2 |
| sbk2 | fam154b | sbk2 |
| kl | kcnk16 | kl |
| sulf1 | baat | sulf1 |
| itgb6 | mrap | itgb6 |
| cdh3 | sytl3 | cdh3 |
| lect1 | scube3 | lect1 |
| cd74 | capsl | cd74 |
| rgd1559696 | bcl2l15 | rgd1559696 |
| mfrp | ccdc114 | mfrp |
| slc5a7 | cdh3 | slc5a7 |
| enpp2 | calcr | enpp2 |
| itpripl1 | slc7a9 | itpripl1 |
| ppp1r1b | ampd1 | ppp1r1b |
| trpv4 | ttc25 | trpv4 |
| hist1h2bo | daw1 | hist1h2bo |
| scgb1c1 | prrg4 | scgb1c1 |
| samd3 | slc44a4 | samd3 |
| col3a1 | guca1b | col3a1 |

| | | |
|---|---|---|
| igfbpl1 | lrrc34 | igfbpl1 |
| inmt | trpv4 | inmt |
| krt71 | ttc29 | krt71 |
| tac1 | slc22a3 | tac1 |
| kcne2 | mpo | kcne2 |
| abca4 | rib1 | abca4 |
| zic1 | bbox1 | zic1 |
| calcr | ccdc40 | calcr |
| adipoq | pkhd1l1 | adipoq |
| col1a2 | mpzl2 | col1a2 |
| lhx1 | azgp1 | lhx1 |
| naglt1 | pglyrp1 | naglt1 |
| ret | ace | ret |
| pappa | galr3 | pappa |
| kcnj13 | ak7 | kcnj13 |
| h1foo | ly9 | h1foo |
| jph2 | enpp2 | jph2 |
| slco1a5 | tmem212 | slco1a5 |
| pdzd2 | krt8 | pdzd2 |
| gapdh-ps1 | slc14a2 | gapdh-ps1 |
| sult1c3 | cd5 | sult1c3 |
| htr1d | lbp | htr1d |
| myl6 | dnah1 | myl6 |
| leap2 | aurkb | leap2 |
| ogn | pla2g5 | ogn |
| nexn | itgb6 | nexn |
| mapk4 | spag6 | mapk4 |
| tbc1d10c | aipl1 | tbc1d10c |
| rmrp | dynlrb2 | rmrp |
| ttr | mlf1 | ttr |
| slc13a4 | abcb11 | slc13a4 |
| lsp1 | kcne2 | lsp1 |
| tex15 | iqcg | tex15 |
| cldn2 | kcnq1 | cldn2 |
| rn28s | slc16a8 | rn28s |
| rasd2 | ahsp | rasd2 |
| slc18a3 | wdr63 | slc18a3 |
| vwce | pdc | vwce |
| otx2 | slpi | otx2 |

| | | |
|---|---|---|
| trh | dcdc5 | trh |
| pla2g5 | akap14 | pla2g5 |
| asic4 | corin | asic4 |
| cd68 | clec7a | cd68 |
| slc35d3 | ubd | slc35d3 |
| neu2 | krt18 | neu2 |
| rgd1561102 | tex28 | rgd1561102 |
| loc287004 | lrit1 | loc287004 |
| ccnb1ip1 | inmt | ccnb1ip1 |
| prlr | ttll10 | prlr |
| trat1 | hoxb5 | trat1 |
| irx3 | rbp3 | irx3 |
| arl11 | best2 | arl11 |
| capg | c1qtnf3 | capg |
| stra6 | tnmd | stra6 |
| lepr | agr3 | lepr |
| hnf4a | col8a1 | hnf4a |
| atp6ap1l | irx4 | atp6ap1l |
| fgf1 | sstr5 | fgf1 |
| cmtm1 | mc3r | cmtm1 |
| clic6 | adipoq | clic6 |
| runx1 | gata6 | runx1 |
| pcp4l1 | lrrc18 | pcp4l1 |
| col5a2 | mak | col5a2 |
| col7a1 | mylk2 | col7a1 |
| ccl9 | selp | ccl9 |
| cartpt | prph2 | cartpt |
| isl1 | cr2 | isl1 |
| rgd1561795 | tmem232 | rgd1561795 |
| gbx2 | lgals5 | gbx2 |
| cd200r1 | fabp12 | cd200r1 |
| sostdc1 | rbp7 | sostdc1 |
| dazl | agtr2 | dazl |
| slc22a8 | abca4 | slc22a8 |
| meis2 | pih1d3 | meis2 |
| itgb1bp2 | slc22a2 | itgb1bp2 |
| mpz | pon3 | mpz |
| tssk3 | aqp1 | tssk3 |
| hp | steap1 | hp |

| | | |
|---|---|---|
| pcp4 | uncx | pcp4 |
| loc691083 | gdf7 | loc691083 |
| tcf21 | padi4 | tcf21 |
| kcnq1 | pde6h | kcnq1 |
| glycam1 | lpo | glycam1 |
| pon3 | crygs | pon3 |
| rbm47 | pon1 | rbm47 |
| gpr52 | hormad2 | gpr52 |
| fam160a1 | zscan10 | fam160a1 |
| pde7b | tmem27 | pde7b |
| tbx6 | pax3 | tbx6 |
| ltbp2 | efhb | ltbp2 |
| tmem184a | ccdc60 | tmem184a |
| frem1 | trim6 | frem1 |
| il13ra2 | lmx1a | il13ra2 |
| mpzl2 | otx2 | mpzl2 |
| ptprq | il13ra2 | ptprq |
| itk | impg1 | itk |
| clul1 | cldn2 | clul1 |
| melk | bhmt2 | melk |
| ccl3 | cd8b | ccl3 |
| tnnt2 | kcnj13 | tnnt2 |
| rgd1561161 | tmem184a | rgd1561161 |
| lgals5 | folr1 | lgals5 |
| spetex-2a | mfrp | spetex-2a |
| fetub | sox14 | fetub |
| rarb | lilrb4 | rarb |
| aqp1 | clic6 | aqp1 |
| klk4 | kl | klk4 |
| trpv5 | slc4a5 | trpv5 |
| pon1 | ttr | pon1 |
| ncf2 | tmem72 | ncf2 |
| kremen1 | tulp1 | kremen1 |
| enkur | sostdc1 | enkur |
| hes3 | nrg4 | hes3 |
| pdp1 | gngt1 | pdp1 |
| igfbp2 | krt19 | igfbp2 |
| slc4a2 | f5 | slc4a2 |
| mdfic | cxcl11 | mdfic |

| | | |
|---|---|---|
| lhx5 | tcf21 | lhx5 |
| steap1 | ttll2 | steap1 |
| pde10a | six6 | pde10a |
| scube3 | hnrnpu | scube3 |
| il20ra | psen1 | il20ra |
| iqgap3 | app | iqgap3 |
| gbx1 | gfap | gbx1 |
| camp | gpnmb | camp |
| slc16a12 | gpr88 | slc16a12 |
| vav1 | rgs9 | vav1 |
| s100g | six3 | s100g |
| mxd3 | adora2a | mxd3 |
| dmkn | lhx8 | dmkn |
| cdr2 | gpr6 | cdr2 |
| kcnv2 | ecel1 | kcnv2 |
| lmx1a | sh3rf2 | lmx1a |
| kcnh4 | akap12 | kcnh4 |
| brs3 | syndig1l | brs3 |
| gucy2f | slc22a3 | gucy2f |
| naa11 | drd1 | naa11 |
| pih1d3 | foxp2 | pih1d3 |
| mcoln3 | slc4a5 | mcoln3 |
| crabp1 | h19 | crabp1 |
| loc100363193 | igf2 | loc100363193 |
| mc3r | folr1 | mc3r |
| cdx4 | tmem27 | cdx4 |
| gucy1a3 | col5a1 | gucy1a3 |
| loc654482 | tmem72 | loc654482 |
| cdk1 | penk | cdk1 |
| zscan10 | fcgr2b | zscan10 |
| veph1 | tnni3 | veph1 |
| ngef | ace | ngef |
| adra2b | mvd | adra2b |
| fbxo39 | scn4b | fbxo39 |
| mcoln2 | kb15 | mcoln2 |
| aldh1a3 | cd22 | aldh1a3 |
| loc102548399 | f5 | loc102548399 |
| slc16a8 | drd2 | slc16a8 |
| tnn | sbk2 | tnn |

| | | |
|---|---|---|
| gnal | kl | gnal |
| trpm3 | sulf1 | trpm3 |
| lag3 | itgb6 | lag3 |
| capsl | cdh3 | capsl |
| hcls1 | lect1 | hcls1 |
| cdhr1 | cd74 | cdhr1 |
| avpr2 | rgd1559696 | avpr2 |
| fam187a | mfrp | fam187a |
| rgd1563692 | slc5a7 | rgd1563692 |
| slc2a12 | enpp2 | slc2a12 |
| pcp2 | itpripl1 | pcp2 |
| lrrk2 | ppp1r1b | lrrk2 |
| bhmt2 | trpv4 | bhmt2 |
| tns4 | hist1h2bo | tns4 |
| igfals | scgb1c1 | igfals |
| pitx2 | samd3 | pitx2 |
| ido1 | col3a1 | ido1 |
| insrr | igfbpl1 | insrr |
| coch | inmt | coch |
| tmem196 | krt71 | tmem196 |
| nr5a1 | tac1 | nr5a1 |
| pde1b | kcne2 | pde1b |
| dclk3 | abca4 | dclk3 |
| rgd1308065 | zic1 | rgd1308065 |
| col18a1 | calcr | col18a1 |
| adamtsl4 | adipoq | adamtsl4 |
| has1 | col1a2 | has1 |
| fhod3 | lhx1 | fhod3 |
| ugt1a6 | naglt1 | ugt1a6 |
| cdkn1c | ret | cdkn1c |
| bcmo1 | pappa | bcmo1 |
| runx3 | kcnj13 | runx3 |
| lilrb4 | h1foo | lilrb4 |
| irx6 | jph2 | irx6 |
| spock3 | slco1a5 | spock3 |
| rbp4 | pdzd2 | rbp4 |
| krt42 | gapdh-ps1 | krt42 |
| retn | sult1c3 | retn |
| bcl2l11 | htr1d | bcl2l11 |

| | | |
|---|---|---|
| c1qtnf3 | myl6 | c1qtnf3 |
| prkar2b | leap2 | prkar2b |
| atp8b1 | ogn | atp8b1 |
| acot12 | nexn | acot12 |
| rasgrp2 | mapk4 | rasgrp2 |
| arpp21 | tbc1d10c | arpp21 |
| fam46a | rmrp | fam46a |
| atp4a | ttr | atp4a |
| prdm1 | slc13a4 | prdm1 |
| mgp | lsp1 | mgp |
| kif23 | tex15 | kif23 |
| reck | cldn2 | reck |
| rt1-da | rn28s | rt1-da |
| hs3st3a1 | rasd2 | hs3st3a1 |
| lrrc18 | slc18a3 | lrrc18 |
| mybl2 | vwce | mybl2 |
| ces1d | otx2 | ces1d |
| foxj1 | trh | foxj1 |
| xcl1 | pla2g5 | xcl1 |
| slpil2 | asic4 | slpil2 |
| arl4d | cd68 | arl4d |
| krt85 | slc35d3 | krt85 |
| pou4f3 | neu2 | pou4f3 |
| hist1h2aa | rgd1561102 | hist1h2aa |
| crb3 | loc287004 | crb3 |
| hist1h2ak | ccnb1ip1 | hist1h2ak |
| aldoart2 | prlr | aldoart2 |
| vom1r16 | trat1 | vom1r16 |
| msx1 | irx3 | msx1 |
| ttc26 | arl11 | ttc26 |
| serpinc1 | capg | serpinc1 |
| hist1h1b | stra6 | hist1h1b |
| rnf152 | lepr | rnf152 |
| arhgap30 | hnf4a | arhgap30 |
| plek | atp6ap1l | plek |
| prkch | fgf1 | prkch |
| mdk | cmtm1 | mdk |
| sgcg | clic6 | sgcg |
| rgd1560608 | runx1 | rgd1560608 |

| | | |
|---|---|---|
| slc10a4 | pcp4l1 | slc10a4 |
| ddc8 | col5a2 | ddc8 |
| c3ar1 | col7a1 | c3ar1 |
| a2m | ccl9 | a2m |
| efcab1 | cartpt | efcab1 |
| ccdc40 | isl1 | ccdc40 |
| mak | rgd1561795 | mak |
| nmu | gbx2 | nmu |
| tmem170a | cd200r1 | tmem170a |
| loc100361092 | sostdc1 | loc100361092 |
| klk7 | dazl | klk7 |
| nkx2-1 | slc22a8 | nkx2-1 |
| zic2 | meis2 | zic2 |
| cxcl13 | itgb1bp2 | cxcl13 |
| loxl4 | mpz | loxl4 |
| rrm2 | tssk3 | rrm2 |
| pabpc2 | hp | pabpc2 |
| rgd1303271 | pcp4 | rgd1303271 |
| slc14a2 | loc691083 | slc14a2 |
| slc5a12 | tcf21 | slc5a12 |
| reg3a | kcnq1 | reg3a |
| top2a | glycam1 | top2a |
| rt1-ba | pon3 | rt1-ba |
| bmp2 | rbm47 | bmp2 |
| elovl7 | gpr52 | elovl7 |
| unc13c | fam160a1 | unc13c |
| uncx | pde7b | uncx |
| itga5 | tbx6 | itga5 |
| foxb1 | ltbp2 | foxb1 |
| klra5 | tmem184a | klra5 |
| ces2 | frem1 | ces2 |
| fam71f1 | il13ra2 | fam71f1 |
| pnpla1 | mpzl2 | pnpla1 |
| pdzk1ip1 | ptprq | pdzk1ip1 |
| nepn | itk | nepn |
| spink5 | clul1 | spink5 |
| rgd1561870 | melk | rgd1561870 |
| loc689713 | ccl3 | loc689713 |
| slc16a2 | tnnt2 | slc16a2 |

| | | |
|---|---|---|
| tshr | rgd1561161 | tshr |
| prkag3 | lgals5 | prkag3 |
| slc7a12 | spetex-2a | slc7a12 |
| sct | fetub | sct |
| rgd1561958 | rarb | rgd1561958 |
| fgf20 | aqp1 | fgf20 |
| klk1 | klk4 | klk1 |
| adcy5 | trpv5 | adcy5 |
| cpne5 | pon1 | cpne5 |
| slc12a7 | ncf2 | slc12a7 |
| st6galnac2 | kremen1 | st6galnac2 |
| akp3 | enkur | akp3 |
| tc2n | hes3 | tc2n |
| loxl2 | pdp1 | loxl2 |
| kcna5 | igfbp2 | kcna5 |
| cmtm8 | slc4a2 | cmtm8 |
| fam109b | mdfic | fam109b |
| cyp11b3 | lhx5 | cyp11b3 |
| fyb | steap1 | fyb |
| ino80d | pde10a | ino80d |
| chad | scube3 | chad |
| cenpf | il20ra | cenpf |
| car13 | iqgap3 | car13 |
| fbp1 | gbx1 | fbp1 |
| vgll2 | camp | vgll2 |
| ldhal6b | slc16a12 | ldhal6b |
| htr6 | vav1 | htr6 |
| kiss1 | s100g | kiss1 |
| ms4a14 | mxd3 | ms4a14 |
| krt18 | dmkn | krt18 |
| prima1 | cdr2 | prima1 |
| gsc2 | kcnv2 | gsc2 |
| aox3 | lmx1a | aox3 |
| chrna9 | kcnh4 | chrna9 |
| otc | brs3 | otc |
| synpo2 | gucy2f | synpo2 |
| slc23a3 | naa11 | slc23a3 |
| sim1 | pih1d3 | sim1 |
| ifna4 | mcoln3 | ifna4 |

| | | |
|---|---|---|
| spag11c | crabp1 | spag11c |
| mlf1 | loc100363193 | mlf1 |
| cpn1 | mc3r | cpn1 |
| lenep | cdx4 | lenep |
| rgd1306625 | gucy1a3 | rgd1306625 |
| rhod | loc654482 | rhod |
| gng7 | cdk1 | gng7 |
| lhfpl1 | zscan10 | lhfpl1 |
| ptpn5 | veph1 | ptpn5 |
| sv2c | ngef | sv2c |
| rgd1309028 | adra2b | rgd1309028 |
| mospd1 | fbxo39 | mospd1 |
| bcl3 | mcoln2 | bcl3 |
| mmp2 | aldh1a3 | mmp2 |
| slc4a11 | loc102548399 | slc4a11 |
| krt75 | slc16a8 | krt75 |
| rgd1562658 | tnn | rgd1562658 |
| loc100910620 | gnal | loc100910620 |
| fmo2 | trpm3 | fmo2 |
| mael | lag3 | mael |
| serpine1 | capsl | serpine1 |
| olr200 | hcls1 | olr200 |
| txk | cdhr1 | txk |
| gck | avpr2 | gck |
| fam222a | fam187a | fam222a |
| clec4e | rgd1563692 | clec4e |
| cgnl1 | slc2a12 | cgnl1 |
| f13a1 | pcp2 | f13a1 |
| six1 | lrrk2 | six1 |
| smim22 | bhmt2 | smim22 |
| lif | tns4 | lif |
| hist2h4 | igfals | hist2h4 |
| slc5a5 | pitx2 | slc5a5 |
| bhmt | ido1 | bhmt |
| itpr3 | insrr | itpr3 |
| efhb | coch | efhb |
| ccl4 | tmem196 | ccl4 |
| art3 | nr5a1 | art3 |
| loc171161 | pde1b | loc171161 |

| | | |
|---|---|---|
| rgd1305627 | dclk3 | rgd1305627 |
| nags | rgd1308065 | nags |
| myo1f | col18a1 | myo1f |
| dlx5 | adamtsl4 | dlx5 |
| wdr16 | has1 | wdr16 |
| grm4 | fhod3 | grm4 |
| corin | ugt1a6 | corin |
| phactr2 | cdkn1c | phactr2 |
| kcnk2 | bcmo1 | kcnk2 |
| kcnn4 | runx3 | kcnn4 |
| gulp1 | lilrb4 | gulp1 |
| wdr63 | irx6 | wdr63 |
| glb1l | spock3 | glb1l |
| pde6b | rbp4 | pde6b |
| rab38 | krt42 | rab38 |
| gpr153 | retn | gpr153 |
| ccdc166 | bcl2l11 | ccdc166 |
| gypc | c1qtnf3 | gypc |
| klhl1 | prkar2b | klhl1 |
| msc | atp8b1 | msc |
| ifltd1 | acot12 | ifltd1 |
| baiap2l1 | rasgrp2 | baiap2l1 |
| cd72 | arpp21 | cd72 |
| mme | fam46a | mme |
| actn3 | atp4a | actn3 |
| sfrp1 | prdm1 | sfrp1 |
| car14 | mgp | car14 |
| rt1-ce3 | kif23 | rt1-ce3 |
| slc31a1 | reck | slc31a1 |
| rab11fip1 | rt1-da | rab11fip1 |
| npr3 | hs3st3a1 | npr3 |
| clec2d | lrrc18 | clec2d |
| tnxa-ps1 | mybl2 | tnxa-ps1 |
| slc16a3 | ces1d | slc16a3 |
| best3 | foxj1 | best3 |
| spag8 | xcl1 | spag8 |
| iqcg | slpil2 | iqcg |
| hist2h3c2 | arl4d | hist2h3c2 |
| ube2l6 | krt85 | ube2l6 |

| | | |
|---|---|---|
| mcm10 | pou4f3 | mcm10 |
| stk32b | hist1h2aa | stk32b |
| entpd3 | crb3 | entpd3 |
| tpm2 | hist1h2ak | tpm2 |
| dnali1 | aldoart2 | dnali1 |
| egfr | vom1r16 | egfr |
| tinagl1 | msx1 | tinagl1 |
| zic3 | ttc26 | zic3 |
| cobl | serpinc1 | cobl |
| ect2 | hist1h1b | ect2 |
| exo1 | rnf152 | exo1 |
| stat4 | arhgap30 | stat4 |
| lysmd3 | plek | lysmd3 |
| tgm3 | prkch | tgm3 |
| fscn2 | mdk | fscn2 |
| ccdc60 | sgcg | ccdc60 |
| rt1-db1 | rgd1560608 | rt1-db1 |
| chrdl2 | slc10a4 | chrdl2 |
| hnf1b | ddc8 | hnf1b |
| rt1-ce15 | c3ar1 | rt1-ce15 |
| clec5a | a2m | clec5a |
| il18bp | efcab1 | il18bp |
| ajuba | ccdc40 | ajuba |
| rgd1559714 | mak | rgd1559714 |
| sh2b2 | nmu | sh2b2 |
| sytl3 | tmem170a | sytl3 |
| dnai2 | loc100361092 | dnai2 |
| bst1 | klk7 | bst1 |
| il2rg | nkx2-1 | il2rg |
| pou1f1 | zic2 | pou1f1 |
| gdf7 | cxcl13 | gdf7 |
| pld5 | loxl4 | pld5 |
| crb1 | rrm2 | crb1 |
| ccdc114 | pabpc2 | ccdc114 |
| cdca7 | rgd1303271 | cdca7 |
| timd2 | slc14a2 | timd2 |
| unc13d | slc5a12 | unc13d |
| loc689927 | reg3a | loc689927 |
| crygs | top2a | crygs |

| | | |
|---|---|---|
| rgd1565655 | rt1-ba | rgd1565655 |
| rrh | bmp2 | rrh |
| spp1 | elovl7 | spp1 |
| def6 | unc13c | def6 |
| gja6 | uncx | gja6 |
| vom1r88 | itga5 | vom1r88 |
| lyzl6 | foxb1 | lyzl6 |
| cmah | klra5 | cmah |
| txndc2 | ces2 | txndc2 |
| mirlet7d | fam71f1 | mirlet7d |
| cb707485 | pnpla1 | cb707485 |
| olr1667 | pdzk1ip1 | olr1667 |
| testin | nepn | testin |
| cyp2c24 | spink5 | cyp2c24 |
| fgg | rgd1561870 | fgg |
| lrrc52 | loc689713 | agtr2 |
| odf3l2 | slc16a2 | |
| dmrtc1c1 | tshr | |
| krt86 | prkag3 | |
| cd69 | slc7a12 | |
| agtr2 | sct | |
| cryge | rgd1561958 | |
| dcm5 | fgf20 | |
| oas1h | klk1 | |
| ugt1a9 | adcy5 | |
| tcp10b | cpne5 | |
| cldn22 | slc12a7 | |
| olr1653 | st6galnac2 | |
| sell | akp3 | |
| adh7 | tc2n | |
| mirlet7c-2 | loxl2 | |
| mir3572 | kcna5 | |
| olr1869 | cmtm8 | |
| efhc1 | fam109b | |
| pbx3 | cyp11b3 | |
| gfi1b | fyb | |
| gykl1 | ino80d | |
| adamts19 | chad | |
| ropn1l | cenpf | |

| | |
|---|---|
| lrr1 | car13 |
| otx1 | fbp1 |
| hepacam2 | vgll2 |
| krt34 | ldhal6b |
| olr387 | htr6 |
| epn3 | kiss1 |
| cyp11a1 | ms4a14 |
| slamf6 | krt18 |
| crygf | prima1 |
| krt1 | gsc2 |
| foxc2 | aox3 |
| dab2 | chrna9 |
| mrgprf | otc |
| troap | synpo2 |
| evpl | slc23a3 |
| prelp | sim1 |
| igsf9 | ifna4 |
| mir421 | spag11c |
| adcy7 | mlf1 |
| vwa5b1 | cpn1 |
| pof1b | lenep |
| rdh7 | rgd1306625 |
| fam83a | rhod |
| rgd1561778 | gng7 |
| tead2 | lhfpl1 |
| cox8b | ptpn5 |
| llgl2 | sv2c |
| dpp4 | rgd1309028 |
| scara5 | mospd1 |
| kcna10 | bcl3 |
| rgd1563104 | mmp2 |
| trim40 | slc4a11 |
| ugt1a5 | krt75 |
| apobec1 | rgd1562658 |
| krt16 | loc100910620 |
| rin3 | fmo2 |
| s100a5 | mael |
| kctd8 | serpine1 |
| zim1 | olr200 |

| | |
|---|---|
| sp8 | txk |
| upk1b | gck |
| nlrp4 | fam222a |
| iqub | clec4e |
| vom2r60 | cgnl1 |
| lipogenin | f13a1 |
| zmynd10 | six1 |
| pax8 | smim22 |
| gpx2 | lif |
| fgf19 | hist2h4 |
| epha1 | slc5a5 |
| il22ra1 | bhmt |
| kcnmb3 | itpr3 |
| magea4 | efhb |
| olr176 | ccl4 |
| | art3 |
| | loc171161 |
| | rgd1305627 |
| | nags |
| | myo1f |
| | dlx5 |
| | wdr16 |
| | grm4 |
| | corin |
| | phactr2 |
| | kcnk2 |
| | kcnn4 |
| | gulp1 |
| | wdr63 |
| | glb1l |
| | pde6b |
| | rab38 |
| | gpr153 |
| | ccdc166 |
| | gypc |
| | klhl1 |
| | msc |
| | ifltd1 |
| | baiap2l1 |

| | |
|---|---|
| | cd72 |
| | mme |
| | actn3 |
| | sfrp1 |
| | car14 |
| | rt1-ce3 |
| | slc31a1 |
| | rab11fip1 |
| | npr3 |
| | clec2d |
| | tnxa-ps1 |
| | slc16a3 |
| | best3 |
| | spag8 |
| | iqcg |
| | hist2h3c2 |
| | ube2l6 |
| | mcm10 |
| | stk32b |
| | entpd3 |
| | tpm2 |
| | dnali1 |
| | egfr |
| | tinagl1 |
| | zic3 |
| | cobl |
| | ect2 |
| | exo1 |
| | stat4 |
| | lysmd3 |
| | tgm3 |
| | fscn2 |
| | ccdc60 |
| | rt1-db1 |
| | chrdl2 |
| | hnf1b |
| | rt1-ce15 |
| | clec5a |
| | il18bp |

| | |
|---|---|
| | ajuba |
| | rgd1559714 |
| | sh2b2 |
| | sytl3 |
| | dnai2 |
| | bst1 |
| | il2rg |
| | pou1f1 |
| | gdf7 |
| | pld5 |
| | crb1 |
| | ccdc114 |
| | cdca7 |
| | timd2 |
| | unc13d |
| | loc689927 |
| | crygs |
| | rgd1565655 |
| | rrh |
| | spp1 |
| | def6 |
| | gja6 |
| | vom1r88 |
| | lyzl6 |
| | cmah |
| | txndc2 |
| | mirlet7d |
| | cb707485 |
| | olr1667 |
| | testin |
| | cyp2c24 |
| | fgg |

#### Supplementary table 6 : Conservation of human AD transcriptome with TgF344-AD rat transcriptome (Females)

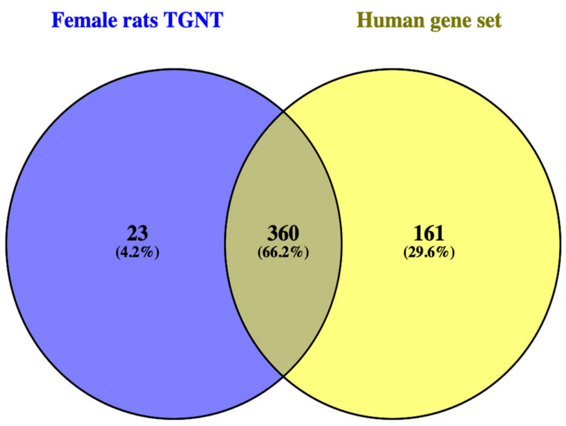

| Males rat TGNT | AGORA dataset for males | Common genes |
|---|---|---|
| psen1 | ifit3 | psen1 |
| app | syne4 | app |
| gfap | efcab10 | gfap |
| gpnmb | lrrc46 | gpnmb |
| gpr88 | odf3b | gpr88 |
| rgs9 | cp | rgs9 |
| six3 | drc1 | six3 |
| adora2a | myo7a | adora2a |
| lhx8 | tgfbi | lhx8 |
| gpr6 | nxnl1 | gpr6 |
| ecel1 | pqlc3 | ecel1 |
| sh3rf2 | aldh1l2 | sh3rf2 |
| akap12 | enkur | akap12 |
| syndig1l | olfm4 | syndig1l |
| slc22a3 | rgs1 | slc22a3 |
| drd1 | rsph1 | drd1 |
| foxp2 | cd68 | foxp2 |
| slc4a5 | sik1 | slc4a5 |
| h19 | erich2 | h19 |
| igf2 | mfap2 | igf2 |
| folr1 | efcab1 | folr1 |
| tmem27 | cdc153 | tmem27 |
| col5a1 | tekt2 | col5a1 |

| | | |
|---|---|---|
| tmem72 | fbxl13 | tmem72 |
| penk | ciita | penk |
| fcgr2b | ccdc146 | fcgr2b |
| tnni3 | prc1 | tnni3 |
| ace | impg2 | ace |
| mvd | tyrp1 | mvd |
| scn4b | ifltd1 | scn4b |
| kb15 | hist1h2bd | kb15 |
| cd22 | dydc2 | cd22 |
| f5 | igfbpl1 | f5 |
| drd2 | gpr139 | drd2 |
| sbk2 | fam154b | sbk2 |
| kl | kcnk16 | kl |
| sulf1 | baat | sulf1 |
| itgb6 | mrap | itgb6 |
| cdh3 | sytl3 | cdh3 |
| lect1 | scube3 | lect1 |
| cd74 | capsl | cd74 |
| rgd1559696 | bcl2l15 | rgd1559696 |
| mfrp | ccdc114 | mfrp |
| slc5a7 | cdh3 | slc5a7 |
| enpp2 | calcr | enpp2 |
| itpripl1 | slc7a9 | itpripl1 |
| ppp1r1b | ampd1 | ppp1r1b |
| trpv4 | ttc25 | trpv4 |
| hist1h2bo | daw1 | hist1h2bo |
| scgb1c1 | prrg4 | scgb1c1 |
| samd3 | slc44a4 | samd3 |
| col3a1 | guca1b | col3a1 |
| igfbpl1 | lrrc34 | igfbpl1 |
| inmt | trpv4 | inmt |
| krt71 | ttc29 | krt71 |
| tac1 | slc22a3 | tac1 |
| kcne2 | mpo | kcne2 |
| abca4 | rib1 | abca4 |
| zic1 | bbox1 | zic1 |
| calcr | ccdc40 | calcr |
| adipoq | pkhd1l1 | adipoq |
| col1a2 | mpzl2 | col1a2 |

| | | |
|---|---|---|
| lhx1 | azgp1 | lhx1 |
| naglt1 | pglyrp1 | naglt1 |
| ret | ace | ret |
| pappa | galr3 | pappa |
| kcnj13 | ak7 | kcnj13 |
| h1foo | ly9 | h1foo |
| jph2 | enpp2 | jph2 |
| slco1a5 | tmem212 | slco1a5 |
| pdzd2 | krt8 | pdzd2 |
| gapdh-ps1 | slc14a2 | gapdh-ps1 |
| sult1c3 | cd5 | sult1c3 |
| htr1d | lbp | htr1d |
| myl6 | dnah1 | myl6 |
| leap2 | aurkb | leap2 |
| ogn | pla2g5 | ogn |
| nexn | itgb6 | nexn |
| mapk4 | spag6 | mapk4 |
| tbc1d10c | aipl1 | tbc1d10c |
| rmrp | dynlrb2 | rmrp |
| ttr | mlf1 | ttr |
| slc13a4 | abcb11 | slc13a4 |
| lsp1 | kcne2 | lsp1 |
| tex15 | iqcg | tex15 |
| cldn2 | kcnq1 | cldn2 |
| rn28s | slc16a8 | rn28s |
| rasd2 | ahsp | rasd2 |
| slc18a3 | wdr63 | slc18a3 |
| vwce | pdc | vwce |
| otx2 | slpi | otx2 |
| trh | dcdc5 | trh |
| pla2g5 | akap14 | pla2g5 |
| asic4 | corin | asic4 |
| cd68 | clec7a | cd68 |
| slc35d3 | ubd | slc35d3 |
| neu2 | krt18 | neu2 |
| rgd1561102 | tex28 | rgd1561102 |
| loc287004 | lrit1 | loc287004 |
| ccnb1ip1 | inmt | ccnb1ip1 |
| prlr | ttll10 | prlr |

| | | |
|---|---|---|
| trat1 | hoxb5 | trat1 |
| irx3 | rbp3 | irx3 |
| arl11 | best2 | arl11 |
| capg | c1qtnf3 | capg |
| stra6 | tnmd | stra6 |
| lepr | agr3 | lepr |
| hnf4a | col8a1 | hnf4a |
| atp6ap1l | irx4 | atp6ap1l |
| fgf1 | sstr5 | fgf1 |
| cmtm1 | mc3r | cmtm1 |
| clic6 | adipoq | clic6 |
| runx1 | gata6 | runx1 |
| pcp4l1 | lrrc18 | pcp4l1 |
| col5a2 | mak | col5a2 |
| col7a1 | mylk2 | col7a1 |
| ccl9 | selp | ccl9 |
| cartpt | prph2 | cartpt |
| isl1 | cr2 | isl1 |
| rgd1561795 | tmem232 | rgd1561795 |
| gbx2 | lgals5 | gbx2 |
| cd200r1 | fabp12 | cd200r1 |
| sostdc1 | rbp7 | sostdc1 |
| dazl | agtr2 | dazl |
| slc22a8 | abca4 | slc22a8 |
| meis2 | pih1d3 | meis2 |
| itgb1bp2 | slc22a2 | itgb1bp2 |
| mpz | pon3 | mpz |
| tssk3 | aqp1 | tssk3 |
| hp | steap1 | hp |
| pcp4 | uncx | pcp4 |
| loc691083 | gdf7 | loc691083 |
| tcf21 | padi4 | tcf21 |
| kcnq1 | pde6h | kcnq1 |
| glycam1 | lpo | glycam1 |
| pon3 | crygs | pon3 |
| rbm47 | pon1 | rbm47 |
| gpr52 | hormad2 | gpr52 |
| fam160a1 | zscan10 | fam160a1 |
| pde7b | tmem27 | pde7b |

| | | |
|---|---|---|
| tbx6 | pax3 | tbx6 |
| ltbp2 | efhb | ltbp2 |
| tmem184a | ccdc60 | tmem184a |
| frem1 | trim6 | frem1 |
| il13ra2 | lmx1a | il13ra2 |
| mpzl2 | otx2 | mpzl2 |
| ptprq | il13ra2 | ptprq |
| itk | impg1 | itk |
| clul1 | cldn2 | clul1 |
| melk | bhmt2 | melk |
| ccl3 | cd8b | ccl3 |
| tnnt2 | kcnj13 | tnnt2 |
| rgd1561161 | tmem184a | rgd1561161 |
| lgals5 | folr1 | lgals5 |
| spetex-2a | mfrp | spetex-2a |
| fetub | sox14 | fetub |
| rarb | lilrb4 | rarb |
| aqp1 | clic6 | aqp1 |
| klk4 | kl | klk4 |
| trpv5 | slc4a5 | trpv5 |
| pon1 | ttr | pon1 |
| ncf2 | tmem72 | ncf2 |
| kremen1 | tulp1 | kremen1 |
| enkur | sostdc1 | enkur |
| hes3 | nrg4 | hes3 |
| pdp1 | gngt1 | pdp1 |
| igfbp2 | krt19 | igfbp2 |
| slc4a2 | f5 | slc4a2 |
| mdfic | cxcl11 | mdfic |
| lhx5 | tcf21 | lhx5 |
| steap1 | ttll2 | steap1 |
| pde10a | six6 | pde10a |
| scube3 | hnrnpu | scube3 |
| il20ra | psen1 | il20ra |
| iqgap3 | app | iqgap3 |
| gbx1 | gfap | gbx1 |
| camp | gpnmb | camp |
| slc16a12 | gpr88 | slc16a12 |
| vav1 | rgs9 | vav1 |

| | | |
|---|---|---|
| s100g | six3 | s100g |
| mxd3 | adora2a | mxd3 |
| dmkn | lhx8 | dmkn |
| cdr2 | gpr6 | cdr2 |
| kcnv2 | ecel1 | kcnv2 |
| lmx1a | sh3rf2 | lmx1a |
| kcnh4 | akap12 | kcnh4 |
| brs3 | syndig1l | brs3 |
| gucy2f | slc22a3 | gucy2f |
| naa11 | drd1 | naa11 |
| pih1d3 | foxp2 | pih1d3 |
| mcoln3 | slc4a5 | mcoln3 |
| crabp1 | h19 | crabp1 |
| loc100363193 | igf2 | loc100363193 |
| mc3r | folr1 | mc3r |
| cdx4 | tmem27 | cdx4 |
| gucy1a3 | col5a1 | gucy1a3 |
| loc654482 | tmem72 | loc654482 |
| cdk1 | penk | cdk1 |
| zscan10 | fcgr2b | zscan10 |
| veph1 | tnni3 | veph1 |
| ngef | ace | ngef |
| adra2b | mvd | adra2b |
| fbxo39 | scn4b | fbxo39 |
| mcoln2 | kb15 | mcoln2 |
| aldh1a3 | cd22 | aldh1a3 |
| loc102548399 | f5 | loc102548399 |
| slc16a8 | drd2 | slc16a8 |
| tnn | sbk2 | tnn |
| gnal | kl | gnal |
| trpm3 | sulf1 | trpm3 |
| lag3 | itgb6 | lag3 |
| capsl | cdh3 | capsl |
| hcls1 | lect1 | hcls1 |
| cdhr1 | cd74 | cdhr1 |
| avpr2 | rgd1559696 | avpr2 |
| fam187a | mfrp | fam187a |
| rgd1563692 | slc5a7 | rgd1563692 |
| slc2a12 | enpp2 | slc2a12 |

| | | |
|---|---|---|
| pcp2 | itpripl1 | pcp2 |
| lrrk2 | ppp1r1b | lrrk2 |
| bhmt2 | trpv4 | bhmt2 |
| tns4 | hist1h2bo | tns4 |
| igfals | scgb1c1 | igfals |
| pitx2 | samd3 | pitx2 |
| ido1 | col3a1 | ido1 |
| insrr | igfbpl1 | insrr |
| coch | inmt | coch |
| tmem196 | krt71 | tmem196 |
| nr5a1 | tac1 | nr5a1 |
| pde1b | kcne2 | pde1b |
| dclk3 | abca4 | dclk3 |
| rgd1308065 | zic1 | rgd1308065 |
| col18a1 | calcr | col18a1 |
| adamtsl4 | adipoq | adamtsl4 |
| has1 | col1a2 | has1 |
| fhod3 | lhx1 | fhod3 |
| ugt1a6 | naglt1 | ugt1a6 |
| cdkn1c | ret | cdkn1c |
| bcmo1 | pappa | bcmo1 |
| runx3 | kcnj13 | runx3 |
| lilrb4 | h1foo | lilrb4 |
| irx6 | jph2 | irx6 |
| spock3 | slco1a5 | spock3 |
| rbp4 | pdzd2 | rbp4 |
| krt42 | gapdh-ps1 | krt42 |
| retn | sult1c3 | retn |
| bcl2l11 | htr1d | bcl2l11 |
| c1qtnf3 | myl6 | c1qtnf3 |
| prkar2b | leap2 | prkar2b |
| atp8b1 | ogn | atp8b1 |
| acot12 | nexn | acot12 |
| rasgrp2 | mapk4 | rasgrp2 |
| arpp21 | tbc1d10c | arpp21 |
| fam46a | rmrp | fam46a |
| atp4a | ttr | atp4a |
| prdm1 | slc13a4 | prdm1 |
| mgp | lsp1 | mgp |

| | | |
|---|---|---|
| kif23 | tex15 | kif23 |
| reck | cldn2 | reck |
| rt1-da | rn28s | rt1-da |
| hs3st3a1 | rasd2 | hs3st3a1 |
| lrrc18 | slc18a3 | lrrc18 |
| mybl2 | vwce | mybl2 |
| ces1d | otx2 | ces1d |
| foxj1 | trh | foxj1 |
| xcl1 | pla2g5 | xcl1 |
| slpil2 | asic4 | slpil2 |
| arl4d | cd68 | arl4d |
| krt85 | slc35d3 | krt85 |
| pou4f3 | neu2 | pou4f3 |
| hist1h2aa | rgd1561102 | hist1h2aa |
| crb3 | loc287004 | crb3 |
| hist1h2ak | ccnb1ip1 | hist1h2ak |
| aldoart2 | prlr | aldoart2 |
| vom1r16 | trat1 | vom1r16 |
| msx1 | irx3 | msx1 |
| ttc26 | arl11 | ttc26 |
| serpinc1 | capg | serpinc1 |
| hist1h1b | stra6 | hist1h1b |
| rnf152 | lepr | rnf152 |
| arhgap30 | hnf4a | arhgap30 |
| plek | atp6ap1l | plek |
| prkch | fgf1 | prkch |
| mdk | cmtm1 | mdk |
| sgcg | clic6 | sgcg |
| rgd1560608 | runx1 | rgd1560608 |
| slc10a4 | pcp4l1 | slc10a4 |
| ddc8 | col5a2 | ddc8 |
| c3ar1 | col7a1 | c3ar1 |
| a2m | ccl9 | a2m |
| efcab1 | cartpt | efcab1 |
| ccdc40 | isl1 | ccdc40 |
| mak | rgd1561795 | mak |
| nmu | gbx2 | nmu |
| tmem170a | cd200r1 | tmem170a |
| loc100361092 | sostdc1 | loc100361092 |

| | | |
|---|---|---|
| klk7 | dazl | klk7 |
| nkx2-1 | slc22a8 | nkx2-1 |
| zic2 | meis2 | zic2 |
| cxcl13 | itgb1bp2 | cxcl13 |
| loxl4 | mpz | loxl4 |
| rrm2 | tssk3 | rrm2 |
| pabpc2 | hp | pabpc2 |
| rgd1303271 | pcp4 | rgd1303271 |
| slc14a2 | loc691083 | slc14a2 |
| slc5a12 | tcf21 | slc5a12 |
| reg3a | kcnq1 | reg3a |
| top2a | glycam1 | top2a |
| rt1-ba | pon3 | rt1-ba |
| bmp2 | rbm47 | bmp2 |
| elovl7 | gpr52 | elovl7 |
| unc13c | fam160a1 | unc13c |
| uncx | pde7b | uncx |
| itga5 | tbx6 | itga5 |
| foxb1 | ltbp2 | foxb1 |
| klra5 | tmem184a | klra5 |
| ces2 | frem1 | ces2 |
| fam71f1 | il13ra2 | fam71f1 |
| pnpla1 | mpzl2 | pnpla1 |
| pdzk1ip1 | ptprq | pdzk1ip1 |
| nepn | itk | nepn |
| spink5 | clul1 | spink5 |
| rgd1561870 | melk | rgd1561870 |
| loc689713 | ccl3 | loc689713 |
| slc16a2 | tnnt2 | slc16a2 |
| tshr | rgd1561161 | tshr |
| prkag3 | lgals5 | prkag3 |
| slc7a12 | spetex-2a | slc7a12 |
| sct | fetub | sct |
| rgd1561958 | rarb | rgd1561958 |
| fgf20 | aqp1 | fgf20 |
| klk1 | klk4 | klk1 |
| adcy5 | trpv5 | adcy5 |
| cpne5 | pon1 | cpne5 |
| slc12a7 | ncf2 | slc12a7 |

| | | |
|---|---|---|
| st6galnac2 | kremen1 | st6galnac2 |
| akp3 | enkur | akp3 |
| tc2n | hes3 | tc2n |
| loxl2 | pdp1 | loxl2 |
| kcna5 | igfbp2 | kcna5 |
| cmtm8 | slc4a2 | cmtm8 |
| fam109b | mdfic | fam109b |
| cyp11b3 | lhx5 | cyp11b3 |
| fyb | steap1 | fyb |
| ino80d | pde10a | ino80d |
| chad | scube3 | chad |
| cenpf | il20ra | cenpf |
| car13 | iqgap3 | car13 |
| fbp1 | gbx1 | fbp1 |
| vgll2 | camp | vgll2 |
| ldhal6b | slc16a12 | ldhal6b |
| htr6 | vav1 | htr6 |
| kiss1 | s100g | kiss1 |
| ms4a14 | mxd3 | ms4a14 |
| krt18 | dmkn | krt18 |
| prima1 | cdr2 | prima1 |
| gsc2 | kcnv2 | gsc2 |
| aox3 | lmx1a | aox3 |
| chrna9 | kcnh4 | chrna9 |
| otc | brs3 | otc |
| synpo2 | gucy2f | synpo2 |
| slc23a3 | naa11 | slc23a3 |
| sim1 | pih1d3 | sim1 |
| ifna4 | mcoln3 | ifna4 |
| spag11c | crabp1 | spag11c |
| mlf1 | loc100363193 | mlf1 |
| cpn1 | mc3r | cpn1 |
| lenep | cdx4 | lenep |
| rgd1306625 | gucy1a3 | rgd1306625 |
| rhod | loc654482 | rhod |
| gng7 | cdk1 | gng7 |
| lhfpl1 | zscan10 | lhfpl1 |
| ptpn5 | veph1 | ptpn5 |
| sv2c | ngef | sv2c |

| | | |
|---|---|---|
| rgd1309028 | adra2b | rgd1309028 |
| mospd1 | fbxo39 | mospd1 |
| bcl3 | mcoln2 | bcl3 |
| mmp2 | aldh1a3 | mmp2 |
| slc4a11 | loc102548399 | slc4a11 |
| krt75 | slc16a8 | krt75 |
| rgd1562658 | tnn | rgd1562658 |
| loc100910620 | gnal | loc100910620 |
| fmo2 | trpm3 | fmo2 |
| mael | lag3 | mael |
| serpine1 | capsl | serpine1 |
| olr200 | hcls1 | olr200 |
| txk | cdhr1 | txk |
| gck | avpr2 | gck |
| fam222a | fam187a | fam222a |
| clec4e | rgd1563692 | clec4e |
| cgnl1 | slc2a12 | cgnl1 |
| f13a1 | pcp2 | f13a1 |
| six1 | lrrk2 | six1 |
| smim22 | bhmt2 | smim22 |
| lif | tns4 | lif |
| hist2h4 | igfals | hist2h4 |
| slc5a5 | pitx2 | slc5a5 |
| bhmt | ido1 | bhmt |
| itpr3 | insrr | itpr3 |
| efhb | coch | efhb |
| ccl4 | tmem196 | ccl4 |
| art3 | nr5a1 | art3 |
| loc171161 | pde1b | loc171161 |
| rgd1305627 | dclk3 | rgd1305627 |
| nags | rgd1308065 | nags |
| myo1f | col18a1 | myo1f |
| dlx5 | adamtsl4 | dlx5 |
| wdr16 | has1 | wdr16 |
| grm4 | fhod3 | grm4 |
| corin | ugt1a6 | corin |
| phactr2 | cdkn1c | phactr2 |
| kcnk2 | bcmo1 | kcnk2 |
| kcnn4 | runx3 | kcnn4 |

| | | |
|---|---|---|
| gulp1 | lilrb4 | gulp1 |
| wdr63 | irx6 | wdr63 |
| glb1l | spock3 | glb1l |
| pde6b | rbp4 | pde6b |
| rab38 | krt42 | rab38 |
| gpr153 | retn | gpr153 |
| ccdc166 | bcl2l11 | ccdc166 |
| gypc | c1qtnf3 | gypc |
| klhl1 | prkar2b | klhl1 |
| msc | atp8b1 | msc |
| ifltd1 | acot12 | ifltd1 |
| baiap2l1 | rasgrp2 | baiap2l1 |
| cd72 | arpp21 | cd72 |
| mme | fam46a | mme |
| actn3 | atp4a | actn3 |
| sfrp1 | prdm1 | sfrp1 |
| car14 | mgp | car14 |
| rt1-ce3 | kif23 | rt1-ce3 |
| slc31a1 | reck | slc31a1 |
| rab11fip1 | rt1-da | rab11fip1 |
| npr3 | hs3st3a1 | npr3 |
| clec2d | lrrc18 | clec2d |
| tnxa-ps1 | mybl2 | tnxa-ps1 |
| slc16a3 | ces1d | slc16a3 |
| best3 | foxj1 | best3 |
| spag8 | xcl1 | spag8 |
| iqcg | slpil2 | iqcg |
| hist2h3c2 | arl4d | hist2h3c2 |
| ube2l6 | krt85 | ube2l6 |
| mcm10 | pou4f3 | mcm10 |
| stk32b | hist1h2aa | stk32b |
| entpd3 | crb3 | entpd3 |
| tpm2 | hist1h2ak | tpm2 |
| dnali1 | aldoart2 | dnali1 |
| egfr | vom1r16 | egfr |
| tinagl1 | msx1 | tinagl1 |
| zic3 | ttc26 | zic3 |
| cobl | serpinc1 | cobl |
| ect2 | hist1h1b | ect2 |

| | | |
|---|---|---|
| exo1 | rnf152 | exo1 |
| stat4 | arhgap30 | stat4 |
| lysmd3 | plek | lysmd3 |
| tgm3 | prkch | tgm3 |
| fscn2 | mdk | fscn2 |
| ccdc60 | sgcg | ccdc60 |
| rt1-db1 | rgd1560608 | rt1-db1 |
| chrdl2 | slc10a4 | chrdl2 |
| hnf1b | ddc8 | hnf1b |
| rt1-ce15 | c3ar1 | rt1-ce15 |
| clec5a | a2m | clec5a |
| il18bp | efcab1 | il18bp |
| ajuba | ccdc40 | ajuba |
| rgd1559714 | mak | rgd1559714 |
| sh2b2 | nmu | sh2b2 |
| sytl3 | tmem170a | sytl3 |
| dnai2 | loc100361092 | dnai2 |
| bst1 | klk7 | bst1 |
| il2rg | nkx2-1 | il2rg |
| pou1f1 | zic2 | pou1f1 |
| gdf7 | cxcl13 | gdf7 |
| pld5 | loxl4 | pld5 |
| crb1 | rrm2 | crb1 |
| ccdc114 | pabpc2 | ccdc114 |
| cdca7 | rgd1303271 | cdca7 |
| timd2 | slc14a2 | timd2 |
| unc13d | slc5a12 | unc13d |
| loc689927 | reg3a | loc689927 |
| crygs | top2a | crygs |
| rgd1565655 | rt1-ba | rgd1565655 |
| rrh | bmp2 | rrh |
| spp1 | elovl7 | spp1 |
| def6 | unc13c | def6 |
| gja6 | uncx | gja6 |
| vom1r88 | itga5 | vom1r88 |
| lyzl6 | foxb1 | lyzl6 |
| cmah | klra5 | cmah |
| txndc2 | ces2 | txndc2 |
| mirlet7d | fam71f1 | mirlet7d |

| | | |
|---|---|---|
| cb707485 | pnpla1 | cb707485 |
| olr1667 | pdzk1ip1 | olr1667 |
| testin | nepn | testin |
| cyp2c24 | spink5 | cyp2c24 |
| fgg | rgd1561870 | fgg |
| lrrc52 | loc689713 | agtr2 |
| odf3l2 | slc16a2 | |
| dmrtc1c1 | tshr | |
| krt86 | prkag3 | |
| cd69 | slc7a12 | |
| agtr2 | sct | |
| cryge | rgd1561958 | |
| dcm5 | fgf20 | |
| oas1h | klk1 | |
| ugt1a9 | adcy5 | |
| tcp10b | cpne5 | |
| cldn22 | slc12a7 | |
| olr1653 | st6galnac2 | |
| sell | akp3 | |
| adh7 | tc2n | |
| mirlet7c-2 | loxl2 | |
| mir3572 | kcna5 | |
| olr1869 | cmtm8 | |
| efhc1 | fam109b | |
| pbx3 | cyp11b3 | |
| gfi1b | fyb | |
| gykl1 | ino80d | |
| adamts19 | chad | |
| ropn1l | cenpf | |
| lrr1 | car13 | |
| otx1 | fbp1 | |
| hepacam2 | vgll2 | |
| krt34 | ldhal6b | |
| olr387 | htr6 | |
| epn3 | kiss1 | |
| cyp11a1 | ms4a14 | |
| slamf6 | krt18 | |
| crygf | prima1 | |
| krt1 | gsc2 | |

| | |
|---|---|
| foxc2 | aox3 |
| dab2 | chrna9 |
| mrgprf | otc |
| troap | synpo2 |
| evpl | slc23a3 |
| prelp | sim1 |
| igsf9 | ifna4 |
| mir421 | spag11c |
| adcy7 | mlf1 |
| vwa5b1 | cpn1 |
| pof1b | lenep |
| rdh7 | rgd1306625 |
| fam83a | rhod |
| rgd1561778 | gng7 |
| tead2 | lhfpl1 |
| cox8b | ptpn5 |
| llgl2 | sv2c |
| dpp4 | rgd1309028 |
| scara5 | mospd1 |
| kcna10 | bcl3 |
| rgd1563104 | mmp2 |
| trim40 | slc4a11 |
| ugt1a5 | krt75 |
| apobec1 | rgd1562658 |
| krt16 | loc100910620 |
| rin3 | fmo2 |
| s100a5 | mael |
| kctd8 | serpine1 |
| zim1 | olr200 |
| sp8 | txk |
| upk1b | gck |
| nlrp4 | fam222a |
| iqub | clec4e |
| vom2r60 | cgnl1 |
| lipogenin | f13a1 |
| zmynd10 | six1 |
| pax8 | smim22 |
| gpx2 | lif |
| fgf19 | hist2h4 |

| | |
|---|---|
| epha1 | slc5a5 |
| il22ra1 | bhmt |
| kcnmb3 | itpr3 |
| magea4 | efhb |
| olr176 | ccl4 |
| | art3 |
| | loc171161 |
| | rgd1305627 |
| | nags |
| | myo1f |
| | dlx5 |
| | wdr16 |
| | grm4 |
| | corin |
| | phactr2 |
| | kcnk2 |
| | kcnn4 |
| | gulp1 |
| | wdr63 |
| | glb1l |
| | pde6b |
| | rab38 |
| | gpr153 |
| | ccdc166 |
| | gypc |
| | klhl1 |
| | msc |
| | ifltd1 |
| | baiap2l1 |
| | cd72 |
| | mme |
| | actn3 |
| | sfrp1 |
| | car14 |
| | rt1-ce3 |
| | slc31a1 |
| | rab11fip1 |
| | npr3 |
| | clec2d |

| | |
|---|---|
| | tnxa-ps1 |
| | slc16a3 |
| | best3 |
| | spag8 |
| | iqcg |
| | hist2h3c2 |
| | ube2l6 |
| | mcm10 |
| | stk32b |
| | entpd3 |
| | tpm2 |
| | dnali1 |
| | egfr |
| | tinagl1 |
| | zic3 |
| | cobl |
| | ect2 |
| | exo1 |
| | stat4 |
| | lysmd3 |
| | tgm3 |
| | fscn2 |
| | ccdc60 |
| | rt1-db1 |
| | chrdl2 |
| | hnf1b |
| | rt1-ce15 |
| | clec5a |
| | il18bp |
| | ajuba |
| | rgd1559714 |
| | sh2b2 |
| | sytl3 |
| | dnai2 |
| | bst1 |
| | il2rg |
| | pou1f1 |
| | gdf7 |
| | pld5 |

| | |
|---|---|
| | crb1 |
| | ccdc114 |
| | cdca7 |
| | timd2 |
| | unc13d |
| | loc689927 |
| | crygs |
| | rgd1565655 |
| | rrh |
| | spp1 |
| | def6 |
| | gja6 |
| | vom1r88 |
| | lyzl6 |
| | cmah |
| | txndc2 |
| | mirlet7d |
| | cb707485 |
| | olr1667 |
| | testin |
| | cyp2c24 |
| | fgg |

#### Supplementary table 7 : List of genes in TGTR upregulated pathways (males)

| Pathway | Constituent Genes (Representative) | Benefit in AD Context | Key References |
|---|---|---|---|
| **Protein synthesis & mitochondria** (TZ-Up) | MRPL11, MRPL16, MRPS17, RPL10, RPL13A, RPS6, RPS27A | Restoration of mitochondrial translation and ribosomal function may improve neuronal bioenergetics and synaptic resilience. Mitochondrial dysfunction is an early event in AD; supporting translational capacity may counteract neurodegeneration. | Swerdlow 2018 Nat Rev Neurosci; Wang et al., 2020 Mol Neurodegener |
| **Immune signaling & homeostatic** | STAT1, IFNAR1, IL1R1, TNFRSF19, SOCS1 | Moderate immune surveillance enhancement may promote protective microglial responses and debris clearance. Early | Heneka et al., 2015 Lancet Neurol; Deczkowska et |

| | | | |
|---|---|---|---|
| **surveillance** (TZ-Up) | | controlled immune activation can be protective before chronic inflammatory shift. | al., 2018 Nat Immunol |
| **Antigen presentation & complement** (TZ-Up) | RT1-DB1, RT1-BB, B2M, C1QC, C1S, CFB | Enhanced complement and antigen pathways may support plaque tagging and clearance. Complement-mediated synaptic pruning is pathological when excessive but beneficial in regulated contexts. | Hong et al., 2016 Science; Shi et al., 2017 Nat Neurosci |
| **Microglia activation & phagolysosome** (TZ-Up) | TREM2, LGMN, CD14, FCGR1A, P2RY12 | Increased phagolysosomal activity may enhance Aβ clearance and barrier formation around plaques. Disease-associated microglia (DAM) can be protective in early AD. | Keren-Shaul et al., 2017 Cell; Yuan et al., 2016 Nat Med |
| **ECM / cytoskeleton / vascular-lipid** (TZ-Up) | COL1A1, COL3A1, VCAM1, ITGB8, MYLK | ECM remodeling and vascular stabilization may support blood-brain barrier integrity and structural resilience. | Zlokovic 2011 Nat Rev Neurosci |

#### Supplementary table 8 : List of genes in TGTR downregulated pathways (males)

| Pathway | Constituent Genes (Representative) | Benefit in AD Context | Key References |
|---|---|---|---|
| **RNA processing (APP/BACE1)** (TZ down) | SF3B2, SRSF2, SNRNP70, PRPF3, DDX5, HNRNPC, RBM22, THOC1 | Spliceosome and RNA-binding proteins regulate stability of APP and BACE1 transcripts. Downregulation may reduce amyloidogenic mRNA stabilization and lower Aβ production. | Kang et al., 2011 Nat Neurosci; Faghihi et al., 2008 Nat Med |
| **Purinergic / GPCR neuroinflammatory coupling** (TZ down) | P2RX2, P2RY1, P2RY2, ADRA1B, HTR6, DRD5, TACR3 | Purinergic and neuromodulatory GPCR signaling drives microglial inflammatory cascades and excitotoxicity. Suppression may reduce neuroinflammation. | Burnstock 2017 Pharmacol Rev; Illes et al., 2019 Nat Rev Drug Discov |
| **Excitatory neurotransmission & ion channels** (TZ down) | GRIA4, GRIN2D, CACNA1A, CHRNA2, SLC17A7, SLC6A1 | Reduced excitatory drive may protect against excitotoxic neuronal injury and synaptic degeneration. | Hynd et al., 2004 Nat Rev Neurosci |

| | | | |
|---|---|---|---|
| **Proteostasis & organelle stress (V-ATPase / HSP)** (TZ down) | ATP6V1C1, ATP6V1D, HSPA1A, HSPA8 | Decreased maladaptive stress-response activation may indicate reduced ER/mitochondrial stress burden. | Scheper & Hoozemans 2015 Cell Mol Life Sci |
| **Endosomal trafficking & APP processing** (TZ down) | CLTA, CLTC, AP2A2, STX3, STXBP1, VAMP2 | Reduced endocytosis and vesicular trafficking may decrease APP–BACE1 convergence in endosomes, limiting amyloidogenic processing. | Rajendran et al., 2006 Science; Small & Gandy 2006 Neuron |

#### Supplementary table 9 : List of genes in TGTR upregulated pathways (females)

| Pathway | Representative Constituent Genes | Benefit in AD Context | Key References |
|---|---|---|---|
| **Cell survival & pro-growth signaling** | AKT2, AKT3, BCL2, CREB5, ERBB4, PRKCA | Enhances neuronal survival signaling, counteracting apoptosis and synaptic vulnerability seen in female AD models. AKT/CREB pathways support synaptic resilience. | Pei et al., 2003 J Neurosci; Talantova et al., 2013 Nature |
| **GPCR & $Ca^{2+}$ signaling** | ITPR1, ITPR2, CACNB4, ADRA1B, PLCB1 | Proper calcium homeostasis is essential for synaptic transmission; restoring regulated $Ca^{2+}$ signaling prevents dysregulated excitotoxic cascades. | LaFerla 2002 Nat Rev Neurosci |
| **cAMP / cGMP second messenger signaling** | ADCY1, ADCY8, GUCY1A1, GUCY1B1, PRKG1 | cAMP/cGMP signaling enhances synaptic plasticity and memory formation; PDE modulation is protective in AD models. | Puzzo et al., 2009 J Neurosci |
| **Synaptic signaling & plasticity** | GRIA1, GRIN1, CAMK2A | Reinforces synaptic strength and long-term potentiation (LTP), counteracting female-biased synaptic vulnerability. | Selkoe 2002 Science |
| **Cell cycle & DNA maintenance** | BRCA1, CDK2, PPP2R5A | Restoration of controlled cell cycle and genomic maintenance reduces aberrant neuronal cell-cycle re-entry seen in AD. | Yang et al., 2003 J Neurosci |

| | | | |
|---|---|---|---|
| **Dendrite growth & cytoskeletal organization** | PAK1, PAK2, ITGA5, FN1 | Supports dendritic spine stability and structural plasticity. | Zhao et al., 2006 Nat Neurosci |

#### Supplementary table 10 : List of genes in TGTR downregulated pathways (females)

| Pathway | Representative Constituent Genes | Benefit in AD Context | References |
|---|---|---|---|
| **Metabolic stress & mitochondrial dysfunction** | NDUFA9, NDUFS1, SDHA, COX5B, ATP5F1A | Chronic mitochondrial stress accelerates ROS production and neuronal injury; suppression suggests normalization of metabolic burden. | Swerdlow, 2018 Nat Rev Neurosci |
| **Proteasome / protein degradation stress** | PSME1, PSMA1, PSMB5, UBE2L6 | Excess UPS activation reflects proteotoxic stress; reduction may indicate decreased misfolded protein load. | Tai & Schuman, 2008 Nat Rev Neurosci |
| **Kinase activation & MAPK stress signaling** | MAPK9 (JNK2), MAPK14 (p38), MAP2K3 | p38/JNK pathways promote tau phosphorylation and neuroinflammation; suppression is neuroprotective. | Hensley et al., 1999 J Neurochem |
| **ECM remodeling & structural reorganization** | COL1A1, COL4A1, LAMA1, ITGA5 | Excess ECM remodeling correlates with BBB dysfunction and neuroinflammation; moderation restores structural balance. | Zlokovic, 2011 Nat Rev Neurosci |
| **Apoptosis & cell death signaling** | CASP1, FAS, BAD, BCL2L11 | Suppression reduces neuronal apoptosis and inflammatory cell death. | Yuan & Yankner, 2000 Nature |
| **APP processing & amyloidogenic machinery** | BACE1, PSEN1, APBB1, APP | Direct reduction of amyloidogenic processing lowers Aβ generation and plaque burden. | Vassar et al., 1999 Science |

#### Supplementary table 11: 2-way ANOVA of NeuN % area of dorsal hippocampus

| Source of Variation | % of total variation | P value | P value summary | Significant? | |
|---|---|---|---|---|---|
| Interaction | 0.08851 | 0.8434 | ns | No | |
| Genotype | 0.1225 | 0.8163 | ns | No | |
| Treatment | 3.400 | 0.2248 | ns | No | |
| ANOVA table | SS (Type III) | DF | MS | F (DFn, DFd) | P value |
| Interaction | 0.01530 | 1 | 0.01530 | F (1, 43) = 0.03948 | P=0.8434 |
| Genotype | 0.02118 | 1 | 0.02118 | F (1, 43) = 0.05464 | P=0.8163 |
| Treatment | 0.5878 | 1 | 0.5878 | F (1, 43) = 1.517 | P=0.2248 |
| Residual | 16.67 | 43 | 0.3876 | | |

#### Supplementary table 12: 2-way ANOVA of NeuN % area of CA1

| Source of Variation | % of total variation | P value | P value summary | Significant? | |
|---|---|---|---|---|---|
| Interaction | 0.9866 | 0.5135 | ns | No | |
| Genotype | 1.132 | 0.4841 | ns | No | |
| Treatment | 0.03435 | 0.9027 | ns | No | |
| ANOVA table | SS (Type III) | DF | MS | F (DFn, DFd) | P value |
| Interaction | 0.09951 | 1 | 0.09951 | F (1, 43) = 0.4341 | P=0.5135 |
| Genotype | 0.1142 | 1 | 0.1142 | F (1, 43) = 0.4981 | P=0.4841 |
| Treatment | 0.003465 | 1 | 0.003465 | F (1, 43) = 0.01512 | P=0.9027 |
| Residual | 9.856 | 43 | 0.2292 | | |

#### Supplementary table 13: 2-way ANOVA of NeuN % area of CA3

| Source of Variation | % of total variation | P value | P value summary | Significant? | |
|---|---|---|---|---|---|
| Interaction | 1.615 | 0.3857 | ns | No | |
| Genotype | 0.04599 | 0.8831 | ns | No | |
| Treatment | 7.679 | 0.0627 | ns | No | |
| ANOVA table | SS (Type III) | DF | MS | F (DFn, DFd) | P value |
| Interaction | 0.3166 | 1 | 0.3166 | F (1, 43) = 0.7680 | P=0.3857 |
| Genotype | 0.009012 | 1 | 0.009012 | F (1, 43) = 0.02186 | P=0.8831 |
| Treatment | 1.505 | 1 | 1.505 | F (1, 43) = 3.651 | P=0.0627 |
| Residual | 17.72 | 43 | 0.4122 | | |

#### Supplementary table 14: 2-way ANOVA of NeuN % area of Hilar

| Source of Variation | % of total variation | P value | P value summary | Significant? | |
|---|---|---|---|---|---|
| Interaction | 0.4602 | 0.6521 | ns | No | |
| Genotype | 1.890 | 0.3628 | ns | No | |
| Treatment | 8.863 | 0.0531 | ns | No | |
| ANOVA table | SS (Type III) | DF | MS | F (DFn, DFd) | P value |
| Interaction | 114168 | 1 | 114168 | F (1, 40) = 0.2063 | P=0.6521 |
| Genotype | 468939 | 1 | 468939 | F (1, 40) = 0.8474 | P=0.3628 |
| Treatment | 2198797 | 1 | 2198797 | F (1, 40) = 3.973 | P=0.0531 |
| Residual | 22135742 | 40 | 553394 | | |

#### Supplementary table 15: 2-way ANOVA of NeuN % area of SB

| Source of Variation | % of total variation | P value | P value summary | Significant? | |
|---|---|---|---|---|---|
| Interaction | 2.616 | 0.2763 | ns | No | |
| Genotype | 0.4333 | 0.6558 | ns | No | |
| Treatment | 6.438 | 0.0909 | ns | No | |
| ANOVA table | SS (Type III) | DF | MS | F (DFn, DFd) | P value |
| Interaction | 0.5320 | 1 | 0.5320 | F (1, 42) = 1.217 | P=0.2763 |
| Genotype | 0.08814 | 1 | 0.08814 | F (1, 42) = 0.2016 | P=0.6558 |
| Treatment | 1.309 | 1 | 1.309 | F (1, 42) = 2.995 | P=0.0909 |
| Residual | 18.36 | 42 | 0.4372 | | |

#### Supplementary table 16: 2-way ANOVA of NET % area of CA1

| Source of Variation | % of total variation | P value | P value summary | Significant? | |
|---|---|---|---|---|---|
| Interaction | 7.141 | 0.0704 | ns | No | |
| Sex | 0.9115 | 0.5107 | ns | No | |
| Treatment | 3.967 | 0.1737 | ns | No | |
| ANOVA table | SS (Type III) | DF | MS | F (DFn, DFd) | P value |
| Interaction | 1.908 | 1 | 1.908 | F (1, 42) = 3.448 | P=0.0704 |
| Genotype | 0.2436 | 1 | 0.2436 | F (1, 42) = 0.4401 | P=0.5107 |
| Treatment | 1.060 | 1 | 1.060 | F (1, 42) = 1.915 | P=0.1737 |
| Residual | 23.25 | 42 | 0.5535 | | |

#### Supplementary table 17: 2-way ANOVA of NET % area of CA3

| Source of Variation | % of total variation | P value | P value summary | Significant? | |
|---|---|---|---|---|---|
| Interaction | 0.5889 | 0.6145 | ns | No | |
| Sex | 0.01275 | 0.9408 | ns | No | |
| Treatment | 3.448 | 0.2264 | ns | No | |
| ANOVA table | SS (Type III) | DF | MS | F (DFn, DFd) | P value |
| Interaction | 281376 | 1 | 281376 | F (1, 42) = 0.2574 | P=0.6145 |
| Sex | 6093 | 1 | 6093 | F (1, 42) = 0.005575 | P=0.9408 |
| Treatment | 1647456 | 1 | 1647456 | F (1, 42) = 1.507 | P=0.2264 |
| Residual | 45906049 | 42 | 1093001 | | |

#### Supplementary table 18: 2-way ANOVA of NET % area of SB

| Source of Variation | % of total variation | P value | P value summary | Significant? | |
|---|---|---|---|---|---|
| Interaction | 7.226 | 0.0792 | ns | No | |
| Sex | 23.58 | 0.32 | ns | No | |
| Treatment | 27.13 | 0.0818 | ns | No | |
| ANOVA table | SS (Type III) | DF | MS | F (DFn, DFd) | P value |
| Interaction | 3.484 | 1 | 3.484 | F (1, 21) = 3.403 | P=0.0792 |
| Sex | 11.37 | 1 | 11.37 | F (1, 21) = 11.11 | P=0.32 |
| Treatment | 13.08 | 1 | 13.08 | F (1, 21) = 12.78 | P=0.0818 |
| Residual | 21.50 | 21 | 1.024 | | |

**Supplementary table 19 : Common genes between LC dataset (Ehrenberg et al) and TgF344-AD rat DEGs (MALES)**

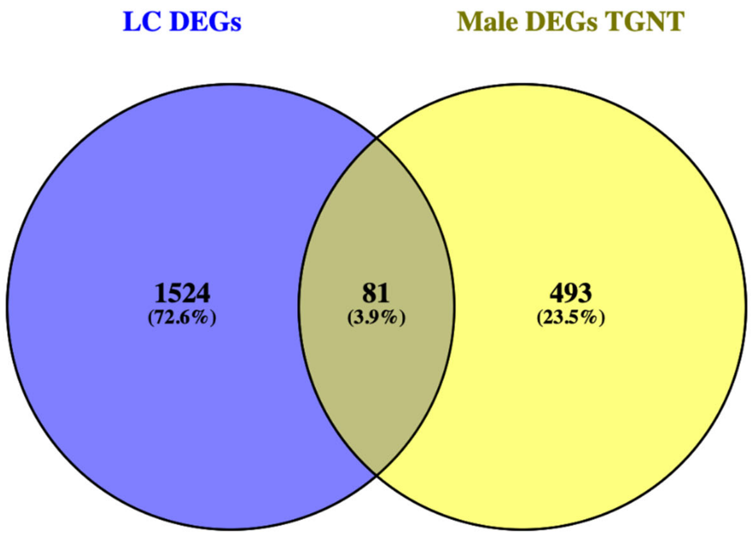

| LC DEGs | Male DEGs TGNT | 81 Shared genes | 63 upregulated out of 81 | 18 downregulated out of 81 |
|---|---|---|---|---|
| cnga1 | psen1 | slc35d3 | tbc1d10c | slc35d3 |
| linc00261 | app | hp | cenpf | hp |
| pitx3 | gfap | klhl1 | col7a1 | klhl1 |
| ensg00000272825 | gpnmb | six3 | zmynd10 | six3 |
| foxa1 | gpr88 | inmt | ptpn5 | inmt |
| slc6a3 | rgs9 | epn3 | kl | epn3 |
| ensg00000227176 | six3 | otx1 | sulf1 | otx1 |
| ensg00000272223 | adora2a | unc13c | clic6 | unc13c |
| ensg00000269986 | lhx8 | lmx1a | lif | lmx1a |
| ensg00000261502 | gpr6 | pax8 | kcnk2 | pax8 |
| hao1 | ecel1 | ret | h19 | ret |
| upb1 | sh3rf2 | corin | ngef | corin |
| foxa2 | akap12 | pappa | kcna5 | pappa |
| ensg00000287748 | syndig1l | drd2 | nmu | drd2 |
| fezf1-as1 | slc22a3 | fcgr2b | rbp4 | slc10a4 |

| | | | | |
|---|---|---|---|---|
| chrnb3 | drd1 | slc10a4 | irx6 | il21r |
| ensg00000225742 | foxp2 | tbc1d10c | gdf7 | ticam2 |
| cckar | slc4a5 | cenpf | gpr88 | gucy1a1 |
| ensg00000231062 | h19 | col7a1 | trh | |
| elovl3 | igf2 | zmynd10 | tac1 | |
| ensg00000287210 | folr1 | sstr3 | ecel1 | |
| ensg00000274248 | tmem27 | cnr1 | samd3 | |
| mt1h | col5a1 | kl | gulp1 | |
| duox2 | tmem72 | sulf1 | crabp1 | |
| pelp1-dt | penk | il21r | grm4 | |
| ensg00000287251 | fcgr2b | clic6 | slc4a11 | |
| ensg00000272909 | tnni3 | lif | ttr | |
| ensg00000290438 | ace | gucy1a1 | sostdc1 | |
| ensg00000290568 | mvd | kcnk2 | prlr | |
| hormad2 | scn4b | h19 | mybl2 | |
| lpo | kb15 | ngef | dclk3 | |
| kiaa1191p2 | cd22 | adra1b | meis2 | |
| aldh1a1 | f5 | kcna5 | evpl | |
| ensg00000260658 | drd2 | slc6a7 | drd1 | |
| ensg00000274902 | sbk2 | slc6a5 | vwa5b1 | |
| loc100128093 | kl | nmu | pitx2 | |
| ensg00000272735 | sulf1 | rbp4 | veph1 | |
| rspo2 | itgb6 | irx6 | htr1d | |
| ensg00000290059 | cdh3 | gdf7 | coch | |
| ntsr1 | lect1 | gpr88 | gsc2 | |
| rasef | cd74 | gal | penk | |
| atp2a3 | rgd1559696 | trh | mme | |
| pawrp2 | mfrp | sstr2 | klk7 | |
| lmod3 | slc5a7 | tac1 | capsl | |
| fezf1 | enpp2 | ecel1 | slc5a7 | |
| ensg00000228204 | itpripl1 | samd3 | slc18a3 | |
| fbxo40 | ppp1r1b | gulp1 | penk | |
| ensg00000288720 | trpv4 | crabp1 | hcrtr1 | |
| tff3 | hist1h2bo | grm4 | adcyap1 | |
| sdc1 | scgb1c1 | slc4a11 | tacr3 | |
| lemd1 | samd3 | ttr | npy2r | |
| st8sia6-as1 | col3a1 | galr1 | slc6a5 | |
| slc24a5 | igfbpl1 | sostdc1 | tac3 | |
| agtr1 | inmt | prlr | slc17a6 | |

| | | | |
|---|---|---|---|
| ensg00000273123 | krt71 | tacr3 | grm4 |
| linc01998 | tac1 | mybl2 | gal |
| snca-as1 | kcne2 | dclk3 | p2ry1 |
| arx | abca4 | npy2r | slc6a7 |
| ensg00000236948 | zic1 | p2ry1 | galr1 |
| linc02251 | calcr | meis2 | adra1d |
| ensg00000279689 | adipoq | hcrtr1 | cnr1 |
| ensg00000235450 | col1a2 | evpl | sstr3 |
| ensg00000233682 | lhx1 | tac3 | adra1b |
| dsel-as1 | naglt1 | drd1 | htr2c |
| slc18a1 | ret | slc17a6 | cartpt |
| daz2 | pappa | vwa5b1 | |
| slc35d3 | kcnj13 | pitx2 | |
| ensg00000272555 | h1foo | veph1 | |
| cyp27c1 | jph2 | adra1d | |
| ensg00000231167 | slco1a5 | htr1d | |
| nr2e1 | pdzd2 | coch | |
| ensg00000271623 | gapdh-ps1 | adcyap1 | |
| part1 | sult1c3 | gsc2 | |
| ensg00000271218 | htr1d | penk | |
| ensg00000239040 | myl6 | mme | |
| loc102723324 | leap2 | klk7 | |
| ensg00000288046 | ogn | capsl | |
| ensg00000287689 | nexn | slc5a7 | |
| ccdc116 | mapk4 | cartpt | |
| ensg00000287663 | tbc1d10c | slc18a3 | |
| ensg00000240499 | rmrp | ticam2 | |
| ensg00000272862 | ttr | | |
| rxfp3 | slc13a4 | | |
| ensg00000266441 | lsp1 | | |
| trpc6 | tex15 | | |
| ensg00000287369 | cldn2 | | |
| map3k20-as1 | rn28s | | |
| acmsd | rasd2 | | |
| ensg00000232457 | slc18a3 | | |
| ensg00000282304 | vwce | | |
| rnu1-3 | otx2 | | |
| ensg00000286201 | trh | | |
| il1rapl2 | pla2g5 | | |

| | |
|---|---|
| ensg00000247324 | asic4 |
| sp7 | cd68 |
| lgr4-as1 | slc35d3 |
| daz4 | neu2 |
| syngr4 | rgd1561102 |
| rnu1-1 | loc287004 |
| htr2b | ccnb1ip1 |
| sim2 | prlr |
| ensg00000285649 | trat1 |
| tmc5 | irx3 |
| rnu1-27p | arl11 |
| rnu1-28p | capg |
| chrna6 | stra6 |
| cyp4f12 | lepr |
| hpgd | hnf4a |
| ensg00000258910 | atp6ap1l |
| alg1l13p | fgf1 |
| rnu1-2 | cmtm1 |
| f2rl2 | clic6 |
| rnu1-4 | runx1 |
| smpx | pcp4l1 |
| cadps2 | col5a2 |
| dand5 | col7a1 |
| angptl5 | ccl9 |
| ensg00000279583 | cartpt |
| ensg00000286104 | isl1 |
| ensg00000239572 | rgd1561795 |
| ensg00000255046 | gbx2 |
| npnt | cd200r1 |
| ensg00000260661 | sostdc1 |
| c1orf127 | dazl |
| loc124904613 | slc22a8 |
| st8sia6 | meis2 |
| ensg00000188681 | itgb1bp2 |
| slc22a18as | mpz |
| ensg00000278231 | tssk3 |
| ensg00000256673 | hp |
| ensg00000279296 | pcp4 |
| mep1a | loc691083 |

| | |
|---|---|
| loc100420423 | tcf21 |
| hp | kcnq1 |
| arg1 | glycam1 |
| ensg00000232713 | pon3 |
| ensg00000251680 | rbm47 |
| en1 | gpr52 |
| deup1 | fam160a1 |
| tnmd | pde7b |
| rnvu1-18 | tbx6 |
| ctage4 | ltbp2 |
| ensg00000280310 | tmem184a |
| asb16 | frem1 |
| gabra4 | il13ra2 |
| ensg00000267751 | mpzl2 |
| ccdc38 | ptprq |
| ensg00000272049 | itk |
| zar 1.00 | clul1 |
| cyp2s1 | melk |
| ensg00000230929 | ccl3 |
| ensg00000261357 | tnnt2 |
| or7e28p | rgd1561161 |
| ensg00000287801 | lgals5 |
| taf12-dt | spetex-2a |
| htr3b | fetub |
| loc124904613 | rarb |
| ensg00000273399 | aqp1 |
| mmp8 | klk4 |
| linc01291 | trpv5 |
| cnih3-as2 | pon1 |
| rdh12 | ncf2 |
| ensg00000253620 | kremen1 |
| loc102724900 | enkur |
| mlph | hes3 |
| ensg00000289701 | pdp1 |
| kcnj6 | igfbp2 |
| mchr2 | slc4a2 |
| srpx2 | mdfic |
| ensg00000257527 | lhx5 |
| ensg00000259146 | steap1 |

| | |
|---|---|
| tafa3 | pde10a |
| ensg00000254481 | scube3 |
| ensg00000227755 | il20ra |
| ensg00000272941 | iqgap3 |
| klhl1 | gbx1 |
| loc107985177 | camp |
| six3 | slc16a12 |
| cyp4f3 | vav1 |
| ensg00000289171 | s100g |
| matn3 | mxd3 |
| hspa8p4 | dmkn |
| siah3 | cdr2 |
| vgll3 | kcnv2 |
| inmt | lmx1a |
| tpbg | kcnh4 |
| cpxm1 | brs3 |
| ensg00000225473 | gucy2f |
| c1orf94 | naa11 |
| gfra1 | pih1d3 |
| ensg00000289149 | mcoln3 |
| stpg2 | crabp1 |
| linc00924 | loc100363193 |
| ntn1 | mc3r |
| ensg00000286646 | cdx4 |
| trpm1 | gucy1a3 |
| linc00486 | loc654482 |
| vav3 | cdk1 |
| birc7 | zscan10 |
| ccdc178 | veph1 |
| ensg00000279587 | ngef |
| ensg00000211829 | adra2b |
| tent5b | fbxo39 |
| gprc5a | mcoln2 |
| chrm5 | aldh1a3 |
| prmt8 | loc102548399 |
| catsperg | slc16a8 |
| ensg00000277867 | tnn |
| twist1 | gnal |
| ensg00000256757 | trpm3 |

| | |
|---|---|
| ensg00000279289 | lag3 |
| gpx8 | capsl |
| ensg00000287828 | hcls1 |
| ensg00000276337 | cdhr1 |
| arhgef4-as1 | avpr2 |
| ensg00000231086 | fam187a |
| ppdpfl | rgd1563692 |
| prok2 | slc2a12 |
| ensg00000228528 | pcp2 |
| ensg00000267422 | lrrk2 |
| epn3 | bhmt2 |
| col12a1 | tns4 |
| ensg00000272779 | igfals |
| sstr1 | pitx2 |
| adm2 | ido1 |
| atxn8os | insrr |
| loc105378355 | coch |
| otx1 | tmem196 |
| galntl6 | nr5a1 |
| chrna4 | pde1b |
| ensg00000287139 | dclk3 |
| linc00484 | rgd1308065 |
| tex22 | col18a1 |
| robo2 | adamtsl4 |
| thcat155 | has1 |
| areg | fhod3 |
| ensg00000255200 | ugt1a6 |
| ensg00000260331 | cdkn1c |
| lilrb5 | bcmo1 |
| mir4500hg | runx3 |
| ensg00000273654 | lilrb4 |
| ensg00000291111 | irx6 |
| unc13c | spock3 |
| znf99 | rbp4 |
| eno3 | krt42 |
| firre | retn |
| cr1 | bcl2l11 |
| mtcp1 | c1qtnf3 |
| ehbp1-as1 | prkar2b |

| | |
|---|---|
| kcnd3 | atp8b1 |
| rerg | acot12 |
| linc00940 | rasgrp2 |
| mcc | arpp21 |
| ensg00000225331 | fam46a |
| ensg00000279578 | atp4a |
| clstn2 | prdm1 |
| ano1 | mgp |
| ensg00000267714 | kif23 |
| psors1c1 | reck |
| linc00706 | rt1-da |
| linc00862 | hs3st3a1 |
| asb4 | lrrc18 |
| pygl | mybl2 |
| ybx2 | ces1d |
| ensg00000270157 | foxj1 |
| cd8b2 | xcl1 |
| ensg00000286282 | slpil2 |
| azgp1 | arl4d |
| ensg00000224842 | krt85 |
| tnfrsf10c | pou4f3 |
| ensg00000278962 | hist1h2aa |
| ensg00000231170 | crb3 |
| linc01019 | hist1h2ak |
| lrrc14b | aldoart2 |
| gsdmc | vom1r16 |
| lrrc55 | msx1 |
| gpr26 | ttc26 |
| aqp9 | serpinc1 |
| misp | hist1h1b |
| linc01819 | rnf152 |
| tmem221 | arhgap30 |
| linc02473 | plek |
| ensg00000286913 | prkch |
| nyap2 | mdk |
| ensg00000280850 | sgcg |
| klhl13 | rgd1560608 |
| cyp39a1 | slc10a4 |
| padi1 | ddc8 |

| | |
|---|---|
| ensg00000227088 | c3ar1 |
| loc101927609 | a2m |
| ensg00000261596 | efcab1 |
| ensg00000272054 | ccdc40 |
| linc01010 | mak |
| kank4 | nmu |
| ensg00000285601 | tmem170a |
| ankrd29 | loc100361092 |
| afap1-as1 | klk7 |
| itpr1 | nkx2-1 |
| lmx1a | zic2 |
| pax8 | cxcl13 |
| spata9 | loxl4 |
| linc00906 | rrm2 |
| apob | pabpc2 |
| ret | rgd1303271 |
| l3mbtl4 | slc14a2 |
| ncapg | slc5a12 |
| ensg00000245248 | reg3a |
| ensg00000260469 | top2a |
| linc00624 | rt1-ba |
| linc00574 | bmp2 |
| sfrp5 | elovl7 |
| dapl1 | unc13c |
| ensg00000273387 | uncx |
| nat1 | itga5 |
| znf385d | foxb1 |
| corin | klra5 |
| lcn2 | ces2 |
| pappa | fam71f1 |
| ensg00000285588 | pnpla1 |
| fbln7 | pdzk1ip1 |
| gbe1 | nepn |
| linc03016 | spink5 |
| ensg00000226268 | rgd1561870 |
| h3c6 | loc689713 |
| ensg00000271452 | slc16a2 |
| lmx1b | tshr |
| ensg00000234911 | prkag3 |

| | |
|---|---|
| ptpn3 | slc7a12 |
| tdrd10 | sct |
| ensg00000248719 | rgd1561958 |
| sox6 | fgf20 |
| mamdc2 | klk1 |
| ensg00000226578 | adcy5 |
| hhip | cpne5 |
| ensg00000268049 | slc12a7 |
| cd209 | st6galnac2 |
| cd300e | akp3 |
| cldn1 | tc2n |
| insyn1 | loxl2 |
| cntn4 | kcna5 |
| slc39a12 | cmtm8 |
| cplx2 | fam109b |
| fut9 | cyp11b3 |
| omd | fyb |
| ensg00000228541 | chad |
| erbb4 | cenpf |
| ensg00000261220 | car13 |
| ensg00000229107 | fbp1 |
| pkib | vgll2 |
| lrrc39 | ldhal6b |
| drd2 | htr6 |
| rcan2 | kiss1 |
| pkd2l2 | ms4a14 |
| pou3f2 | krt18 |
| dnmbp-as1 | prima1 |
| rfesd | gsc2 |
| aard | aox3 |
| mmp9 | mmp2 |
| ensg00000272910 | slc4a11 |
| cpb1 | krt75 |
| klf5 | rgd1562658 |
| ensg00000215302 | loc100910620 |
| tox3 | fmo2 |
| efcab5 | mael |
| ensg00000278416 | serpine1 |
| linc02177 | olr200 |

| | |
|---|---|
| znf728 | txk |
| ensg00000230359 | gck |
| pgm5p2 | fam222a |
| znf157 | clec4e |
| steap1b | cgnl1 |
| trdn | f13a1 |
| ensg00000235421 | six1 |
| ppp1r1c | smim22 |
| ensg00000272345 | lif |
| lgi2 | hist2h4 |
| ensg00000269940 | slc5a5 |
| slc2a13 | bhmt |
| asb5 | itpr3 |
| adrb1 | efhb |
| ensg00000210100 | ccl4 |
| garin1b | art3 |
| ensg00000290034 | loc171161 |
| fer1l6-as2 | rgd1305627 |
| ensg00000272630 | nags |
| tcf12-dt | myo1f |
| gpr161 | dlx5 |
| pgr | wdr16 |
| acp7 | grm4 |
| znf503-as2 | corin |
| plcb1 | phactr2 |
| fcgr2b | kcnk2 |
| znf503 | kcnn4 |
| mt1g | gulp1 |
| c10orf90 | wdr63 |
| ensg00000286558 | glb1l |
| spanxa2-ot1 | pde6b |
| cedora | rab38 |
| rnu6-415p | gpr153 |
| pcdhb7 | ccdc166 |
| inpp4b | gypc |
| ddr2 | klhl1 |
| st6galnac5 | msc |
| jtb-dt | ifltd1 |
| pbx1 | baiap2l1 |

| | |
|---|---|
| olfml2b | cd72 |
| map3k20 | mme |
| ensg00000262097 | actn3 |
| mitf | sfrp1 |
| loc100505715 | car14 |
| aunip | rt1-ce3 |
| ctxn3 | slc31a1 |
| ensg00000232504 | rab11fip1 |
| linc02884 | npr3 |
| pcdha5 | clec2d |
| dab1 | tnxa-ps1 |
| fap | slc16a3 |
| dkk 1.00 | best3 |
| ensg00000277304 | spag8 |
| myom1 | iqcg |
| ank1 | hist2h3c2 |
| ensg00000205106 | ube2l6 |
| b3galt1 | mcm10 |
| cdh19 | stk32b |
| tmt1b | entpd3 |
| linc01554 | tpm2 |
| pid1 | dnali1 |
| ube2fp3 | egfr |
| shisa3 | tinagl1 |
| styk1 | zic3 |
| rlbp1 | cobl |
| rgs6 | ect2 |
| cdcp1 | exo1 |
| ensg00000210107 | stat4 |
| mpl | lysmd3 |
| loc124901457 | tgm3 |
| lmo3 | fscn2 |
| epha6 | ccdc60 |
| kcnn3 | rt1-db1 |
| lyve1 | chrdl2 |
| slc16a9 | hnf1b |
| plcxd2 | rt1-ce15 |
| tmem132b | clec5a |
| loc101929130 | il18bp |

| | |
|---|---|
| adamts18 | ajuba |
| marco | rgd1559714 |
| vxn | sh2b2 |
| hycc1 | sytl3 |
| rgs8 | dnai2 |
| eda2r | bst1 |
| mcm2 | il2rg |
| mrc1 | pou1f1 |
| slc10a4 | gdf7 |
| chrna2 | pld5 |
| mpped1 | crb1 |
| diaph3 | ccdc114 |
| fabp7 | cdca7 |
| tmprss5 | timd2 |
| nek10 | unc13d |
| ptgis | loc689927 |
| ensg00000249790 | crygs |
| cyp4x1 | rgd1565655 |
| nek5 | rrh |
| sytl5 | spp1 |
| grik1 | def6 |
| ensg00000272501 | gja6 |
| alb | vom1r88 |
| enpep | lyzl6 |
| cacna2d1 | cmah |
| ensg00000288538 | txndc2 |
| shc3 | mirlet7d |
| ednra | cb707485 |
| tmem132e | olr1667 |
| chrna7 | testin |
| ttty14 | cyp2c24 |
| ensg00000261542 | fgg |
| gckr | lrrc52 |
| ccna1 | odf3l2 |
| ensg00000277701 | dmrtc1c1 |
| ensg00000286592 | krt86 |
| ensg00000210144 | cd69 |
| mmd2 | agtr2 |
| gpr83 | cryge |

| | |
|---|---|
| n4bp2 | dcm5 |
| flrt2-as1 | oas1h |
| pcdha3 | ugt1a9 |
| linc01238 | tcp10b |
| blacat1 | cldn22 |
| loc101928416 | olr1653 |
| brinp3 | sell |
| ly6h | adh7 |
| txndc12 | mirlet7c-2 |
| ensg00000290818 | mir3572 |
| ptger1 | olr1869 |
| vangl1 | efhc1 |
| tbc1d10c | pbx3 |
| ensg00000263731 | gfi1b |
| cacng4 | gykl1 |
| b3gat2 | adamts19 |
| ensg00000261770 | ropn1l |
| oaz3 | lrr1 |
| mapk13 | otx1 |
| akr1c1 | hepacam2 |
| ncald | krt34 |
| ensg00000273247 | olr387 |
| dnase1l2 | epn3 |
| lzts1 | cyp11a1 |
| ensg00000287574 | slamf6 |
| ass1 | crygf |
| l3mbtl1 | krt1 |
| crlf1 | foxc2 |
| cytl1 | dab2 |
| grem2 | mrgprf |
| ensg00000215908 | troap |
| gsto2 | evpl |
| sox1 | prelp |
| bdnf | igsf9 |
| ensg00000227388 | mir421 |
| ensg00000248126 | adcy7 |
| cacng3 | vwa5b1 |
| isyna1 | pof1b |
| amh | rdh7 |

| | |
|---|---|
| rbm24 | fam83a |
| fam184b | rgd1561778 |
| linc02611 | tead2 |
| znf876p | cox8b |
| cenpf | llgl2 |
| baat | dpp4 |
| cfap210 | scara5 |
| cfap44 | kcna10 |
| ensg00000287737 | rgd1563104 |
| cd247 | trim40 |
| phactr1 | ugt1a5 |
| mir7-3hg | apobec1 |
| linc01115 | krt16 |
| ankrd30b | rin3 |
| letm2 | s100a5 |
| linc02930 | kctd8 |
| gpc1-as1 | zim1 |
| linc02983 | sp8 |
| mrpl23-as1 | upk1b |
| fam246c | nlrp4 |
| fer1l4 | iqub |
| gxylt1p5 | vom2r60 |
| itgb2-as1 | lipogenin |
| lpin3 | zmynd10 |
| ensg00000267605 | pax8 |
| ensg00000288802 | gpx2 |
| ensg00000259347 | fgf19 |
| ensg00000279113 | epha1 |
| lratd1 | il22ra1 |
| ensg00000226530 | kcnmb3 |
| slc7a14 | magea4 |
| nxph2 | olr176 |
| adamts7p1 | penk |
| loc105373299 | hcrtr1 |
| loc124907392 | adcyap1 |
| nt5dc3 | tacr3 |
| sytl1 | npy2r |
| linc03126 | slc6a5 |
| cfap43 | sstr2 |

| | |
|---|---|
| aarsd1 | tac3 |
| nos2 | slc17a6 |
| ptk7 | grm4 |
| ypel1 | gal |
| kcnh5 | p2ry1 |
| lancl3 | slc6a7 |
| ensg00000280157 | galr1 |
| rn7sl268p | adra1d |
| ensg00000269694 | cnr1 |
| ecm1 | sstr3 |
| il17re | adra1b |
| ensg00000272374 | htr2c |
| hmx1 | il21r |
| cfap74 | ticam2 |
| ccl3l1 | gucy1a1 |
| mybph | |
| hcg27 | |
| radil | |
| ensg00000291054 | |
| spag5 | |
| ensg00000225964 | |
| htr1b | |
| galnt16 | |
| peg3 | |
| mamld1 | |
| egr4 | |
| magee2 | |
| myo5b | |
| scn9a | |
| ensg00000272148 | |
| znf215 | |
| kctd16 | |
| ensg00000279259 | |
| trim74 | |
| loc124905142 | |
| grik2 | |
| fcmr | |
| uqcrb-as1 | |
| brwd1-as2 | |

| |
|---|
| prrg3 |
| linc00313 |
| gpr150 |
| mtmr11 |
| dlec1 |
| kif11 |
| sst |
| ensg00000279894 |
| ensg00000230587 |
| arhgef16 |
| mtmr9lp |
| kncn |
| ensg00000276445 |
| tmem74b |
| tspan32 |
| mir646hg |
| ensg00000272512 |
| wnt4 |
| rskr |
| linc00319 |
| lrp8-dt |
| maoa |
| rpl7ap43 |
| loc284798 |
| fam106a |
| plac8l1 |
| uchl3 |
| fgfr1 |
| wasir2 |
| mzf1-as1 |
| gjd2 |
| anos1 |
| col7a1 |
| zmynd10 |
| sstr3 |
| ensg00000242242 |
| cfap99 |
| tmie |
| loc100505716 |

| |
|---|
| pnma6f |
| nynrin |
| psma2 |
| prdm12 |
| cfap251 |
| ensg00000280057 |
| magel2 |
| ensg00000279138 |
| tcerg1l |
| sox1-ot |
| ensg00000288764 |
| linc02082 |
| prkar1b-as1 |
| cnr1 |
| zap70 |
| dnai1 |
| ensg00000289334 |
| linc01144 |
| sh2d2a |
| akr1c2 |
| kcnc4 |
| palm3 |
| lekr1 |
| gemin7-as1 |
| col1a1 |
| vars2 |
| tmsb15b |
| erich6-as1 |
| wnt5b |
| ensg00000231443 |
| fam246a |
| ensg00000247373 |
| mgat4c |
| ensg00000134297 |
| gng13 |
| cenpw |
| nnat |
| ptpn5 |
| trbv20or9-2 |

| |
|---|
| plekhg4b |
| rnf207-as1 |
| rnf182 |
| tent5a |
| galr2 |
| itga2b |
| trip13 |
| traf5 |
| kl |
| unc5d |
| ensg00000258520 |
| ceacam19 |
| ensg00000214783 |
| tmem72-as1 |
| hap1 |
| trank1 |
| cited2 |
| tmc4 |
| morn3 |
| vat1l |
| ensg00000271576 |
| ensg00000175658 |
| ensg00000255310 |
| tec |
| sulf1 |
| lrrc46 |
| msantd1 |
| ankrd36bp2 |
| ovgp1 |
| dpy19l2p4 |
| agap6 |
| ensg00000279108 |
| trib3 |
| calb1 |
| lrrc9 |
| klhl41 |
| slc30a3 |
| amotl1 |
| proc |

| |
|---|
| c1orf220 |
| kif24 |
| dpy19l2p1 |
| slc35f4 |
| ensg00000269918 |
| ensg00000250986 |
| ensg00000284633 |
| sertm1 |
| chgb |
| rem2 |
| ensg00000285804 |
| slc6a17 |
| myh7 |
| linc02691 |
| fam153a |
| ensg00000275180 |
| ensg00000235994 |
| znf534 |
| zglp1 |
| tspan18 |
| hsd17b7p2 |
| ensg00000289731 |
| arc |
| ensg00000286457 |
| wdr97 |
| loc100505915 |
| ensg00000234636 |
| spata18 |
| cenpvl2 |
| il21r |
| c20orf204 |
| ca10 |
| mns1 |
| ensg00000232721 |
| acot4 |
| cenpvl1 |
| tunar |
| als2cl |
| glb1l3 |

| |
|---|
| ensg00000289440 |
| plk5 |
| pou4f1 |
| ensg00000253361 |
| trpc7 |
| ensg00000267543 |
| ensg00000272953 |
| ensg00000277767 |
| linc02593 |
| dut-as1 |
| nr1i2 |
| stard4-as1 |
| eola1-dt |
| fam238c |
| ano3 |
| pwwp3b |
| hunk |
| loc388242 |
| loc105377209 |
| gng2 |
| ensg00000286299 |
| ensg00000263126 |
| dlk1 |
| il12rb2 |
| clic6 |
| sctr-as1 |
| ensg00000184303 |
| sall3 |
| ensg00000273674 |
| klrc2 |
| ildr2 |
| nkx2-8 |
| slf1 |
| icam5 |
| ensg00000259321 |
| ensg00000239556 |
| mir381hg |
| ensg00000257935 |
| lif |

| |
|---|
| prss56 |
| ensg00000223361 |
| pmfbp1 |
| mei1 |
| tmem132e-dt |
| linc03095 |
| ensg00000291067 |
| cngb1 |
| gucy1a1 |
| lum |
| syce1l |
| slc4a9 |
| ensg00000264456 |
| ensg00000274718 |
| trim17 |
| spag17 |
| ensg00000290683 |
| erich5 |
| linc01497 |
| pah |
| cpne9 |
| ensg00000271966 |
| plppr5 |
| linc01902 |
| ensg00000287625 |
| loc730101 |
| tspan10 |
| kiss1r |
| kcnk2 |
| dio3 |
| ensg00000267319 |
| ensg00000264666 |
| h19 |
| ensg00000256008 |
| c2orf66 |
| bean1 |
| bmper |
| mei4 |
| ensg00000278107 |

| |
|---|
| ensg00000288548 |
| ensg00000285955 |
| rps6ka2-it1 |
| ensg00000259683 |
| skor1 |
| kif23-as1 |
| ngef |
| meis3 |
| epb41l4b |
| espnl |
| ensg00000274330 |
| abracl |
| prmt5-dt |
| fgf14 |
| linc00923 |
| nhlh1 |
| scml4 |
| wnt11 |
| sfn |
| chp2 |
| nebl-as1 |
| calhm1 |
| kcnh3 |
| camk2a |
| ensg00000227953 |
| fndc3b |
| slc27a2 |
| angptl7 |
| adgrg2 |
| gpr68 |
| linc00951 |
| hmcn1 |
| ensg00000272420 |
| ensg00000261159 |
| map3k19 |
| prrt4 |
| muc4 |
| ensg00000278727 |
| fgf17 |

| |
|---|
| lrrn4cl |
| kif4a |
| ensg00000205740 |
| jmjd7-pla2g4b |
| rtn4r |
| pkp2 |
| ensg00000285725 |
| kif18b |
| sez6 |
| paqr9-as1 |
| rbfox1 |
| ensg00000273855 |
| ensg00000270071 |
| loc102724354 |
| ensg00000206706 |
| ensg00000277511 |
| ramp3 |
| tmc2 |
| ensg00000278000 |
| ensg00000288815 |
| p3h3 |
| cfap47 |
| npy |
| sept5-gp1bb |
| dnaaf1 |
| tmsb15b |
| crip3 |
| fstl5 |
| cfl1p1 |
| lhx4 |
| rnd3 |
| cfap57 |
| fras1 |
| cenpvl3 |
| stac |
| linc02910 |
| akr7l |
| dhrs2 |
| adra1b |

| |
|---|
| ensg00000289183 |
| ensg00000267251 |
| rab3b |
| cntn3 |
| cygb |
| ensg00000288557 |
| klk10 |
| fam153cp |
| ensg00000265194 |
| loricrin |
| foxd4l3 |
| iqcn |
| ensg00000231720 |
| cyp4z1 |
| h2bc8 |
| cntnap5-dt |
| gnb3 |
| pdyn |
| ensg00000278917 |
| ensg00000273139 |
| gng4 |
| napsb |
| sox3 |
| linc03112 |
| lncog |
| grm2 |
| cryba2 |
| ensg00000289061 |
| il11ra |
| dcbld2 |
| rna5sp508 |
| zbbx |
| ensg00000229127 |
| ensg00000205959 |
| lingo4 |
| chrm4 |
| slc25a41 |
| loc400622 |
| dnah2 |

| |
|---|
| pcolce2 |
| kif25 |
| ensg00000260274 |
| grm7 |
| foxp1-dt |
| linc01711 |
| daw1 |
| ensg00000282390 |
| pcdhgc5 |
| rpl7p23 |
| igsf1 |
| doc2b |
| oprd1 |
| c11orf21 |
| exosc10-as1 |
| ankrd55 |
| prokr1 |
| calb2 |
| islr2 |
| htr1a |
| loc100131939 |
| slc17a6-dt |
| col4a6 |
| aox1 |
| rbp3 |
| adgrf3 |
| muc12-as1 |
| kcna5 |
| ensg00000258847 |
| ccdc154 |
| mab21l1 |
| drd5 |
| treml1 |
| slc6a7 |
| pyy2 |
| mctp1 |
| ensg00000273188 |
| cfap45 |
| ensg00000288830 |

| |
|---|
| ensg00000272853 |
| ccdc162p |
| loc102723670 |
| vgf |
| syn2 |
| pspc1p1 |
| slc6a5 |
| bhlha9 |
| crabp2 |
| odad1 |
| nxnl2 |
| fgfr4 |
| igfl4 |
| tle6 |
| spmip5 |
| sod2-ot1 |
| ptger3 |
| slc25a28-dt |
| ensg00000288934 |
| loc101927237 |
| ensg00000271797 |
| ensg00000260927 |
| htr1e |
| ensg00000280255 |
| ensg00000267219 |
| ensg00000265984 |
| ensg00000287763 |
| crybb2 |
| foxo6 |
| b3gat1-dt |
| kif25-as1 |
| rhoxf1 |
| hint2 |
| morn5 |
| ensg00000291166 |
| met |
| gchfr |
| hspd1p10 |
| ensg00000254921 |

| |
|---|
| flj38576 |
| nmu |
| amigo2 |
| ensg00000269054 |
| grm8 |
| syt6 |
| ensg00000272807 |
| ensg00000289983 |
| ensg00000253477 |
| lrrn4 |
| rbp4 |
| irx6 |
| scgn |
| kit |
| ensg00000224680 |
| shisal2b |
| ensg00000287252 |
| adam21 |
| slco5a1 |
| ensg00000260306 |
| gdf7 |
| gpr88 |
| efcab12 |
| ensg00000277159 |
| ensg00000290673 |
| pklr |
| ptprr |
| golt1a |
| ensg00000271653 |
| loc100505978 |
| ensg00000271392 |
| zcchc18 |
| fam95a |
| cited1 |
| ensg00000284705 |
| ensg00000271396 |
| chrm1 |
| kctd14 |
| il11 |

| |
|---|
| rit2 |
| hypk |
| ensg00000286360 |
| ensg00000233765 |
| ensg00000263011 |
| st8sia2 |
| bmp3 |
| ensg00000279852 |
| ensg00000267645 |
| glyatl2 |
| cdh18 |
| cfap100 |
| fbxl22 |
| gal |
| ensg00000227455 |
| pou4f2 |
| doc2a |
| rgmb |
| ensg00000283689 |
| spdye3 |
| kiz-as1 |
| ensg00000259414 |
| c22orf42 |
| ensg00000224222 |
| ensg00000288107 |
| alk |
| scarna17 |
| npas4 |
| mimt1 |
| cyb561 |
| ensg00000261386 |
| slc34a3 |
| mmp10 |
| ensg00000287900 |
| ensg00000237629 |
| ensg00000272769 |
| trh |
| cfap65 |
| sstr2 |

| |
|---|
| rbfox3 |
| tac1 |
| nova1-dt |
| edn3 |
| dynlt2b |
| acr |
| ensg00000257366 |
| tfap2a |
| tgm1 |
| vwa3a |
| ensg00000286063 |
| ensg00000261669 |
| gpr139 |
| cpa2 |
| ensg00000196796 |
| ecel1 |
| sphkap |
| samd3 |
| ensg00000272459 |
| samd5 |
| hrk |
| ensg00000260296 |
| wdr93 |
| gulp1 |
| crabp1 |
| grm4 |
| slc4a11 |
| ensg00000186831 |
| picart1 |
| nt5dc2 |
| ttr |
| tnnc2 |
| cfap157 |
| ube2q2p13 |
| ensg00000256196 |
| nos1 |
| loc613038 |
| cd36 |
| ensg00000286550 |

| |
|---|
| sorcs3 |
| galr1 |
| linc02033 |
| abcc6p1 |
| nts |
| pitx1-as1 |
| ensg00000230537 |
| ifi27 |
| ensg00000279196 |
| sostdc1 |
| ndst4 |
| linc00958 |
| ensg00000280179 |
| ensg00000213089 |
| myl7 |
| spag6 |
| ensg00000288818 |
| tincr |
| linc02348 |
| hpd |
| loc105376654 |
| sntg2 |
| ensg00000289305 |
| loc124906789 |
| rmst |
| prss3 |
| casp12 |
| neto1 |
| loc101927245 |
| prok1 |
| onecut2 |
| nrg3-as1 |
| ensg00000271789 |
| ensg00000282870 |
| nptx1 |
| samd11 |
| linc01933 |
| nhlrc4 |
| zmat4 |

| |
|---|
| kcns2 |
| sorcs1 |
| tekt1 |
| prame |
| ensg00000228852 |
| hmcn2 |
| kcnk9 |
| tmem215 |
| kcnj3 |
| ensg00000253227 |
| ensg00000204791 |
| ensg00000233117 |
| ensg00000266473 |
| prlr |
| ensg00000278966 |
| ism2 |
| ensg00000272267 |
| dach1 |
| mycnos |
| cdh9 |
| ensg00000288721 |
| arrdc5 |
| tacr3 |
| sp6 |
| linc01586 |
| dnah6 |
| ensg00000275202 |
| mybl2 |
| ensg00000230882 |
| cdh22 |
| ensg00000279673 |
| kcnv1 |
| kiaa2012 |
| crhr2 |
| ensg00000230804 |
| ensg00000279601 |
| ensg00000176268 |
| neurod2 |
| ensg00000251495 |

| |
|---|
| agbl1 |
| ensg00000207294 |
| slc5a4 |
| pcdh19 |
| hadhap1 |
| ensg00000265282 |
| foxd4l6 |
| nxph4 |
| nell1 |
| necab2 |
| slc35g5 |
| zbtb32 |
| ensg00000279900 |
| npw |
| linc02192 |
| baiap3 |
| ensg00000259869 |
| sco2 |
| col2a1 |
| sctr |
| ensg00000273113 |
| ankrd26p3 |
| cbln4 |
| ighm |
| casr |
| ensg00000270557 |
| tph2 |
| ensg00000285873 |
| ensg00000271857 |
| ensg00000263326 |
| rpl7ap39 |
| linc01305 |
| dclk3 |
| ensg00000275613 |
| ensg00000286044 |
| thsd7b |
| rtp1 |
| neurod1 |
| krtap5-9 |

| |
|---|
| npy2r |
| ror2 |
| ensg00000279057 |
| linc00642 |
| ensg00000272971 |
| ensg00000289225 |
| loc105373390 |
| ptprt |
| npffr2 |
| ensg00000260487 |
| cyp4f26p |
| ensg00000280053 |
| wnt2 |
| acta2-as1 |
| p2ry1 |
| cyp26a1 |
| ndst3 |
| loc124906712 |
| linc02182 |
| rn7sl417p |
| ensg00000291062 |
| gpc3 |
| ensg00000281100 |
| ensg00000226747 |
| loc728307 |
| wnt6 |
| otx2-as1 |
| linc00606 |
| ensg00000277156 |
| ensg00000272386 |
| oprm1 |
| meis2 |
| cacna1i |
| kcnq5 |
| ensg00000259940 |
| hcrtr1 |
| efhd2-as1 |
| ct45a1 |
| klk5 |

| |
|---|
| rn7sl608p |
| linc02249 |
| ensg00000290918 |
| ensg00000276863 |
| tex26 |
| ensg00000285865 |
| ensg00000260410 |
| linc01201 |
| pou2f1-dt |
| ct45a3 |
| evpl |
| ensg00000188828 |
| linc02574 |
| ensg00000272321 |
| tac3 |
| ensg00000260743 |
| ensg00000272103 |
| ptgfr |
| ensg00000278012 |
| rtp5 |
| tuba3d |
| ensg00000267108 |
| sertm2 |
| kcnk3 |
| drd1 |
| enc1 |
| ensg00000272942 |
| slc17a6 |
| ensg00000237422 |
| glra2 |
| ensg00000287925 |
| ensg00000228444 |
| vwa5b1 |
| ensg00000260011 |
| ensg00000275178 |
| ensg00000288770 |
| ensg00000259031 |
| ensg00000269889 |
| c20orf203 |

| |
|---|
| pth2r |
| linc00494 |
| ensg00000225398 |
| ensg00000279821 |
| c7orf57 |
| prr33 |
| cfap58 |
| ensg00000249937 |
| pitx2 |
| ensg00000267986 |
| ensg00000271105 |
| pnma5 |
| npr 3.00 |
| tmem275 |
| prss3p4 |
| lhx2 |
| veph1 |
| linc01572 |
| ensg00000288271 |
| ensg00000269954 |
| ensg00000287792 |
| tpsd1 |
| fibcd1 |
| ensg00000254528 |
| linc00269 |
| ensg00000267466 |
| sema5b |
| ensg00000250053 |
| loc101929552 |
| rdm1p2 |
| ensg00000288808 |
| nxf2 |
| ensg00000287720 |
| ensg00000227256 |
| gpr176-dt |
| neurod6 |
| chat |
| ensg00000223298 |
| loc101927702 |

| |
|---|
| mrgpre |
| susd2 |
| ensg00000228950 |
| mylk3 |
| ensg00000223914 |
| ptprt-dt |
| ensg00000287617 |
| tnf |
| tfap2a-as2 |
| diras3 |
| linc00528 |
| nusap1 |
| ensg00000275769 |
| lrrc71 |
| adra1d |
| ensg00000287751 |
| htr1d |
| slear |
| linc02289 |
| morc1 |
| ensg00000290427 |
| gda |
| ensg00000280073 |
| tacr1 |
| neto1-dt |
| fev |
| linc01470 |
| e2f2 |
| pou2f2-as2 |
| ensg00000235020 |
| coch |
| c2cd4a |
| prlhr |
| slc30a8 |
| ensg00000289123 |
| prtn3 |
| drgx |
| ensg00000287209 |
| atp2c2-as1 |

| |
|---|
| art5 |
| ensg00000286527 |
| ensg00000283511 |
| cilp |
| otof |
| linc00487 |
| ensg00000230333 |
| fgf10 |
| tmem233 |
| ensg00000288090 |
| tnnc1 |
| meis1-as2 |
| kcnk17 |
| slc22a1 |
| linc01798 |
| pdgfd |
| ngb |
| adcyap1 |
| lhx9 |
| rprm |
| ensg00000240793 |
| ensg00000288990 |
| loc100133077 |
| adad2 |
| dutp6 |
| cpa4 |
| ensg00000258647 |
| shox2 |
| shank2-as1 |
| ensg00000288858 |
| egfl6 |
| linc02343 |
| fam163a |
| or51e1 |
| nwd2 |
| scgb3a1 |
| ensg00000265413 |
| gsc2 |
| ebf2 |

| |
|---|
| myh6 |
| pwwp4 |
| gabre |
| ccno |
| ensg00000248794 |
| cdc42bpg |
| ensg00000272081 |
| adra2a |
| calcb |
| fgl1 |
| otp |
| penk |
| ensg00000265179 |
| ensg00000204584 |
| pter |
| l1td1 |
| chodl |
| rfpl3s |
| lpar3 |
| ensg00000124549 |
| mab21l2 |
| ensg00000259977 |
| ensg00000260891 |
| mme |
| cyp24a1 |
| c1ql4 |
| ensg00000180438 |
| rprml |
| ensg00000280604 |
| ensg00000237886 |
| ct45a9 |
| rps17p16 |
| ensg00000285535 |
| lrrc53 |
| ensg00000287064 |
| umodl1-as1 |
| klk7 |
| c6orf141 |
| adgrd2 |

| |
|---|
| syt10 |
| fgf14-it1 |
| linc02642 |
| cntfr-as1 |
| dgkk |
| dusp13b |
| dnah3 |
| ankrd45 |
| ezhip |
| irx4 |
| ensg00000276603 |
| kctd19 |
| capsl |
| skor2 |
| trpa1 |
| ensg00000249741 |
| ct45a5 |
| six2 |
| slc7a3 |
| ensg00000279845 |
| crh |
| ensg00000267506 |
| tmc3 |
| lrrc38 |
| ensg00000205424 |
| gfra4 |
| linc01659 |
| ensg00000257681 |
| chrna3 |
| loc401442 |
| vsx2 |
| alx4 |
| ensg00000291225 |
| c1ql3 |
| ensg00000285696 |
| loc124904403 |
| ensg00000275678 |
| evx2 |
| ensg00000287827 |

| |
|---|
| cibar2 |
| gabrq |
| loc101928499 |
| linc00290 |
| frem3 |
| ensg00000251535 |
| dydc2 |
| slc6a4 |
| kcnj4 |
| tlx3 |
| myb |
| sec16b |
| ensg00000261886 |
| qrfpr |
| ensg00000278986 |
| ttll10 |
| slc5a7 |
| ensg00000280285 |
| ensg00000234891 |
| arhgap36 |
| evx1 |
| mir149 |
| ensg00000169662 |
| cartpt |
| uts2 |
| ct45a6 |
| ct45a7 |
| ensg00000257509 |
| ensg00000274080 |
| ensg00000203335 |
| lpa |
| ptf1a |
| cga |
| ensg00000276923 |
| slc9c2 |
| upk3a |
| ct45a10 |
| ensg00000248528 |
| tfap2b |

| |
|---|
| ino80b |
| ensg00000241720 |
| ensg00000280089 |
| ensg00000286407 |
| ensg00000236858 |
| dnaaf3 |
| slc18a3 |
| tmem114 |
| defa3 |
| dbh |
| ensg00000265018 |
| slc6a2 |
| cfap73 |
| ensg00000255372 |
| ensg00000286812 |
| ensg00000270462 |
| ct45a8 |
| linc01229 |
| ensg00000239377 |
| adamts16-dt |
| cfap107 |
| linc00682 |
| ensg00000254270 |
| cdhr4 |
| alpk2 |
| cfap299 |
| ensg00000287922 |
| slc35d3 |

**Supplementary table 20 : Common genes between LC dataset (Ehrenberg et al) and TgF344-AD rat DEGs (FEMALES)**

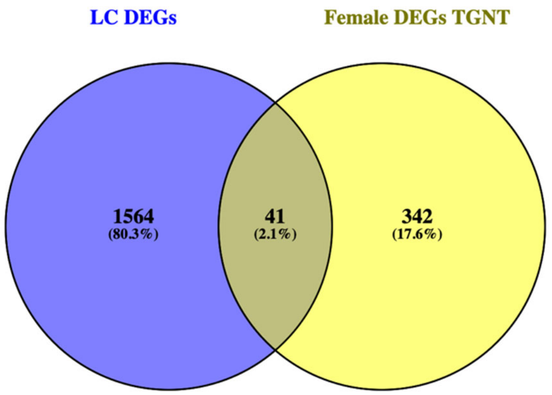

| LC DEGs | Female DEGs TGNT | Shared genes females | 21 upregulated out of 41 | 20 downregulated out of 41 |
|---|---|---|---|---|
| cnga1 | rgd1561102 | slc35d3 | mybl2 | slc35d3 |
| linc00261 | gapdh-ps1 | hp | dclk3 | hp |
| pitx3 | rmrp | klhl1 | meis2 | klhl1 |
| ensg00000272825 | ddc8 | epn3 | drd1 | epn3 |
| foxa1 | hist1h2aa | otx1 | veph1 | otx1 |
| slc6a3 | hp | unc13c | coch | unc13c |
| ensg00000227176 | akp3 | ret | penk | ret |
| ensg00000272223 | olr200 | corin | mme | corin |
| ensg00000269986 | tssk3 | fcgr2b | klk7 | slc10a4 |
| ensg00000261502 | krt42 | slc10a4 | capsl | tfrc |
| hao1 | tnn | cenpf | cartpt | gucy1a1 |
| upb1 | col5a1 | col7a1 | slc18a3 | myh6 |
| foxa2 | naa11 | ptpn5 | hap1 | myh7 |
| ensg00000287748 | avpr2 | sulf1 | psen2 | adra1b |
| fezf1-as1 | cd200r1 | lif | wnt4 | camk2a |
| chrnb3 | bcmo1 | kcnk2 | pdyn | cacng3 |
| ensg00000225742 | lif | ngef | nos1 | atp2a1 |
| cckar | loc100910620 | kcna5 | cnr1 | atp2a3 |
| ensg00000231062 | col7a1 | rbp4 | chrna7 | pdgfd |
| elovl3 | runx1 | irx6 | chrm5 | ntrk1 |
| ensg00000287210 | aldoart2 | gdf7 | slc18a1 | |

| | | |
|---|---|---|
| ensg00000274248 | prdm1 | trh |
| mt1h | loc100363193 | tac1 |
| duox2 | loc287004 | ecel1 |
| pelp1-dt | naglt1 | gulp1 |
| ensg00000287251 | ccl4 | crabp1 |
| ensg00000272909 | fam187a | grm4 |
| ensg00000290438 | pde6b | slc4a11 |
| ensg00000290568 | bcl2l11 | prlr |
| hormad2 | hist2h4 | mybl2 |
| lpo | hist1h2ak | dclk3 |
| kiaa1191p2 | serpine1 | meis2 |
| aldh1a1 | irx6 | drd1 |
| ensg00000260658 | loxl2 | veph1 |
| ensg00000274902 | has1 | coch |
| loc100128093 | fam109b | penk |
| ensg00000272735 | sim1 | mme |
| rspo2 | tnnt2 | klk7 |
| ensg00000290059 | ccl3 | capsl |
| ntsr1 | pabpc2 | cartpt |
| rasef | atp8b1 | slc18a3 |
| atp2a3 | hist2h3c2 | |
| pawrp2 | ltbp2 | |
| lmod3 | rgd1561958 | |
| fezf1 | ncf2 | |
| ensg00000228204 | cd72 | |
| fbxo40 | rgd1559714 | |
| ensg00000288720 | bst1 | |
| tff3 | slc16a3 | |
| sdc1 | kcnn4 | |
| lemd1 | tmem170a | |
| st8sia6-as1 | itga5 | |
| slc24a5 | lag3 | |
| agtr1 | bcl3 | |
| ensg00000273123 | spp1 | |
| linc01998 | col5a2 | |
| snca-as1 | def6 | |
| arx | il2rg | |
| ensg00000236948 | lsp1 | |
| linc02251 | f13a1 | |

| | |
|---|---|
| ensg00000279689 | cd68 |
| ensg00000235450 | rt1-ce3 |
| ensg00000233682 | psen1 |
| dsel-as1 | egfr |
| slc18a1 | cd22 |
| daz2 | rt1-ce15 |
| slc35d3 | ccdc166 |
| ensg00000272555 | app |
| cyp27c1 | rgd1309028 |
| ensg00000231167 | actn3 |
| nr2e1 | chad |
| ensg00000271623 | itpr3 |
| part1 | gfap |
| ensg00000271218 | cd74 |
| ensg00000239040 | arl11 |
| loc102723324 | ttc26 |
| ensg00000288046 | tpm2 |
| ensg00000287689 | rt1-da |
| ccdc116 | arhgap30 |
| ensg00000287663 | vav1 |
| ensg00000240499 | fyb |
| ensg00000272862 | rt1-ba |
| rxfp3 | clec5a |
| ensg00000266441 | ino80d |
| trpc6 | rab38 |
| ensg00000287369 | rt1-db1 |
| map3k20-as1 | gpnmb |
| acmsd | capg |
| ensg00000232457 | fcgr2b |
| ensg00000282304 | c3ar1 |
| rnu1-3 | rn28s |
| ensg00000286201 | stk32b |
| il1rapl2 | ube2l6 |
| ensg00000247324 | akap12 |
| sp7 | hcls1 |
| lgr4-as1 | mvd |
| daz4 | ret |
| syngr4 | timd2 |
| rnu1-1 | plek |

| | |
|---|---|
| htr2b | sh2b2 |
| sim2 | unc13d |
| ensg00000285649 | myo1f |
| tmc5 | arl4d |
| rnu1-27p | lrrk2 |
| rnu1-28p | tnxa-ps1 |
| chrna6 | lysmd3 |
| cyp4f12 | spock3 |
| hpgd | cobl |
| ensg00000258910 | prkar2b |
| alg1l13p | grm4 |
| rnu1-2 | pdp1 |
| f2rl2 | ptpn5 |
| rnu1-4 | mapk4 |
| smpx | fhod3 |
| cadps2 | cpne5 |
| dand5 | col18a1 |
| angptl5 | otx1 |
| ensg00000279583 | tinagl1 |
| ensg00000286104 | pdzd2 |
| ensg00000239572 | gnal |
| ensg00000255046 | pbx3 |
| npnt | unc13c |
| ensg00000260661 | rbp4 |
| c1orf127 | reck |
| loc124904613 | slc31a1 |
| st8sia6 | slc22a8 |
| ensg00000188681 | gpr153 |
| slc22a18as | prima1 |
| ensg00000278231 | slc16a2 |
| ensg00000256673 | dlx5 |
| ensg00000279296 | dab2 |
| mep1a | zic2 |
| loc100420423 | npr3 |
| hp | dnali1 |
| arg1 | slc4a11 |
| ensg00000232713 | mospd1 |
| ensg00000251680 | fam222a |
| en1 | prkch |

| | |
|---|---|
| deup1 | il18bp |
| tnmd | kcnk2 |
| rnvu1-18 | ngef |
| ctage4 | slc5a5 |
| ensg00000280310 | synpo2 |
| asb16 | mgp |
| gabra4 | ajuba |
| ensg00000267751 | dnai2 |
| ccdc38 | pde1b |
| ensg00000272049 | entpd3 |
| zar 1.00 | glb1l |
| cyp2s1 | adcy5 |
| ensg00000230929 | crb1 |
| ensg00000261357 | efhc1 |
| or7e28p | htr6 |
| ensg00000287801 | gulp1 |
| taf12-dt | pde7b |
| htr3b | wdr16 |
| loc124904613 | arpp21 |
| ensg00000273399 | gypc |
| mmp8 | pcp4 |
| linc01291 | car14 |
| cnih3-as2 | cgnl1 |
| rdh12 | elovl7 |
| ensg00000253620 | tmem196 |
| loc102724900 | ropn1l |
| mlph | rnf152 |
| ensg00000289701 | cdhr1 |
| kcnj6 | gng7 |
| mchr2 | slc12a7 |
| srpx2 | enkur |
| ensg00000257527 | sv2c |
| ensg00000259146 | art3 |
| tafa3 | cmtm8 |
| ensg00000254481 | fscn2 |
| ensg00000227755 | pnpla1 |
| ensg00000272941 | trpm3 |
| klhl1 | mrgprf |
| loc107985177 | sfrp1 |

| | |
|---|---|
| six3 | pde10a |
| cyp4f3 | rab11fip1 |
| ensg00000289171 | nags |
| matn3 | epn3 |
| hspa8p4 | wdr63 |
| siah3 | ccdc114 |
| vgll3 | pld5 |
| inmt | rrm2 |
| tpbg | tshr |
| cpxm1 | kcnh4 |
| ensg00000225473 | dclk3 |
| c1orf94 | cdca7 |
| gfra1 | zic3 |
| ensg00000289149 | nkx2-1 |
| stpg2 | phactr2 |
| linc00924 | meis2 |
| ntn1 | mme |
| ensg00000286646 | cdr2 |
| trpm1 | rgd1562658 |
| linc00486 | slc2a12 |
| vav3 | foxc2 |
| birc7 | fmo2 |
| ccdc178 | ugt1a6 |
| ensg00000279587 | ccdc40 |
| ensg00000211829 | a2m |
| tent5b | mmp2 |
| gprc5a | efcab1 |
| chrm5 | mlf1 |
| prmt8 | spag8 |
| catsperg | st6galnac2 |
| ensg00000277867 | tc2n |
| twist1 | iqcg |
| ensg00000256757 | sgcg |
| ensg00000279289 | rhod |
| gpx8 | cdkn1c |
| ensg00000287828 | crabp1 |
| ensg00000276337 | mdk |
| arhgef4-as1 | veph1 |
| ensg00000231086 | foxj1 |

| | |
|---|---|
| ppdpfl | gucy1a3 |
| prok2 | rgd1305627 |
| ensg00000228528 | bmp2 |
| ensg00000267422 | tex15 |
| epn3 | kif23 |
| col12a1 | gck |
| ensg00000272779 | kcna5 |
| sstr1 | aox3 |
| adm2 | kremen1 |
| atxn8os | ect2 |
| loc105378355 | stra6 |
| otx1 | adamts19 |
| galntl6 | baiap2l1 |
| chrna4 | loxl4 |
| ensg00000287139 | capsl |
| linc00484 | slc4a2 |
| tex22 | klhl1 |
| robo2 | msx1 |
| thcat155 | best3 |
| areg | adra2b |
| ensg00000255200 | mdfic |
| ensg00000260331 | rasd2 |
| lilrb5 | fam46a |
| mir4500hg | slamf6 |
| ensg00000273654 | rarb |
| ensg00000291111 | cenpf |
| unc13c | slc10a4 |
| znf99 | top2a |
| eno3 | krt1 |
| firre | iqgap3 |
| cr1 | mybl2 |
| mtcp1 | frem1 |
| ehbp1-as1 | rasgrp2 |
| kcnd3 | lhfpl1 |
| rerg | pcp4l1 |
| linc00940 | mcm10 |
| mcc | slc16a12 |
| ensg00000225331 | krt18 |
| ensg00000279578 | loc654482 |

| | |
|---|---|
| clstn2 | coch |
| ano1 | adamtsl4 |
| ensg00000267714 | hnf1b |
| psors1c1 | exo1 |
| linc00706 | krt75 |
| linc00862 | ifltd1 |
| asb4 | mael |
| pygl | rgd1560608 |
| ybx2 | txk |
| ensg00000270157 | lepr |
| cd8b2 | trh |
| ensg00000286282 | ccl9 |
| azgp1 | ogn |
| ensg00000224842 | otc |
| tnfrsf10c | bhmt |
| ensg00000278962 | loc100361092 |
| ensg00000231170 | asic4 |
| linc01019 | fam160a1 |
| lrrc14b | slc35d3 |
| gsdmc | crb3 |
| lrrc55 | cpn1 |
| gpr26 | cdk1 |
| aqp9 | atp6ap1l |
| misp | tgm3 |
| linc01819 | sell |
| tmem221 | hist1h1b |
| linc02473 | ccdc60 |
| ensg00000286913 | car13 |
| nyap2 | zic1 |
| ensg00000280850 | atp4a |
| klhl13 | pou1f1 |
| cyp39a1 | lrr1 |
| padi1 | chrdl2 |
| ensg00000227088 | gpr52 |
| loc101927609 | sytl3 |
| ensg00000261596 | efhb |
| ensg00000272054 | rgd1561161 |
| linc01010 | igfbp2 |
| kank4 | krt71 |

| | |
|---|---|
| ensg00000285601 | ppp1r1b |
| ankrd29 | mak |
| afap1-as1 | slc16a8 |
| itpr1 | corin |
| lmx1a | stat4 |
| pax8 | cartpt |
| spata9 | itk |
| linc00906 | rrh |
| apob | fgf1 |
| ret | sulf1 |
| l3mbtl4 | scube3 |
| ncapg | cyp11b3 |
| ensg00000245248 | klk7 |
| ensg00000260469 | scn4b |
| linc00624 | nexn |
| linc00574 | gdf7 |
| sfrp5 | ecel1 |
| dapl1 | clec2d |
| ensg00000273387 | cyp2c24 |
| nat1 | prlr |
| znf385d | cldn22 |
| corin | msc |
| lcn2 | lrrc18 |
| pappa | tac1 |
| ensg00000285588 | rbm47 |
| fbln7 | pih1d3 |
| gbe1 | penk |
| linc03016 | kcnv2 |
| ensg00000226268 | smim22 |
| h3c6 | drd1 |
| ensg00000271452 | igfbpl1 |
| lmx1b | clec4e |
| ensg00000234911 | slc13a4 |
| ptpn3 | mpzl2 |
| tdrd10 | mcoln3 |
| ensg00000248719 | neu2 |
| sox6 | krt85 |
| mamdc2 | crygf |
| ensg00000226578 | slc18a3 |

| | |
|---|---|
| hhip | cyp11a1 |
| ensg00000268049 | hepacam2 |
| cd209 | lrpprc |
| cd300e | dlg4 |
| cldn1 | pcdha6 |
| insyn1 | cox5b |
| cntn4 | map1b |
| slc39a12 | dock3 |
| cplx2 | spcs1 |
| fut9 | parp8 |
| omd | hdac9 |
| ensg00000228541 | actn2 |
| erbb4 | tmx3 |
| ensg00000261220 | atp5po |
| ensg00000229107 | dnm1l |
| pkib | syn1 |
| lrrc39 | efna5 |
| drd2 | tomm70 |
| rcan2 | atcay |
| pkd2l2 | fbxl16 |
| pou3f2 | aadat |
| dnmbp-as1 | col25a1 |
| rfesd | ica1l |
| aard | mrpl30 |
| mmp9 | ndufa11 |
| ensg00000272910 | ap3b2 |
| cpb1 | ndufs3 |
| klf5 | snx32 |
| ensg00000215302 | nnt |
| tox3 | mdh2 |
| efcab5 | stxbp1 |
| ensg00000278416 | slc1a2 |
| linc02177 | slit3 |
| znf728 | stx1b |
| ensg00000230359 | uqcrc2 |
| pgm5p2 | ablim2 |
| znf157 | ddx1 |
| steap1b | slc25a4 |
| trdn | slit1 |

| | |
|---|---|
| ensg00000235421 | aspm |
| ppp1r1c | mmp17 |
| ensg00000272345 | mrpl1 |
| lgi2 | slc9a7 |
| ensg00000269940 | atp1a3 |
| slc2a13 | cox6b1 |
| asb5 | dchs2 |
| adrb1 | mrpl15 |
| ensg00000210100 | dld |
| garin1b | pnpla3 |
| ensg00000290034 | atp6v1e1 |
| fer1l6-as2 | |
| ensg00000272630 | |
| tcf12-dt | |
| gpr161 | |
| pgr | |
| acp7 | |
| znf503-as2 | |
| plcb1 | |
| fcgr2b | |
| znf503 | |
| mt1g | |
| c10orf90 | |
| ensg00000286558 | |
| spanxa2-ot1 | |
| cedora | |
| rnu6-415p | |
| pcdhb7 | |
| inpp4b | |
| ddr2 | |
| st6galnac5 | |
| jtb-dt | |
| pbx1 | |
| olfml2b | |
| map3k20 | |
| ensg00000262097 | |
| mitf | |
| loc100505715 | |
| aunip | |

| |
|---|
| ctxn3 |
| ensg00000232504 |
| linc02884 |
| pcdha5 |
| dab1 |
| fap |
| dkk 1.00 |
| ensg00000277304 |
| myom1 |
| ank1 |
| ensg00000205106 |
| b3galt1 |
| cdh19 |
| tmt1b |
| linc01554 |
| pid1 |
| ube2fp3 |
| shisa3 |
| styk1 |
| rlbp1 |
| rgs6 |
| cdcp1 |
| ensg00000210107 |
| mpl |
| loc124901457 |
| lmo3 |
| epha6 |
| kcnn3 |
| lyve1 |
| slc16a9 |
| plcxd2 |
| tmem132b |
| loc101929130 |
| adamts18 |
| marco |
| vxn |
| hycc1 |
| rgs8 |
| eda2r |

| |
|---|
| mcm2 |
| mrc1 |
| slc10a4 |
| chrna2 |
| mpped1 |
| diaph3 |
| fabp7 |
| tmprss5 |
| nek10 |
| ptgis |
| ensg00000249790 |
| cyp4x1 |
| nek5 |
| sytl5 |
| grik1 |
| ensg00000272501 |
| alb |
| enpep |
| cacna2d1 |
| ensg00000288538 |
| shc3 |
| ednra |
| tmem132e |
| chrna7 |
| ttty14 |
| ensg00000261542 |
| gckr |
| ccna1 |
| ensg00000277701 |
| ensg00000286592 |
| ensg00000210144 |
| mmd2 |
| gpr83 |
| n4bp2 |
| flrt2-as1 |
| pcdha3 |
| linc01238 |
| blacat1 |
| loc101928416 |

| |
|---|
| brinp3 |
| ly6h |
| txndc12 |
| ensg00000290818 |
| ptger1 |
| vangl1 |
| tbc1d10c |
| ensg00000263731 |
| cacng4 |
| b3gat2 |
| ensg00000261770 |
| oaz3 |
| mapk13 |
| akr1c1 |
| ncald |
| ensg00000273247 |
| dnase1l2 |
| lzts1 |
| ensg00000287574 |
| ass1 |
| l3mbtl1 |
| crlf1 |
| cytl1 |
| grem2 |
| ensg00000215908 |
| gsto2 |
| sox1 |
| bdnf |
| ensg00000227388 |
| ensg00000248126 |
| cacng3 |
| isyna1 |
| amh |
| rbm24 |
| fam184b |
| linc02611 |
| znf876p |
| cenpf |
| baat |

| |
|---|
| cfap210 |
| cfap44 |
| ensg00000287737 |
| cd247 |
| phactr1 |
| mir7-3hg |
| linc01115 |
| ankrd30b |
| letm2 |
| linc02930 |
| gpc1-as1 |
| linc02983 |
| mrpl23-as1 |
| fam246c |
| fer1l4 |
| gxylt1p5 |
| itgb2-as1 |
| lpin3 |
| ensg00000267605 |
| ensg00000288802 |
| ensg00000259347 |
| ensg00000279113 |
| lratd1 |
| ensg00000226530 |
| slc7a14 |
| nxph2 |
| adamts7p1 |
| loc105373299 |
| loc124907392 |
| nt5dc3 |
| sytl1 |
| linc03126 |
| cfap43 |
| aarsd1 |
| nos2 |
| ptk7 |
| ypel1 |
| kcnh5 |
| lancl3 |

| |
|---|
| ensg00000280157 |
| rn7sl268p |
| ensg00000269694 |
| ecm1 |
| il17re |
| ensg00000272374 |
| hmx1 |
| cfap74 |
| ccl3l1 |
| mybph |
| hcg27 |
| radil |
| ensg00000291054 |
| spag5 |
| ensg00000225964 |
| htr1b |
| galnt16 |
| peg3 |
| mamld1 |
| egr4 |
| magee2 |
| myo5b |
| scn9a |
| ensg00000272148 |
| znf215 |
| kctd16 |
| ensg00000279259 |
| trim74 |
| loc124905142 |
| grik2 |
| fcmr |
| uqcrb-as1 |
| brwd1-as2 |
| prrg3 |
| linc00313 |
| gpr150 |
| mtmr11 |
| dlec1 |
| kif11 |

| |
|---|
| sst |
| ensg00000279894 |
| ensg00000230587 |
| arhgef16 |
| mtmr9lp |
| kncn |
| ensg00000276445 |
| tmem74b |
| tspan32 |
| mir646hg |
| ensg00000272512 |
| wnt4 |
| rskr |
| linc00319 |
| lrp8-dt |
| maoa |
| rpl7ap43 |
| loc284798 |
| fam106a |
| plac8l1 |
| uchl3 |
| fgfr1 |
| wasir2 |
| mzf1-as1 |
| gjd2 |
| anos1 |
| col7a1 |
| zmynd10 |
| sstr3 |
| ensg00000242242 |
| cfap99 |
| tmie |
| loc100505716 |
| pnma6f |
| nynrin |
| psma2 |
| prdm12 |
| cfap251 |
| ensg00000280057 |

| |
|---|
| magel2 |
| ensg00000279138 |
| tcerg1l |
| sox1-ot |
| ensg00000288764 |
| linc02082 |
| prkar1b-as1 |
| cnr1 |
| zap70 |
| dnai1 |
| ensg00000289334 |
| linc01144 |
| sh2d2a |
| akr1c2 |
| kcnc4 |
| palm3 |
| lekr1 |
| gemin7-as1 |
| col1a1 |
| vars2 |
| tmsb15b |
| erich6-as1 |
| wnt5b |
| ensg00000231443 |
| fam246a |
| ensg00000247373 |
| mgat4c |
| ensg00000134297 |
| gng13 |
| cenpw |
| nnat |
| ptpn5 |
| trbv20or9-2 |
| plekhg4b |
| rnf207-as1 |
| rnf182 |
| tent5a |
| galr2 |
| itga2b |

| |
|---|
| trip13 |
| traf5 |
| kl |
| unc5d |
| ensg00000258520 |
| ceacam19 |
| ensg00000214783 |
| tmem72-as1 |
| hap1 |
| trank1 |
| cited2 |
| tmc4 |
| morn3 |
| vat1l |
| ensg00000271576 |
| ensg00000175658 |
| ensg00000255310 |
| tec |
| sulf1 |
| lrrc46 |
| msantd1 |
| ankrd36bp2 |
| ovgp1 |
| dpy19l2p4 |
| agap6 |
| ensg00000279108 |
| trib3 |
| calb1 |
| lrrc9 |
| klhl41 |
| slc30a3 |
| amotl1 |
| proc |
| c1orf220 |
| kif24 |
| dpy19l2p1 |
| slc35f4 |
| ensg00000269918 |
| ensg00000250986 |

| |
|---|
| ensg00000284633 |
| sertm1 |
| chgb |
| rem2 |
| ensg00000285804 |
| slc6a17 |
| myh7 |
| linc02691 |
| fam153a |
| ensg00000275180 |
| ensg00000235994 |
| znf534 |
| zglp1 |
| tspan18 |
| hsd17b7p2 |
| ensg00000289731 |
| arc |
| ensg00000286457 |
| wdr97 |
| loc100505915 |
| ensg00000234636 |
| spata18 |
| cenpvl2 |
| il21r |
| c20orf204 |
| ca10 |
| mns1 |
| ensg00000232721 |
| acot4 |
| cenpvl1 |
| tunar |
| als2cl |
| glb1l3 |
| ensg00000289440 |
| plk5 |
| pou4f1 |
| ensg00000253361 |
| trpc7 |
| ensg00000267543 |

| |
|---|
| ensg00000272953 |
| ensg00000277767 |
| linc02593 |
| dut-as1 |
| nr1i2 |
| stard4-as1 |
| eola1-dt |
| fam238c |
| ano3 |
| pwwp3b |
| hunk |
| loc388242 |
| loc105377209 |
| gng2 |
| ensg00000286299 |
| ensg00000263126 |
| dlk1 |
| il12rb2 |
| clic6 |
| sctr-as1 |
| ensg00000184303 |
| sall3 |
| ensg00000273674 |
| klrc2 |
| ildr2 |
| nkx2-8 |
| slf1 |
| icam5 |
| ensg00000259321 |
| ensg00000239556 |
| mir381hg |
| ensg00000257935 |
| lif |
| prss56 |
| ensg00000223361 |
| pmfbp1 |
| mei1 |
| tmem132e-dt |
| linc03095 |

| |
|---|
| ensg00000291067 |
| cngb1 |
| gucy1a1 |
| lum |
| syce1l |
| slc4a9 |
| ensg00000264456 |
| ensg00000274718 |
| trim17 |
| spag17 |
| ensg00000290683 |
| erich5 |
| linc01497 |
| pah |
| cpne9 |
| ensg00000271966 |
| plppr5 |
| linc01902 |
| ensg00000287625 |
| loc730101 |
| tspan10 |
| kiss1r |
| kcnk2 |
| dio3 |
| ensg00000267319 |
| ensg00000264666 |
| h19 |
| ensg00000256008 |
| c2orf66 |
| bean1 |
| bmper |
| mei4 |
| ensg00000278107 |
| ensg00000288548 |
| ensg00000285955 |
| rps6ka2-it1 |
| ensg00000259683 |
| skor1 |
| kif23-as1 |

| |
|---|
| ngef |
| meis3 |
| epb41l4b |
| espnl |
| ensg00000274330 |
| abracl |
| prmt5-dt |
| fgf14 |
| linc00923 |
| nhlh1 |
| scml4 |
| wnt11 |
| sfn |
| chp2 |
| nebl-as1 |
| calhm1 |
| kcnh3 |
| camk2a |
| ensg00000227953 |
| fndc3b |
| slc27a2 |
| angptl7 |
| adgrg2 |
| gpr68 |
| linc00951 |
| hmcn1 |
| ensg00000272420 |
| ensg00000261159 |
| map3k19 |
| prrt4 |
| muc4 |
| ensg00000278727 |
| fgf17 |
| lrrn4cl |
| kif4a |
| ensg00000205740 |
| jmjd7-pla2g4b |
| rtn4r |
| pkp2 |

| |
|---|
| ensg00000285725 |
| kif18b |
| sez6 |
| paqr9-as1 |
| rbfox1 |
| ensg00000273855 |
| ensg00000270071 |
| loc102724354 |
| ensg00000206706 |
| ensg00000277511 |
| ramp3 |
| tmc2 |
| ensg00000278000 |
| ensg00000288815 |
| p3h3 |
| cfap47 |
| npy |
| sept5-gp1bb |
| dnaaf1 |
| tmsb15b |
| crip3 |
| fstl5 |
| cfl1p1 |
| lhx4 |
| rnd3 |
| cfap57 |
| fras1 |
| cenpvl3 |
| stac |
| linc02910 |
| akr7l |
| dhrs2 |
| adra1b |
| ensg00000289183 |
| ensg00000267251 |
| rab3b |
| cntn3 |
| cygb |
| ensg00000288557 |

| |
|---|
| klk10 |
| fam153cp |
| ensg00000265194 |
| loricrin |
| foxd4l3 |
| iqcn |
| ensg00000231720 |
| cyp4z1 |
| h2bc8 |
| cntnap5-dt |
| gnb3 |
| pdyn |
| ensg00000278917 |
| ensg00000273139 |
| gng4 |
| napsb |
| sox3 |
| linc03112 |
| lncog |
| grm2 |
| cryba2 |
| ensg00000289061 |
| il11ra |
| dcbld2 |
| rna5sp508 |
| zbbx |
| ensg00000229127 |
| ensg00000205959 |
| lingo4 |
| chrm4 |
| slc25a41 |
| loc400622 |
| dnah2 |
| pcolce2 |
| kif25 |
| ensg00000260274 |
| grm7 |
| foxp1-dt |
| linc01711 |

| |
|---|
| daw1 |
| ensg00000282390 |
| pcdhgc5 |
| rpl7p23 |
| igsf1 |
| doc2b |
| oprd1 |
| c11orf21 |
| exosc10-as1 |
| ankrd55 |
| prokr1 |
| calb2 |
| islr2 |
| htr1a |
| loc100131939 |
| slc17a6-dt |
| col4a6 |
| aox1 |
| rbp3 |
| adgrf3 |
| muc12-as1 |
| kcna5 |
| ensg00000258847 |
| ccdc154 |
| mab21l1 |
| drd5 |
| treml1 |
| slc6a7 |
| pyy2 |
| mctp1 |
| ensg00000273188 |
| cfap45 |
| ensg00000288830 |
| ensg00000272853 |
| ccdc162p |
| loc102723670 |
| vgf |
| syn2 |
| pspc1p1 |

| |
|---|
| slc6a5 |
| bhlha9 |
| crabp2 |
| odad1 |
| nxnl2 |
| fgfr4 |
| igfl4 |
| tle6 |
| spmip5 |
| sod2-ot1 |
| ptger3 |
| slc25a28-dt |
| ensg00000288934 |
| loc101927237 |
| ensg00000271797 |
| ensg00000260927 |
| htr1e |
| ensg00000280255 |
| ensg00000267219 |
| ensg00000265984 |
| ensg00000287763 |
| crybb2 |
| foxo6 |
| b3gat1-dt |
| kif25-as1 |
| rhoxf1 |
| hint2 |
| morn5 |
| ensg00000291166 |
| met |
| gchfr |
| hspd1p10 |
| ensg00000254921 |
| flj38576 |
| nmu |
| amigo2 |
| ensg00000269054 |
| grm8 |
| syt6 |

| |
|---|
| ensg00000272807 |
| ensg00000289983 |
| ensg00000253477 |
| lrrn4 |
| rbp4 |
| irx6 |
| scgn |
| kit |
| ensg00000224680 |
| shisal2b |
| ensg00000287252 |
| adam21 |
| slco5a1 |
| ensg00000260306 |
| gdf7 |
| gpr88 |
| efcab12 |
| ensg00000277159 |
| ensg00000290673 |
| pklr |
| ptprr |
| golt1a |
| ensg00000271653 |
| loc100505978 |
| ensg00000271392 |
| zcchc18 |
| fam95a |
| cited1 |
| ensg00000284705 |
| ensg00000271396 |
| chrm1 |
| kctd14 |
| il11 |
| rit2 |
| hypk |
| ensg00000286360 |
| ensg00000233765 |
| ensg00000263011 |
| st8sia2 |

| |
|---|
| bmp3 |
| ensg00000279852 |
| ensg00000267645 |
| glyatl2 |
| cdh18 |
| cfap100 |
| fbxl22 |
| gal |
| ensg00000227455 |
| pou4f2 |
| doc2a |
| rgmb |
| ensg00000283689 |
| spdye3 |
| kiz-as1 |
| ensg00000259414 |
| c22orf42 |
| ensg00000224222 |
| ensg00000288107 |
| alk |
| scarna17 |
| npas4 |
| mimt1 |
| cyb561 |
| ensg00000261386 |
| slc34a3 |
| mmp10 |
| ensg00000287900 |
| ensg00000237629 |
| ensg00000272769 |
| trh |
| cfap65 |
| sstr2 |
| rbfox3 |
| tac1 |
| nova1-dt |
| edn3 |
| dynlt2b |
| acr |

| |
|---|
| ensg00000257366 |
| tfap2a |
| tgm1 |
| vwa3a |
| ensg00000286063 |
| ensg00000261669 |
| gpr139 |
| cpa2 |
| ensg00000196796 |
| ecel1 |
| sphkap |
| samd3 |
| ensg00000272459 |
| samd5 |
| hrk |
| ensg00000260296 |
| wdr93 |
| gulp1 |
| crabp1 |
| grm4 |
| slc4a11 |
| ensg00000186831 |
| picart1 |
| nt5dc2 |
| ttr |
| tnnc2 |
| cfap157 |
| ube2q2p13 |
| ensg00000256196 |
| nos1 |
| loc613038 |
| cd36 |
| ensg00000286550 |
| sorcs3 |
| galr1 |
| linc02033 |
| abcc6p1 |
| nts |
| pitx1-as1 |

| |
|---|
| ensg00000230537 |
| ifi27 |
| ensg00000279196 |
| sostdc1 |
| ndst4 |
| linc00958 |
| ensg00000280179 |
| ensg00000213089 |
| myl7 |
| spag6 |
| ensg00000288818 |
| tincr |
| linc02348 |
| hpd |
| loc105376654 |
| sntg2 |
| ensg00000289305 |
| loc124906789 |
| rmst |
| prss3 |
| casp12 |
| neto1 |
| loc101927245 |
| prok1 |
| onecut2 |
| nrg3-as1 |
| ensg00000271789 |
| ensg00000282870 |
| nptx1 |
| samd11 |
| linc01933 |
| nhlrc4 |
| zmat4 |
| kcns2 |
| sorcs1 |
| tekt1 |
| prame |
| ensg00000228852 |
| hmcn2 |

| |
|---|
| kcnk9 |
| tmem215 |
| kcnj3 |
| ensg00000253227 |
| ensg00000204791 |
| ensg00000233117 |
| ensg00000266473 |
| prlr |
| ensg00000278966 |
| ism2 |
| ensg00000272267 |
| dach1 |
| mycnos |
| cdh9 |
| ensg00000288721 |
| arrdc5 |
| tacr3 |
| sp6 |
| linc01586 |
| dnah6 |
| ensg00000275202 |
| mybl2 |
| ensg00000230882 |
| cdh22 |
| ensg00000279673 |
| kcnv1 |
| kiaa2012 |
| crhr2 |
| ensg00000230804 |
| ensg00000279601 |
| ensg00000176268 |
| neurod2 |
| ensg00000251495 |
| agbl1 |
| ensg00000207294 |
| slc5a4 |
| pcdh19 |
| hadhap1 |
| ensg00000265282 |

| |
|---|
| foxd4l6 |
| nxph4 |
| nell1 |
| necab2 |
| slc35g5 |
| zbtb32 |
| ensg00000279900 |
| npw |
| linc02192 |
| baiap3 |
| ensg00000259869 |
| sco2 |
| col2a1 |
| sctr |
| ensg00000273113 |
| ankrd26p3 |
| cbln4 |
| ighm |
| casr |
| ensg00000270557 |
| tph2 |
| ensg00000285873 |
| ensg00000271857 |
| ensg00000263326 |
| rpl7ap39 |
| linc01305 |
| dclk3 |
| ensg00000275613 |
| ensg00000286044 |
| thsd7b |
| rtp1 |
| neurod1 |
| krtap5-9 |
| npy2r |
| ror2 |
| ensg00000279057 |
| linc00642 |
| ensg00000272971 |
| ensg00000289225 |

| |
|---|
| loc105373390 |
| ptprt |
| npffr2 |
| ensg00000260487 |
| cyp4f26p |
| ensg00000280053 |
| wnt2 |
| acta2-as1 |
| p2ry1 |
| cyp26a1 |
| ndst3 |
| loc124906712 |
| linc02182 |
| rn7sl417p |
| ensg00000291062 |
| gpc3 |
| ensg00000281100 |
| ensg00000226747 |
| loc728307 |
| wnt6 |
| otx2-as1 |
| linc00606 |
| ensg00000277156 |
| ensg00000272386 |
| oprm1 |
| meis2 |
| cacna1i |
| kcnq5 |
| ensg00000259940 |
| hcrtr1 |
| efhd2-as1 |
| ct45a1 |
| klk5 |
| rn7sl608p |
| linc02249 |
| ensg00000290918 |
| ensg00000276863 |
| tex26 |
| ensg00000285865 |

| |
|---|
| ensg00000260410 |
| linc01201 |
| pou2f1-dt |
| ct45a3 |
| evpl |
| ensg00000188828 |
| linc02574 |
| ensg00000272321 |
| tac3 |
| ensg00000260743 |
| ensg00000272103 |
| ptgfr |
| ensg00000278012 |
| rtp5 |
| tuba3d |
| ensg00000267108 |
| sertm2 |
| kcnk3 |
| drd1 |
| enc1 |
| ensg00000272942 |
| slc17a6 |
| ensg00000237422 |
| glra2 |
| ensg00000287925 |
| ensg00000228444 |
| vwa5b1 |
| ensg00000260011 |
| ensg00000275178 |
| ensg00000288770 |
| ensg00000259031 |
| ensg00000269889 |
| c20orf203 |
| pth2r |
| linc00494 |
| ensg00000225398 |
| ensg00000279821 |
| c7orf57 |
| prr33 |

| |
|---|
| cfap58 |
| ensg00000249937 |
| pitx2 |
| ensg00000267986 |
| ensg00000271105 |
| pnma5 |
| npr 3.00 |
| tmem275 |
| prss3p4 |
| lhx2 |
| veph1 |
| linc01572 |
| ensg00000288271 |
| ensg00000269954 |
| ensg00000287792 |
| tpsd1 |
| fibcd1 |
| ensg00000254528 |
| linc00269 |
| ensg00000267466 |
| sema5b |
| ensg00000250053 |
| loc101929552 |
| rdm1p2 |
| ensg00000288808 |
| nxf2 |
| ensg00000287720 |
| ensg00000227256 |
| gpr176-dt |
| neurod6 |
| chat |
| ensg00000223298 |
| loc101927702 |
| mrgpre |
| susd2 |
| ensg00000228950 |
| mylk3 |
| ensg00000223914 |
| ptprt-dt |

| |
|---|
| ensg00000287617 |
| tnf |
| tfap2a-as2 |
| diras3 |
| linc00528 |
| nusap1 |
| ensg00000275769 |
| lrrc71 |
| adra1d |
| ensg00000287751 |
| htr1d |
| slear |
| linc02289 |
| morc1 |
| ensg00000290427 |
| gda |
| ensg00000280073 |
| tacr1 |
| neto1-dt |
| fev |
| linc01470 |
| e2f2 |
| pou2f2-as2 |
| ensg00000235020 |
| coch |
| c2cd4a |
| prlhr |
| slc30a8 |
| ensg00000289123 |
| prtn3 |
| drgx |
| ensg00000287209 |
| atp2c2-as1 |
| art5 |
| ensg00000286527 |
| ensg00000283511 |
| cilp |
| otof |
| linc00487 |

| |
|---|
| ensg00000230333 |
| fgf10 |
| tmem233 |
| ensg00000288090 |
| tnnc1 |
| meis1-as2 |
| kcnk17 |
| slc22a1 |
| linc01798 |
| pdgfd |
| ngb |
| adcyap1 |
| lhx9 |
| rprm |
| ensg00000240793 |
| ensg00000288990 |
| loc100133077 |
| adad2 |
| dutp6 |
| cpa4 |
| ensg00000258647 |
| shox2 |
| shank2-as1 |
| ensg00000288858 |
| egfl6 |
| linc02343 |
| fam163a |
| or51e1 |
| nwd2 |
| scgb3a1 |
| ensg00000265413 |
| gsc2 |
| ebf2 |
| myh6 |
| pwwp4 |
| gabre |
| ccno |
| ensg00000248794 |
| cdc42bpg |

| |
|---|
| ensg00000272081 |
| adra2a |
| calcb |
| fgl1 |
| otp |
| penk |
| ensg00000265179 |
| ensg00000204584 |
| pter |
| l1td1 |
| chodl |
| rfpl3s |
| lpar3 |
| ensg00000124549 |
| mab21l2 |
| ensg00000259977 |
| ensg00000260891 |
| mme |
| cyp24a1 |
| c1ql4 |
| ensg00000180438 |
| rprml |
| ensg00000280604 |
| ensg00000237886 |
| ct45a9 |
| rps17p16 |
| ensg00000285535 |
| lrrc53 |
| ensg00000287064 |
| umodl1-as1 |
| klk7 |
| c6orf141 |
| adgrd2 |
| syt10 |
| fgf14-it1 |
| linc02642 |
| cntfr-as1 |
| dgkk |
| dusp13b |

| |
|---|
| dnah3 |
| ankrd45 |
| ezhip |
| irx4 |
| ensg00000276603 |
| kctd19 |
| capsl |
| skor2 |
| trpa1 |
| ensg00000249741 |
| ct45a5 |
| six2 |
| slc7a3 |
| ensg00000279845 |
| crh |
| ensg00000267506 |
| tmc3 |
| lrrc38 |
| ensg00000205424 |
| gfra4 |
| linc01659 |
| ensg00000257681 |
| chrna3 |
| loc401442 |
| vsx2 |
| alx4 |
| ensg00000291225 |
| c1ql3 |
| ensg00000285696 |
| loc124904403 |
| ensg00000275678 |
| evx2 |
| ensg00000287827 |
| cibar2 |
| gabrq |
| loc101928499 |
| linc00290 |
| frem3 |
| ensg00000251535 |

| |
|---|
| dydc2 |
| slc6a4 |
| kcnj4 |
| tlx3 |
| myb |
| sec16b |
| ensg00000261886 |
| qrfpr |
| ensg00000278986 |
| ttll10 |
| slc5a7 |
| ensg00000280285 |
| ensg00000234891 |
| arhgap36 |
| evx1 |
| mir149 |
| ensg00000169662 |
| cartpt |
| uts2 |
| ct45a6 |
| ct45a7 |
| ensg00000257509 |
| ensg00000274080 |
| ensg00000203335 |
| lpa |
| ptf1a |
| cga |
| ensg00000276923 |
| slc9c2 |
| upk3a |
| ct45a10 |
| ensg00000248528 |
| tfap2b |
| ino80b |
| ensg00000241720 |
| ensg00000280089 |
| ensg00000286407 |
| ensg00000236858 |
| dnaaf3 |

| |
|---|
| slc18a3 |
| tmem114 |
| defa3 |
| dbh |
| ensg00000265018 |
| slc6a2 |
| cfap73 |
| ensg00000255372 |
| ensg00000286812 |
| ensg00000270462 |
| ct45a8 |
| linc01229 |
| ensg00000239377 |
| adamts16-dt |
| cfap107 |
| linc00682 |
| ensg00000254270 |
| cdhr4 |
| alpk2 |
| cfap299 |
| ensg00000287922 |
| slc35d3 |

**Supplementary table 21 : RNAseq comparison of human LC/TGNT vs TZ treatment in rats (MALE)**

#### Upregulated (3 genes)

| Pathway Cluster | Genes | AD Relevance in LC |
|---|---|---|
| **Innate Immune / Cytokine Signaling** | IL21R | Upregulation may heighten local inflammatory tone, which can support microglial activation and enhanced clearance of debris, potentially partially compensating for early LC vulnerability. |
| **Toll-Like Receptor (TLR) / Antiviral Innate Immunity** | TICAM2 | Increased TICAM2 can strengthen TLR-driven immune signaling, improving LC-associated immune surveillance and response to AD-related danger signals. |
| **cGMP / Nitric Oxide Signaling** | GUCY1A1 | Higher GUCY1A1 can boost cGMP-mediated vasodilation and neurovascular coupling, improving |

| | | |
|---|---|---|
| | | perfusion and metabolic support to stressed LC neurons in early AD. |

#### Downregulated (18 genes)

| Pathway / System | Genes | AD Relevance in LC |
|---|---|---|
| **Neuropeptide / GPCR Signaling** | PENK, TACR3, TAC3, NPY2R, GAL, GALR1, ADCYAP1 | Dysregulated neuropeptides over activate LC; downregulation may reduce excitotoxic stress. |
| **Adrenergic Signaling** | ADRA1B, ADRA1D | Overactive noradrenaline contributes to LC degeneration; downregulation may protect neurons. |
| **Glutamatergic / Excitatory Transport** | SLC17A6, GRM4 | Excess excitatory signaling stresses LC; downregulation may preserve function. |
| **Inhibitory Neurotransmission** | SLC6A5, SLC6A7 | Impaired inhibition destabilizes LC firing; downregulation may normalize activity. |
| **Somatostatin / Neuromodulatory Signaling** | SSTR2, SSTR3 | Loss of inhibitory modulation increases LC vulnerability; downregulation may reduce stress. |
| **Cannabinoid / Serotonin Signaling** | CNR1, HTR2C | Dysregulated neuromodulation worsens LC stress; downregulation may restore balance. |
| **Orexin / Wakefulness Signaling** | HCRTR1 | Hyperactive orexin signaling stresses LC; downregulation may improve neuron survival. |

**Supplementary table 22 : RNAseq comparison of human LC/TGNT vs TZ treatment in rats (FEMALE)**

#### Upregulated (11 genes)

| Pathway / System | Genes | AD Relevance in LC |
|---|---|---|
| **Neurotrophic / Survival Signaling** | NTRK1, PDGFD | Supports LC neuron survival and resilience; upregulation may protect against degeneration. |
| **Calcium / Excitability Regulation** | CAMK2A, CACNG3, ATP2A1, ATP2A3 | Maintains calcium homeostasis and neuronal firing; upregulation may prevent LC hyperexcitability and degeneration. |
| **Adrenergic Signaling** | ADRA1B | Enhances adaptive noradrenergic signaling; upregulation may improve LC-mediated cognitive support. |

| Source of Variation | % of total variation | P value | P value summary | Significant? | |
|---|---|---|---|---|---|
| Interaction | 0.3112 | 0.7854 | ns | No | |
| Sex | 12.00 | 0.1027 | ns | No | |
| Treatment | 9.124 | 0.1513 | ns | No | |
| ANOVA table | SS (Type III) | DF | MS | F (DFn, DFd) | P value |
| Interaction | 0.004233 | 1 | 0.004233 | F (1, 19) = 0.07626 | P=0.7854 |
| Sex | 0.1631 | 1 | 0.1631 | F (1, 19) = 2.939 | P=0.1027 |
| Treatment | 0.1241 | 1 | 0.1241 | F (1, 19) = 2.236 | P=0.1513 |
| Residual | 1.055 | 19 | 0.05551 | | |

| | | |
|---|---|---|
| **Cardiac / Structural Myosin Genes** | MYH6, MYH7 | May indirectly support metabolic/energy balance in neurons; upregulation could maintain LC cellular health. |
| **cGMP / NO Signaling** | GUCY1A1 | Promotes neurovascular and synaptic signaling; upregulation may improve LC function and resilience. |
| **Iron / Metabolic Homeostasis** | TFRC | Maintains iron balance and supports mitochondrial function; upregulation may protect LC neurons from metabolic stress. |

### Downregulated (9 genes)

| Pathway Cluster | Genes | AD Relevance in LC |
|---|---|---|
| **Excitatory/Arousal Circuit Signaling** | CHRNA7, CHRM5, PDYN, CNR1, SLC18A1, NOS1 | Reducing cholinergic receptor activity may protect LC neurons from excessive depolarization and calcium influx triggered by Aβ. |
| **Wnt / Neuroplasticity Signaling** | WNT4 | Lower Wnt activity may prevent maladaptive plasticity and structural stress responses in LC neurons during AD progression. |
| **Neurodegeneration / APP Processing** | PSEN2, HAP1 | Reduced PSEN2 could limit local Aβ generation; lower HAP1 may decrease vesicle trafficking burden, together reducing cellular stress in LC neurons. |

**Supplementary table 23: 2-way ANOVA of Abeta % area of Hilar**

#### Supplementary table 24: 2-way ANOVA of Abeta % area of DG

#### Supplementary table 25: 2-way ANOVA of Abeta % area of CA3

| Source of Variation | % of total variation | P value | P value summary | Significant? | |
|---|---|---|---|---|---|
| Interaction | 9.707 | 0.1431 | ns | No | |
| Sex | 12.17 | 0.1035 | ns | No | |
| Treatment | 0.006014 | 0.9701 | ns | No | |
| ANOVA table | SS (Type III) | DF | MS | F (DFn, DFd) | P value |
| Interaction | 0.1151 | 1 | 0.1151 | F (1, 19) = 2.333 | P=0.1431 |
| Sex | 0.1444 | 1 | 0.1444 | F (1, 19) = 2.925 | P=0.1035 |
| Treatment | 7.134e-005 | 1 | 7.134e-005 | F (1, 19) = 0.001445 | P=0.9701 |
| Residual | 0.9378 | 19 | 0.04936 | | |

| Source of Variation | % of total variation | P value | P value summary | Significant? | |
|---|---|---|---|---|---|
| Interaction | 0.6033 | 0.7054 | ns | No | |
| Sex | 6.378 | 0.2272 | ns | No | |
| Treatment | 14.33 | 0.0769 | ns | No | |
| ANOVA table | SS (Type III) | DF | MS | F (DFn, DFd) | P value |
| Interaction | 0.01237 | 1 | 0.01237 | F (1, 19) = 0.1473 | P=0.7054 |
| Sex | 0.1308 | 1 | 0.1308 | F (1, 19) = 1.557 | P=0.2272 |
| Treatment | 0.2939 | 1 | 0.2939 | F (1, 19) = 3.499 | P=0.0769 |
| Residual | 1.596 | 19 | 0.08400 | | |

#### Supplementary table 26: 2-way ANOVA of AT8 (pTAU) counts/nm$^2$ of CA1

| Source of Variation | % of total variation | P value | P value summary | Significant? | |
|---|---|---|---|---|---|
| Interaction | 9.002 | 0.1113 | ns | No | |
| Genotype | 0.3886 | 0.7350 | ns | No | |
| Treatment | 0.7985 | 0.6279 | ns | No | |
| ANOVA table | SS (Type III) | DF | MS | F (DFn, DFd) | P value |
| Interaction | 365068 | 1 | 365068 | F (1, 27) = 2.710 | P=0.1113 |
| Sex | 15759 | 1 | 15759 | F (1, 27) = 0.1170 | P=0.7350 |
| Treatment | 32383 | 1 | 32383 | F (1, 27) = 0.2404 | P=0.6279 |
| Residual | 3636753 | 27 | 134695 | | |

**Supplementary table 27: 2-way ANOVA of AT8 (pTAU) counts/nm$^2$ of CA3**

| Source of Variation | % of total variation | P value | P value summary | Significant? | |
|---|---|---|---|---|---|
| Interaction | 7.653 | 0.1033 | ns | No | |
| Sex | 10.46 | 0.0591 | ns | No | |
| Treatment | 9.070 | 0.0774 | ns | No | |
| ANOVA table | SS (Type III) | DF | MS | F (DFn, DFd) | P value |
| Interaction | 332554 | 1 | 332554 | F (1, 27) = 2.844 | P=0.1033 |
| Sex | 454309 | 1 | 454309 | F (1, 27) = 3.885 | P=0.0591 |
| Treatment | 394108 | 1 | 394108 | F (1, 27) = 3.370 | P=0.0774 |
| Residual | 3157566 | 27 | 116947 | | |

**Supplementary table 28: 2-way ANOVA of AT8 (pTAU) counts/nm$^2$ of SB**

| Source of Variation | % of total variation | P value | P value summary | Significant? | |
|---|---|---|---|---|---|
| Interaction | 18.59 | 0.0918 | ns | Yes | |
| Sex | 7.447 | 0.0988 | ns | No | |
| Treatment | 6.350 | 0.1260 | ns | No | |
| ANOVA table | SS (Type III) | DF | MS | F (DFn, DFd) | P value |
| Interaction | 1178730 | 1 | 1178730 | F (1, 27) = 2.298 | P=0.0918 |
| Sex | 472213 | 1 | 472213 | F (1, 27) = 2.924 | P=0.0988 |
| Treatment | 402680 | 1 | 402680 | F (1, 27) = 2.493 | P=0.1260 |
| Residual | 4360810 | 27 | 161511 | | |

**Supplementary table 29: 2-way ANOVA of AT8 (pTAU) counts/nm$^2$ of Hilar**

| Source of Variation | % of total variation | P value | P value summary | Significant? | |
|---|---|---|---|---|---|
| Interaction | 4.337 | 0.2738 | ns | No | |
| Sex | 1.009 | 0.5944 | ns | No | |
| Treatment | 1.019 | 0.5927 | ns | No | |
| ANOVA table | SS (Type III) | DF | MS | F (DFn, DFd) | P value |
| Interaction | 261994 | 1 | 261994 | F (1, 27) = 1.248 | P=0.2738 |
| Sex | 60966 | 1 | 60966 | F (1, 27) = 0.2904 | P=0.5944 |
| Treatment | 61535 | 1 | 61535 | F (1, 27) = 0.2931 | P=0.5927 |
| Residual | 5668202 | 27 | 209933 | | |

#### Supplementary table 30: 2-way ANOVA of microglia amoeboid/ramified %area of CA3

| Source of Variation | % of total variation | P value | P value summary | Significant? | |
|---|---|---|---|---|---|
| Interaction | 13.73 | 0.0741 | ns | No | |
| Sex | 5.911 | 0.2299 | ns | No | |
| Treatment | 6.265 | 0.2170 | ns | No | |
| ANOVA table | SS (Type III) | DF | MS | F (DFn, DFd) | P value |
| Interaction | 0.1702 | 1 | 0.1702 | F (1, 19) = 3.573 | P=0.0741 |
| Sex | 0.07329 | 1 | 0.07329 | F (1, 19) = 1.539 | P=0.2299 |
| Treatment | 0.07768 | 1 | 0.07768 | F (1, 19) = 1.631 | P=0.2170 |
| Residual | 0.9050 | 19 | 0.04763 | | |

#### Supplementary table 31: 2-way ANOVA of microglia amoeboid/ramified %area of SB

| Source of Variation | % of total variation | P value | P value summary | Significant? | |
|---|---|---|---|---|---|
| Interaction | 7.924 | 0.1928 | ns | No | |
| Sex | 10.82 | 0.1311 | ns | No | |
| Treatment | 0.1465 | 0.8563 | ns | No | |
| ANOVA table | SS (Type III) | DF | MS | F (DFn, DFd) | P value |
| Interaction | 0.03200 | 1 | 0.03200 | F (1, 19) = 1.823 | P=0.1928 |
| Sex | 0.04369 | 1 | 0.04369 | F (1, 19) = 2.489 | P=0.1311 |
| Treatment | 0.0005917 | 1 | 0.0005917 | F (1, 19) = 0.03371 | P=0.8563 |
| Residual | 0.3335 | 19 | 0.01755 | | |

#### Supplementary table 32: 2-way ANOVA of microglia amoeboid/ramified %area of Hilar

| Source of Variation | % of total variation | P value | P value summary | Significant? | |
|---|---|---|---|---|---|
| Interaction | 24.93 | 0.0109 | * | Yes | |
| Sex | 11.96 | 0.0654 | ns | No | |
| Treatment | 4.069 | 0.2683 | ns | No | |
| ANOVA table | SS (Type III) | DF | MS | F (DFn, DFd) | P value |
| Interaction | 0.1391 | 1 | 0.1391 | F (1, 19) = 7.968 | P=0.0109 |
| Sex | 0.06672 | 1 | 0.06672 | F (1, 19) = 3.823 | P=0.0654 |
| Treatment | 0.02270 | 1 | 0.02270 | F (1, 19) = 1.301 | P=0.2683 |

| | | | |
|---|---|---|---|
| Residual | 0.3316 | 19 | 0.01745 |

##### Supplementary table 33: 2-way ANOVA total path/speed of habituation data at 11 months

| Source of Variation | % of total variation | P value | P value summary | Significant? | |
|---|---|---|---|---|---|
| Interaction | 1.555 | 0.3986 | ns | No | |
| Treatment | 0.2523 | 0.7329 | ns | No | |
| Genotype | 3.411 | 0.2133 | ns | No | |
| ANOVA table | SS (Type III) | DF | MS | F (DFn, DFd) | P value |
| Interaction | 0.0001508 | 1 | 0.0001508 | F (1, 44) = 0.7267 | P=0.3986 |
| Treatment | 2.448e-005 | 1 | 2.448e-005 | F (1, 44) = 0.1180 | P=0.7329 |
| Genotype | 0.0003309 | 1 | 0.0003309 | F (1, 44) = 1.594 | P=0.2133 |
| Residual | 0.009132 | 44 | 0.0002075 | | |

##### Supplementary table 34: 2-way ANOVA 24h test data at 11 months

| Source of Variation | % of total variation | P value | P value summary | Significant? | |
|---|---|---|---|---|---|
| Interaction | 1.408 | 0.4367 | ns | No | |
| Treatment | 0.09659 | 0.8380 | ns | No | |
| Genotype | 0.2330 | 0.7510 | ns | No | |
| ANOVA table | SS (Type III) | DF | MS | F (DFn, DFd) | P value |
| Interaction | 11273 | 1 | 11273 | F (1, 43) = 0.6164 | P=0.4367 |
| Treatment | 773.6 | 1 | 773.6 | F (1, 43) = 0.04230 | P=0.8380 |
| Genotype | 1866 | 1 | 1866 | F (1, 43) = 0.1020 | P=0.7510 |
| Residual | 786418 | 43 | 18289 | | |

##### Supplementary table 35: Primary antibodies used in IHC

| **Antibody** | **Target** | **Dilution** | **Species Raised In** | **Manufacturer** |
|---|---|---|---|---|
| Aβ 4G8 | Amyloid beta plaques | 1:1000 | Mouse | BioLegend, Cat#800708 |
| Iba1 | Microglia | 1:500 | Rabbit | Wako, Cat#019-19742 |

| | | | | |
|---|---|---|---|---|
| AT8 | P-tau (Ser202, Thr205) | 1:250 | Mouse | ThermoFisher, Cat#MN1020 |
| NeuN | Neurons | 1:200 | Chicken | Millipore, Cat#ABN91 |
| NET | Norepinephrine transporter | 1:200 | Mouse | Invitrogen, Cat# MA5-24647 |

#### Supplementary table 36: Secondary antibodies used in IHC

| Fluorophore | Reacts With | Raised In | Manufacturer Info |
|---|---|---|---|
| Alexa Fluor 568 | Mouse | Goat | ThermoFisher, Cat#A-11031 |
| Alexa Fluor 488 | Rabbit | Goat | ThermoFisher, Cat#A-11008 |
| Alexa Fluor 488 | Chicken | Goat | ThermoFisher, Cat#11039 |
| Alexa Fluor 568 | Rabbit | Goat | ThermoFisher, Cat#A-11036 |

#### Supplementary table 37: Parameters used in IHC quantification macro

| Stain | Size | SD (start here and adjust accordingly) | Circularity |
|---|---|---|---|
| Abeta 4G8 | 350-infinity | 4 | 0-1 |
| IBA1 | 80-infinity | 1 | Ramified .7-1, Reactive .5-.69,<br>Amoeboid 0-.49 |
| NeuN | 10-5,000 | 2 | 0-1 |
| AT8 | 2-250 | 2 | 0-1 |
| NET | 2-1000 | 2 | 0-1 |

#### Supplementary figure 1: Weight change over time of female rats

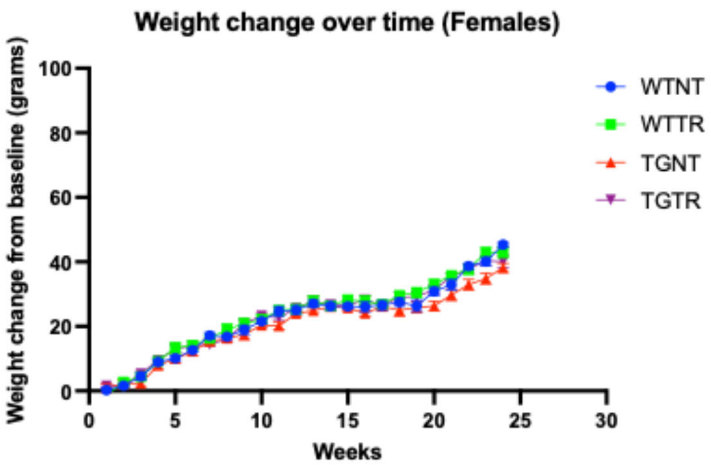

| | | | | |
|---|---|---|---|---|
| Fixed effects (type III) | P value | P value summary | Statistically significant (P < 0.05)? | F (DFn, DFd) |
| Time | <0.0001 | **** | Yes | F (10.55, 200.4) = 527.9 |
| Group | 0.0885 | ns | No | F (3, 19) = 2.523 |
| Time x Group | 0.0059 | ** | Yes | F (31.64, 200.4) = 1.856 |

**Supplementary figure 2: Weight change over time of male rats**

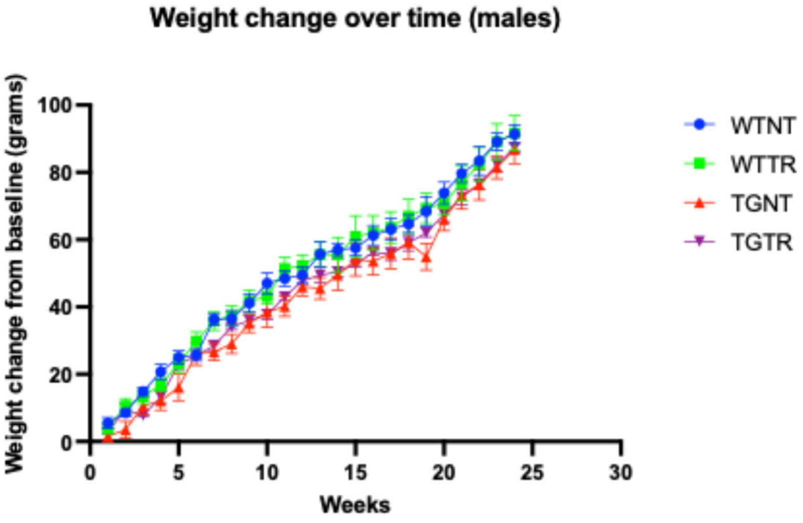

| | | | | | |
|---|---|---|---|---|---|
| Fixed effects (type III) | P value | P value summary | Statistically significant (P < 0.05)? | F (DFn, DFd) | Geisser-Greenhouse's epsilon |
| Time | <0.0001 | **** | Yes | F (6.999, 133.0) = 927.7 | 0.3043 |
| Group | 0.1663 | ns | No | F (3, 19) = 1.886 | |
| Time x Group | 0.2185 | ns | No | F (21.00, 133.0) = 1.254 | 0.3043 |

**Supplementary figure 3: TZ dosage curve over time in females**

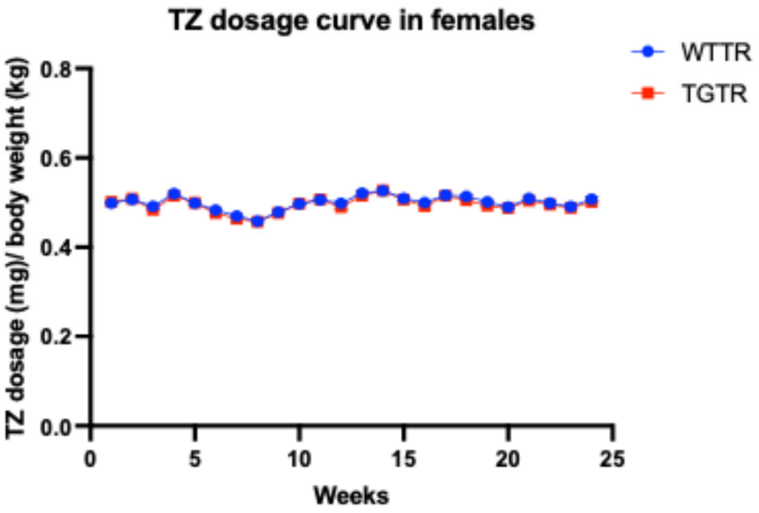

| | | | | | |
|---|---|---|---|---|---|
| Fixed effects (type III) | P value | P value summary | Statistically significant (P < 0.05)? | F (DFn, DFd) | Geisser-Greenhouse's epsilon |
| Time | <0.0001 | **** | Yes | F (5.385, 43.08) = 65.41 | 0.2341 |
| Genotype | 0.4002 | ns | No | F (1, 8) = 0.7896 | |
| Time x Genotype | 0.5671 | ns | No | F (5.385, 43.08) = 0.7953 | 0.2341 |

#### Supplementary figure 4: TZ dosage curve over time in males

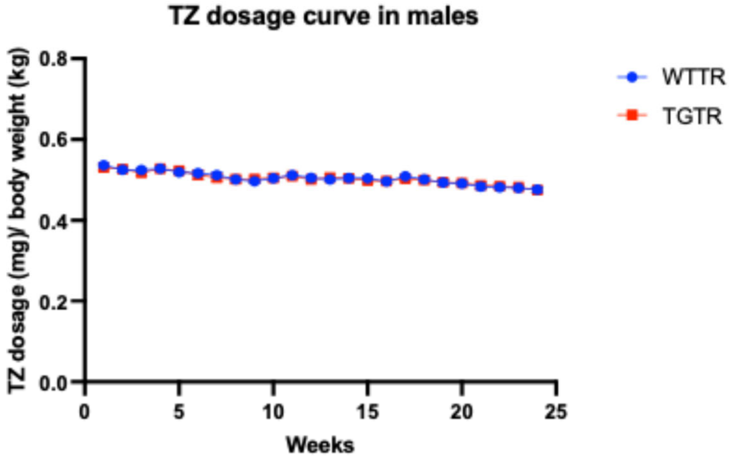

| | | | | | |
|---|---|---|---|---|---|
| Fixed effects (type III) | P value | P value summary | Statistically significant (P < 0.05)? | F (DFn, DFd) | Geisser-Greenhouse's epsilon |
| Time | <0.0001 | **** | Yes | F (6.409, 57.68) = 52.17 | 0.2787 |
| Genotype | 0.9438 | ns | No | F (1, 9) = 0.005254 | |
| Time x Genotype | 0.5668 | ns | No | F (6.409, 57.68) = 0.8186 | 0.2787 |
